# GWAS of flag leaf primary metabolites uncovers metabolic QTL hubs linked to grain yield stability under drought in field-grown *indica* rice

**DOI:** 10.64898/2026.09.28.754946

**Authors:** Chosen Ekene Obih, Zachary Dickens, Giovanni Melandri

## Abstract

Reproductive stage drought disrupts flag leaf metabolism in rice (*Oryza sativa*), limiting photoassimilate translocation to developing grains and reducing grain yield (GY). How natural variation in this response relates to GY remains poorly understood. This study combines untargeted metabolomics of flag leaves from 271 *indica* rice accessions grown under well-watered (WW) and reproductive-stage water-limited (WL) field conditions with genome-wide association studies of 89 annotated polar metabolites. A total of 180 metabolite quantitative trait loci (mQTLs) were identified: 93 detected only under WL, 68 only under WW, and 19 under both conditions, indicating a predominant stress-dependent genetic control; mapping of GY itself detected only four marginal QTLs, none overlapping the mQTLs. Four high-density regions (HDRs) of mQTLs were identified: HDR1-1, detected only under WW conditions and associated mainly with raffinose family oligosaccharides; HDR1-2, HDR4, and HDR11, detected only under WL conditions and associated mainly with tricarboxylic acid cycle intermediates and amino acids. After correcting for genetic relatedness, accessions carrying the favourable haplotype at both HDR1-2 and HDR4 yielded 21.5% more under WL conditions than accessions carrying the unfavourable haplotype at both loci and lost 11.4 percentage points less GY to stress. Accessions carrying the favourable haplotype at only one locus showed a comparable reduction in yield loss. Haplotype-guided variant detection against the 3,000 Rice Genomes Project panel identified major-impact variants in candidate genes, including a polyphenol oxidase at HDR1-2 and tandem cytochrome P450s at HDR4. These hubs are promising, metabolically defined targets for breeding drought-resilient rice.

**Key message:** Combinations of favourable alleles at two metabolic QTL hubs was associated with 21.5% higher grain yield and 11.4 percentage point lower yield loss in *indica* rice, providing promising targets for marker-assisted selection.

## Introduction

Rice (*Oryza sativa*) is the primary caloric staple food source for over half of the global population, with its demand projected to rise substantially by 2050, particularly across Asia, and Sub-Saharan African (Rezvi et al. 2023; Bin Rahman and Zhang 2023). Most of the global rice production (approx. 65%) comes from the *indica* subspecies, cultivated predominantly across Southeast Asia, South Asia and China, where it forms the foundation of food security in tropical and subtropical regions (Bin Rahman and Zhang 2023). Yet rice productivity is threatened by the increasing frequency and severity of drought events (Daryanto et al. 2017). Drought is one of the most damaging abiotic stresses for rice, particularly at the reproductive stage when reduction of panicle fertility and pollen viability results in 50 to 100% grain yield loss in rainfed lowland and upland environments, which account for approximately 45% of global rice production area (Venuprasad et al. 2007; Serraj et al. 2011). Despite decades of breeding effort, progress in improving drought tolerance in rice has been constrained by the highly polygenic architecture of yield, its strong genotype-by-environment interaction, and the physiological complexity of drought responses (Sandhu et al. 2014; Kumar et al. 2014). There is consequently an urgent need for new research approaches to identify genetic loci able to confer higher grain yield stability under drought conditions without compromising grain yield potential under optimal growing conditions.

To overcome this complexity barrier, researchers are increasingly focusing on "intermediate" molecular phenotypes that are closer to gene action than macro-phenotypes such as yield (Razzaq et al. 2019; Fernandez et al. 2021). Metabolite levels, as the downstream products of cellular transcription and protein activity, reflect the immediate physiological state of a plant tissue under the influence of environmental conditions (Großkinsky et al. 2015; Melandri et al. 2022). Metabolite-based genome-wide association (mGWA) mapping has been successfully conducted in multiple plant species, including the model plant *Arabidopsis thaliana* (Chan et al. 2010; Wu et al. 2018; Luzarowska et al. 2023; Naake et al. 2024), and important crops such as tomato (Tohge et al. 2020; Alseekh et al. 2020), rice, maize, and wheat (Chen et al. 2020; Zhu et al. 2024). Owing to the comparatively simple genetic architecture of individual metabolites, these studies have generally resolved metabolite quantitative trait loci (mQTLs) of larger effect sizes than those detected for complex agronomic traits such as grain yield (Chen et al. 2016; Razzaq et al. 2019). This higher genetic tractability makes mQTLs valuable entry points for identifying causal genes contributing to more complex traits.

In rice, the flag leaf acts as the critical metabolic source, contributing approximately 50% of the assimilates required for grain filling (Li et al. 1998; Yoshida, 1981). Regardless of water availability, disruptions to flag leaf metabolism directly impair source-to-sink carbon allocation, ultimately altering both the nutritional quality and quantity of the grains (Hu et al. 2020). Under drought stress, the flag leaf metabolome undergoes substantial reprogramming. The accumulation of amino acids, particularly proline, branched-chain amino acids (BCAAs) and other free amino acids, is one of the most consistent metabolic responses to water deficit in cereals. This response reflects both osmoprotective functions and the catabolic signatures of stress-induced protein degradation and nitrogen remobilization (Hayat et al. 2012; Shim et al. 2023; Rossi and Huang 2025). Concurrently, changes in tricarboxylic acid (TCA) cycle intermediates, such as citric and isocitric acid, reflect the remodelling of central carbon metabolism under stress-induced reductions in photosynthetic activity (Obata and Fernie 2012; Zhang and Fernie 2023). Thus, the capacity to maintain metabolic homeostasis under stress is a major determinant of yield stability. In our previous work, we showed that metabolite-based models accurately predicted grain yield (GY) and GY loss in rice under drought conditions, outperforming genomic-based models and further underscoring the functional relevance of metabolic traits for drought adaptation (Melandri et al. 2020a; Melandri et al. 2022). Whether the natural genetic variation governing these drought-induced metabolic shifts is directly associated with GY performance, however, remains an open question.

A key limitation of GWA mapping in crop diversity panels genotyped at low marker density (e.g., genotyping-by-sequencing or low-density arrays) is the difficulty of narrowing QTL intervals, which hinders the identification of causal variants. Advances in sequencing have progressively eased this constraint: first next-generation sequencing (NGS) dramatically increased throughput and lowered cost, and current third-generation single-molecule technologies (e.g., PacBio and Oxford Nanopore) have further improved read length, throughput and accuracy in genomics (Amarasinghe et al. 2020; Marx 2023; Melandri et al. 2025). In rice, the first crop species to be sequenced, these advances have generated extensive genomic resources since the early adoption of NGS (International Rice Genome Sequencing Project and Sasaki 2005), culminating in large-scale resequencing efforts. The 3,000 Rice Genomes Project (3K RGP; (Wang et al. 2018b) provides a dense reference of millions of single nucleotide polymorphisms (SNPs), Insertions and Deletions (INDELs), and structural variants across a globally diverse panel of rice accessions. This kind of resource offers the opportunity to increase genetic resolution and detect variants at QTL regions by leveraging reference panel-based haplotypes, an approach conceptually similar to genotype imputation strategies that have already used the 3K RGP as a reference panel to increase marker density in other rice association panels (Wang et al. 2018a).

In this study, we combined untargeted metabolomics profiling of flag leaves with GWA mapping in a diversity panel of 271 *indica* rice accessions grown under well-watered and reproductive-stage water-limited conditions to dissect the genetic diversity associated with the levels of 89 annotated polar metabolites. This approach enabled the discovery of metabolic QTL hubs whose favourable alleles were associated with higher GY and reduced drought-induced GY loss. Furthermore, candidate genes and candidate causal variants underlying these metabolic QTL hubs were also explored. Pyramiding the favourable alleles at these hubs/loci could help improve GY stability in rice under drought stress.

## Materials and methods

### Genetic resources and field trial

A diversity panel of 271 *indica* rice accessions was grown at the International Rice Research Institute (IRRI), Los Baños, Philippines, during the 2013 dry season, as described by Kadam et al. (2018) and Melandri et al. (2020a) (Fig.1A; Fig. S1A-S1C). Briefly, the trial comprised adjacent well-watered (WW) and water-limited (WL) fields, each containing three replicated blocks of the accessions. Seeds of the 271 accessions were initially sown in a greenhouse and then transplanted in groups based on their days-to-flowering to achieve flowering synchronization during stress imposition. The WW field was maintained under continuous flooding, while the WL field was subjected to 14 consecutive days of water withholding at the reproductive stage (starting at 50% flowering), followed by re-watering to flooded conditions until maturity, when grain yield (GY, g m^-2^) and GY loss (%) were determined.

**Fig 1.**
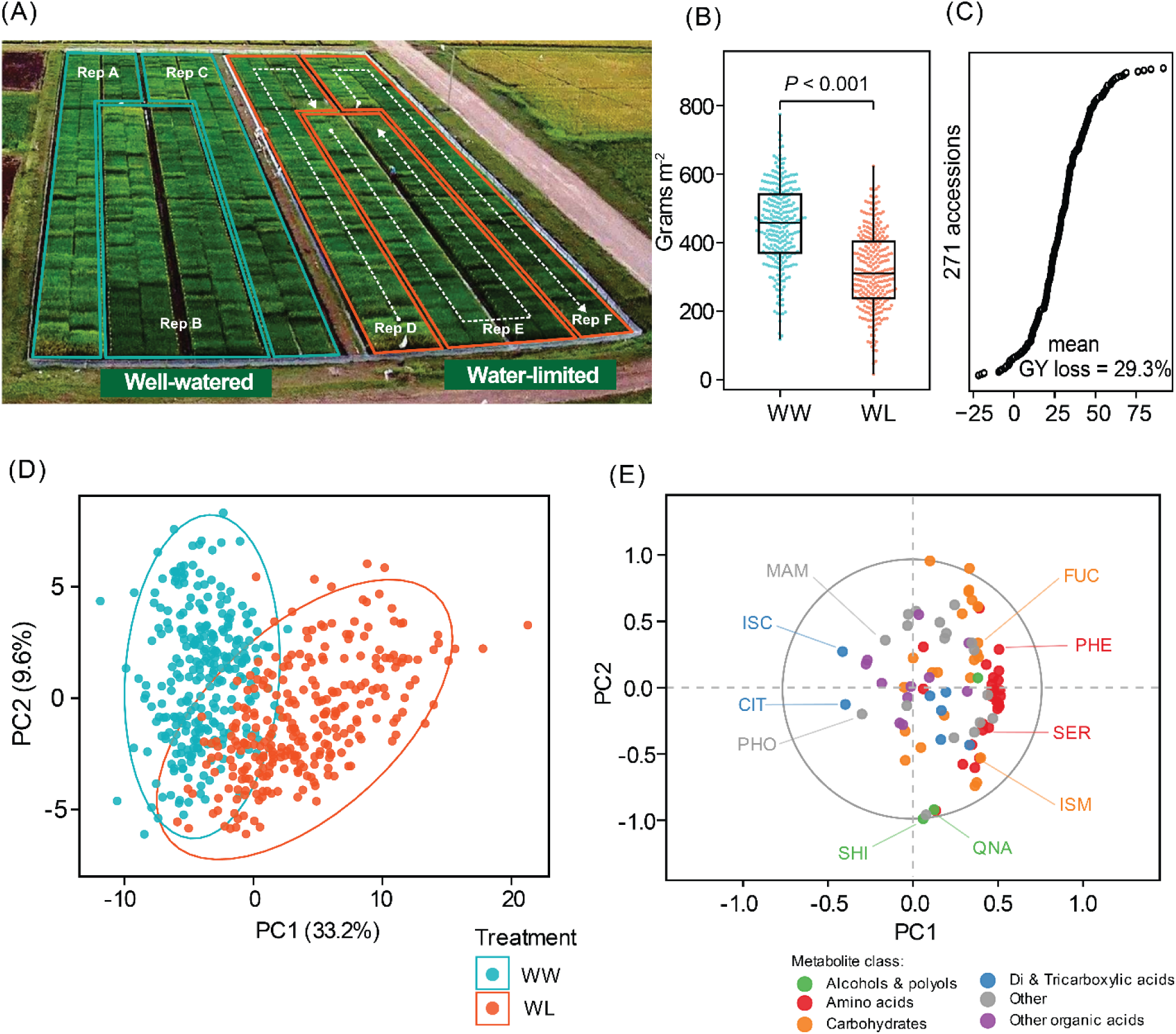
Field experimental design, grain yield response, and flag leaf metabolome shifts under well-watered and water-limited conditions in the 271 *indica* rice accessions. (A) Aerial view of the field trial at the International Rice research Institute (IRRI), Los Baños, Philippines, showing the three replicated blocks of the well-watered (WW, teal outlines; Reps A - C) field and the three replicated blocks of the water-limited (WL, red outlines; Reps D - F) field. (B) Grain yield (GY, g m^-2^) of the 271 accessions under WW (teal) and WL (red) conditions. Box plots show the median and interquartile ranges, overlaid with individual data points; statistical significance was assessed using a Welch’s t-test (*P* < 0.001). (C) The 271 accessions ranked by percentage grain yield loss (GY loss) under WL relative to WW, illustrating the distribution of drought-induced yield loss across the panel. (D) Principal component analysis (PCA) score plot of the 271 accessions based on the 89 polar metabolites quantified by GC-MS under WW (teal) and WL (red) conditions. Ellipses represent the 95% confidence regions for each treatment. (E) Loadings of the 89 metabolites on PC1 and PC2. Metabolites are coloured by chemical class: amino acids (red), carbohydrates (orange), di- and tricarboxylic acids (blue), alcohols and polyols (green), other organic acids (purple) and other compounds (grey).

## Metabolic profiling and chemical taxonomy

On day 14 of water withholding, flag leaves from the WL field were sampled between 09:30 and 11:00 h from the two middle rows of each experimental plot (Fig.1A; Fig. S1B) and snap-frozen in liquid nitrogen; sampling of WW field followed two days later during the same time window. For each accession and condition, tissues from biological replicates were pooled and subjected to untargeted gas chromatography-mass spectrometry (GC-MS)-based metabolomics profiling. In total, 89 polar metabolites were annotated and quantified. Their chemical identifiers (InChIKeys) of these metabolites were used in ClassyFire (Djoumbou Feunang et al. 2016), a web-based application, for classification into six ‘chemical groups’ (alcohols & polyols, amino acids, carbohydrates, di- & tricarboxylic acids, other organic acids, and others) based largely on their chemical subclass (Table S1). Due to their high concentrations in leaf tissues and potential detector saturation for the mass spectrometer, the soluble sugars glucose, fructose, and sucrose were instead quantified spectrophotometrically (Riewe et al. 2008) and used for all subsequent analyses. All metabolite values were log_10_-transformed prior to statistical analyses (Table S2) to improve normality.

## Genome-wide association mapping of metabolite traits

The panel was genotyped with 45,505 SNPs (Data S1) filtered for minor allele frequency (MAF) ≥ 0.05. Further details on the genetic map can be found in Melandri et al. (2020b).

Genome-wide association (GWA) mapping between the SNPs and log_10_-transformed values of each metabolite was performed using a mixed linear model (MLM) in rTASSEL (Monier et al. 2022) incorporating a kinship matrix (centered IBS matrix; Endelman and Jannink 2012) as a random effect. Principal component analysis (PCA) based on the 45,505 SNPs was conducted using the *prcomp* function in R to quantify subpopulation structure (Fig. S2A). Because the first principal component (PC1) explained only 8.3% of the genetic variance and the first two PCs combined failed to clearly separate groups of accessions, no PC covariates were included in the MLM to adjust for subpopulation structure (Fig. S2A; Melandri et al. 2020b).

## Definition of quantitative trait loci (QTLs) and high-density regions of QTLs

A quantitative trait locus (QTL) was defined based on three simultaneous criteria: (i) the presence of at least one SNP exceeding with a -log_10_ (*P*) ≥ 4, with significance tiers defined as high (≥ 5.96, Bonferroni-corrected at α = 0.05), medium (≥ 5) and low (≥ 4); (ii) a QTL interval spanning ±100 kb upstream and downstream of the most significant SNP(s), a window selected based on the estimated genome-wide linkage disequilibrium (LD) decay of the association panel, where the average pairwise LD (*r^2^*) declined to 0.2 at approximately 100 kb (Fig. S2B; Crowell et al. 2016; Zhou et al. 2017); and (iii) the presence of a minimum of three SNPs with a -log_10_ (*P*) ≥ 3.5 within the interval, including the most significant SNP(s) from criterion (i). QTLs were named following the standard nomenclature proposed by McCouch et al. (1997). Overlapping QTLs were defined as those sharing ≥ 50% of their intervals and genomic regions harbouring four or more overlapping QTLs associated with different metabolites were designated as high-density regions (HDRs) of QTLs.

## Genome-wide association mapping of principal components of HDRs’ metabolites

To determine whether the coordinated variation among metabolites associated with each high-density region (HDR) was attributable to a common genetic locus, a principal component analysis (PCA) was performed using the values of metabolites associated with each HDR as variables. The accession scores on the first principal component (PC1), representing the dominant axis of metabolite variation across the panel, were extracted and used as quantitative phenotypes for GWA mapping that was performed using the same MLM framework implemented in rTASSEL as described above. The same significance thresholds and QTL definition criteria applied to single metabolite traits were used to define HDR-related QTLs.

## Linkage disequilibrium structure and haplotype block definition

Pairwise linkage disequilibrium (LD; *r^2^*) was computed across ±500 kb flanking regions of the most significant SNP at each HDR-related QTL using LDBlockShow v1.41 (Dong et al. 2021). To visualize regional association signals alongside the underlying LD structure, local Manhattan plots were generated for SNPs exceeding -log_10_ (*P*) ≥ 4.0. The actual LD block computation used all available SNPs within the ±500 kb interval, irrespective of their significance. LD blocks were defined using the Gabriel method (Gabriel et al. 2002) implemented within LDBlockShow v1.41(Dong et al. 2021).

## Haplotype analysis and association with metabolites and GY

Within the QTL LD blocks of the identified HDRs, SNPs were filtered according to their pairwise LD (*r^2^*) with the most significant SNP, calculated using PLINK v1.9 (Chang et al. 2015). SNPs showing strong LD with the most significant SNP (*r^2^* ≥ 0.6 for HDR1-1, HDR1-2, and HDR11; *r^2^* ≥ 0.8 for HDR4) were retained for haplotype construction. The higher threshold applied to HDR4 reflected its stronger local LD, thereby restricting haplotype construction to variants most tightly linked to the association signal. Haplotypes were constructed using the R package ‘geneHapR’ v1.2.4 (Fig. S3; Zhang et al. 2023), and haplotype groups containing fewer than ten accessions were excluded from downstream analyses. To assess the effects of haplotype variation on metabolite abundance, grain yield (GY) and GY loss, differences among haplotype groups were tested using a kinship-corrected mixed linear model (MLM) to account for genetic relatedness among accessions, which could otherwise confound haplotype-group differences with underlying population structure. The MLM was applied both to the single-locus haplotype groups (HDR1-1, HDR1-2, HDR4 and HDR11 individually) and to multi- HDR haplotype groups defined by concatenating genotype calls at HDR1-2 and HDR4. Multi-HDR haplotype groups representing fewer than ten accessions were excluded. GY was modelled separately for well-watered (WW) and water-limited (WL) conditions considering the haplotype group as a fixed effect and the accession as a random effect, with the random effect’s covariance structured by the same kinship matrix used in GWA mapping; consistent with its limited contribution to the GWA mapping described above, no population-structure principal components were included as covariates. Models were fitted by restricted maximum likelihood using the *mmes* function of the R package ‘sommer’ v4.4.1 (Covarrubias-Pazaran 2016). The significance of the haplotype-group effect was assessed with a joint Wald chi-square (Wald χ^2^) test and pairwise differences among haplotype-group means (expressed as best linear unbiased estimates) were tested using Tukey-Kramer-adjusted contrasts (α = 0.05)

## Identification of candidate variants within HDRs using the 3K Rice Genomes Project reference panel

To increase the resolution for detecting genetic variants underlying the mapped HDR QTLs, a reference panel comprising approximately 29 million SNPs and 2.3 million INDELs from the 3,000 Rice Genomes Project (Wang et al. 2018b) was used for variant discovery. Genotype data for the 1,785 *indica* and *indica*-admixed accessions (ind1A, ind1B, ind2, ind3, and indx) were extracted from the 3K RGP and merged with the 271 *indica* accessions of the GWA mapping panel using the 45,505 SNPs of the genetic map as anchoring markers. Merging was performed using BCFtools v1.19 (Danecek et al. 2021). Following the resolution of allele mismatches and the application of standard quality filters (SNP missingness < 10%, sample missingness < 20%, and MAF ≥ 0.02), the combined dataset was LD-pruned (*r^2^* < 0.3) and subjected to a principal component analysis (PCA) using PLINK v1.9 (Chang et al. 2015). PC1-3 were used to ensure that clustering reflected population structure rather than platform-specific batch effects between the two panels (Fig. S11). To identify the 3K RGP *indica* and *indica*-admixed accessions most genetically similar to the GWA mapping panel, pairwise Euclidean distances were calculated between each of the 1,785 3K RGP *indica* accessions and each of the 271 GWA mapping panel accessions. These distances were computed across the multidimensional space defined by the first 10 PCs, using the *dist* function in R. For each 3K RGP *indica* accession, the mean Euclidean distance across all 271 GWA mapping panel accessions was computed, and accessions were ranked accordingly. The 500 accessions with the smallest mean Euclidean distances to the GWA mapping panel were selected as the reference subset for all subsequent analyses.

## Haplotype-based anchoring, variant extraction and annotation, and haplotype-variant association analysis

For each HDR, the SNPs defining the haplotypes (anchoring SNPs), identified by GWA mapping (6 and 20 SNPs for HDR1-2 and HDR4, respectively) were queried against the 3K RGP variant call format (VCF) file. Using the selected 500 reference *indica* accessions, high-density variants were extracted. Accessions were grouped into haplotypes defined by their multi-genotype strings across the anchoring SNPs, and only haplotype groups containing ≥ 50 accessions were retained as major haplotypes.

SNPs and INDELs within the LD block boundaries of each HDR were extracted from the 3K RGP dataset for the 500 reference accessions. INDELs called against the IRGSP-1.0 reference genome were converted to VCF format, harmonized with the SNP calls, and merged using BCFtools v1.19 (Danecek et al. 2021). Monomorphic sites and variants with > 20% missingness were removed (Data S3 - S4). The merged SNP-INDEL VCF file was annotated using SnpEff v5.2 (Cingolani et al. 2012) with a custom database built from the IRGSP-1.0 genome assembly and gene annotations (Ensembl Plants release 62; https://plants.ensembl.org, archived at https://sep2025- plants.ensembl.org<u>/index.html</u>). Each variant was assigned to a predicted functional effect category: HIGH and MODERATE impact variants (e.g. missense, frameshift, splice site mutations); LOW impact variants (e.g. synonymous); and MODIFIER variants (e.g. intronic and UTR), representing potential regulatory contributors (Data S5). To identify candidate causal variants within each HDR, alternative (ALT) allele frequencies for each variant were computed within each major haplotype group. Variants were classified as ’haplotype-enriched’ (i.e., showing a pronounced difference in allele frequency between haplotype groups) if the maximum pairwise allele frequency difference between any two groups was ≥ 0.5. Variants were classified as ’haplotype-specific’ if the ALT frequency reached ≥ 0.8 in one group while remaining ≤ 0.1 in all others, indicating near-exclusive occurrence within a single haplotype.

## Statistical analysis

All statistical analyses were conducted in R v4.4.3 (R core team, 2025) and data visualizations were produced using the R package ‘ggplot2’ v4.0.3 (Wickham 2016).

## Results

### Water limitation reshapes flag leaf metabolism and its association with grain yield

A panel of 271 *indica* rice accessions from the major rice tropical and subtropical rice-growing regions of the world were grown under well-watered (WW) and water-limited (WL) field conditions to identify the biochemical basis of drought tolerance in rice (Fig. 1A; Fig. S1B). Water limitation, consisting of 14 consecutive days of water withholding at flowering, induced a highly significant reduction (Welch’s *t*-test, *P* < 0.001) in mean grain yield (GY) across the panel (mean reduction of 132 g m^-2^; Fig. 1B). A corresponding overall shift in the flag leaf metabolism of the panel was revealed by a principal component analysis (PCA) of the samples based on all the 89 annotated polar metabolites (Table S1- S2) quantified by gas chromatography-mass spectrometry (GC-MS). The first two principal components explained 42.8% of the total sample variation, with PC1 alone accounting for 33.2% (Table S3). Their combination separated the WW and WL samples, albeit with some overlap (Fig. 1D).

Loadings of the metabolites on PC1 revealed that samples with the highest PC1 scores (rightmost in Fig. 1D; Table S4) were characterized by elevated levels of most free amino acids and, to a lesser extent, carbohydrates. Conversely, WL samples with lower PC1 scores (leftmost, overlapping with WW) were associated with higher levels of the tricarboxylic acid (TCA) cycle intermediates citric and isocitric acid, as well as and other organic acids (Fig. 1E). Interestingly, the PC1 scores of WL samples were significantly and negatively correlated with GY under WL conditions (Spearman’s *r_s_* = -0.51, Bonferroni-corrected *P* < 0.001), whereas no significant correlation was detected between the PC1 scores of WW samples and GY under WW conditions (*r_s_* = -0.19, Bonferroni-corrected *P* = 0.12; Table S5). Among individual metabolites, all amino acids displayed negative correlations with GY under WL conditions (mean *r_s_* = -0.4), while citric and isocitric acid showed the strongest positive correlations (*r_s_* = 0.44 and 0.56, respectively; Table S6). These results indicate that PC1 represents a composite proxy of flag leaf stress metabolism that is predictive of GY performance under WL conditions.

## Genetic basis of natural variation in flag leaf metabolism revealed by genome-wide association mapping

The 271 accessions of the panel were genotyped with 45,505 single nucleotide polymorphic markers (SNPs; Data S1). GY under WW and WL conditions displayed low estimates of SNP-based narrow-sense heritability (*h^2^*) at 0.25 and 0.27, respectively. By contrast, flag leaf metabolites were considerably more heritable: seven metabolites under WW and ten under WL conditions exceeded *h^2^* = 0.4. Overall, 26 of 89 (29.2%) metabolites under WW and 41 (46.1%) under WL exceeded the heritability of their corresponding GY value (Table S7). The higher heritability of metabolites relative to GY, particularly under stress, was mirrored in the GWA mapping results: 112 metabolic QTLs (mQTLs) were detected under WL conditions and 68 mQTLs under WW conditions, whereas only four QTLs were identified for GY (Fig. 2A - 2C; Table S8). Within each treatment, the significance values of the mQTLs were similarly distributed. Low-significance mQTLs (4 ≤ -log_10_ (*P*) < 5) predominated, accounting for 69.1% under WW and 78.6%under WL. These were followed by high-significance mQTLs (-log_10_ (*P*) ≥ 5.96; 14.7% under WW; 13.4% under WL) and medium-significance ones (5 ≤ -log_10_ (*P)* < 5.96; 16.2% under WW; 8% under WL; Fig. 2C). Notably, of the 19 mQTLs detected in both treatments, 12 were of medium- or high-significance (Table S9). Manhattan and Quantile-Quantile (QQ) plots for all 89 metabolite traits and GY mapped under WW and WL conditions are provided in Supplementary Information: Fig. S8A - S96D.All four QTLs mapped for GY were of low-significance (-log_10_ (*P*) = 4.0-4.9), located on chromosomes 4, 8, 9, and 12, and did not colocalize with any mQTLs. Among metabolite classes, most mQTLs were mapped for carbohydrates (46), followed by amino acids (43), other compounds (37), alcohols and polyols (31), di- and tricarboxylic acids (13), and other organic acids (10) (Table S8). The wide distribution of mQTLs across all 12 rice chromosomes indicates a dispersed genetic architecture underlying the total variation in the flag leaf metabolome (Fig. 2A).

**Fig. 2.**
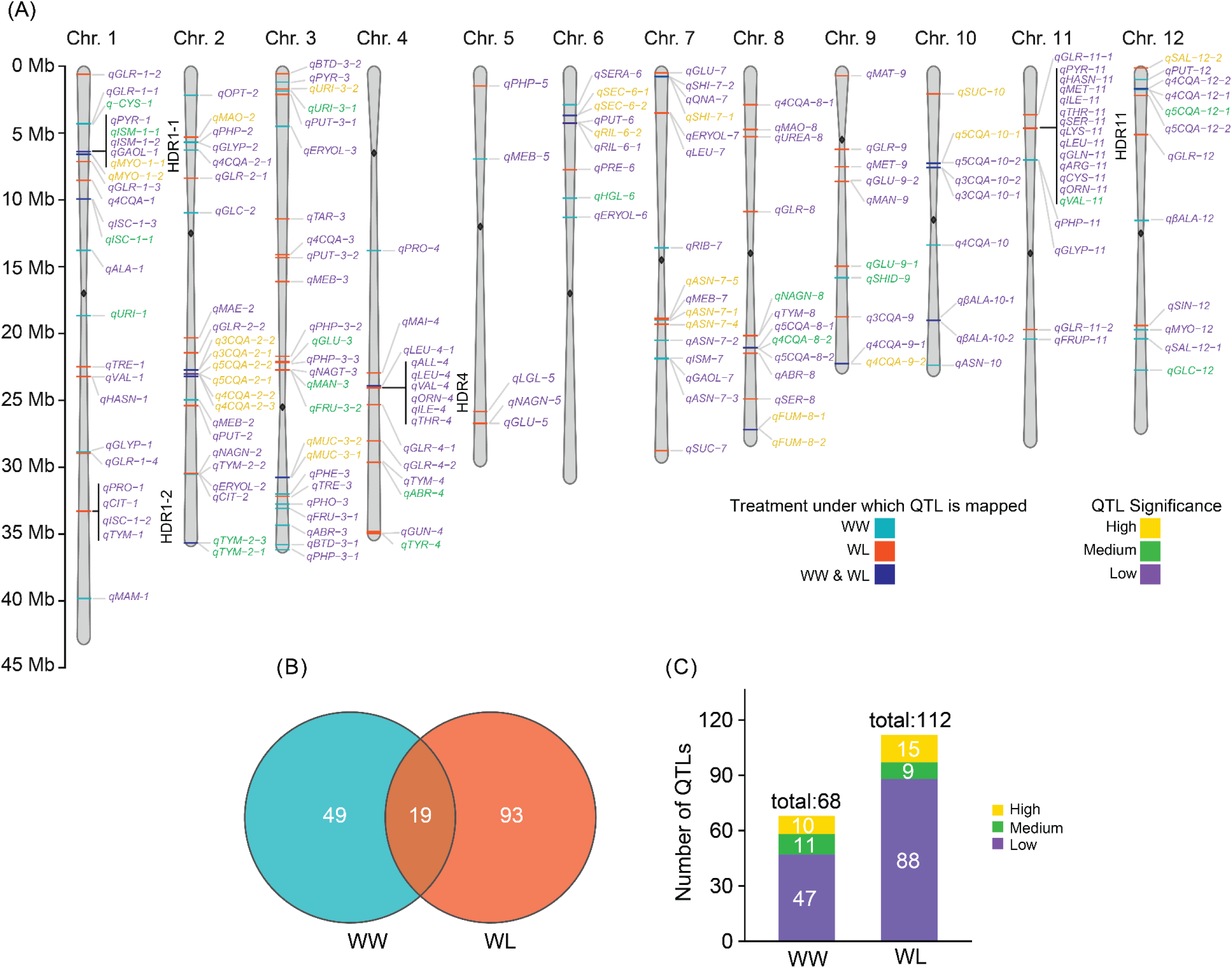
Genome-wide landscape of metabolite QTLs detected in the 271 *indica* rice accessions under well-watered and water-limited conditions. (A) Genome-wide distribution of 180 metabolite QTLs across the 12 rice chromosomes. Vertical bars on each chromosome indicate QTL positions, coloured based on the treatment under which they were detected: WW (teal), WL (red), and both WW and WL (blue). QTL names are colour-coded by significance tier: yellow (high significance, -log_10_ (*P*) ≥ 5.96), green (medium significance, 5 ≤ -log_10_ (*P*) < 5.96) and purple (low significance, 4 ≤ -log_10_ (*P*) < 5). (B) Venn diagram illustrating the overlap between QTLs detected under WL (93 unique QTLs) and WW (49 unique QTLs), with 19 QTLs shared across both treatments. (C) Number of QTLs detected under each treatment, stratified by significance tier (high, medium, and low).

The strongest purely genetic signals were mapped on chromosome 2 (22.63-23.13 Mbp) for three caffeoylquinic acid isomers – 3-O-caffeoylquinic acid (3-CQA); 4-O-caffeoylquinic acid (4-CQA); 5-O-caffeoylquinic acid (5-CQA) – comprising six overlapping mQTLs with -log_10_ (*P)* values ranging from 6.3 to 12.1, detected under both treatments. The two most significant mQTLs (-log_10_ (*P*) = 26.6 and 27.2) were detected for galactaric acid on chromosome 3 (30.66-30.86 Mbp) under WW and WL conditions, respectively, followed by three asparagine mQTLs on chromosome 7 (18.76-19.41 Mbp;-log_10_ (*P*) = 11.7-18.9). For these metabolites, the combination of high-significance and consistent detection of their mQTLs across treatments highlights that strong constitutive genetic differences drove their variation, with the environment exerting only a marginal influence.

## Identification of four mQTL hubs associated with multiple metabolic pathways

To prioritize loci associated with variation in multiple metabolites, we focused our analysis on high-density regions (HDRs) of QTLs, defined as genomic intervals harbouring four or more overlapping mQTLs associated with four or more distinct metabolites. Four HDRs were identified: HDR1-1, which was detected under WW conditions, and three stress-responsive ones (HDR1-2, HDR4, and HDR11), detected exclusively under WL conditions (Fig. 2A).

HDR1-1 (Chr 1: 6.49-6.70 Mbp) comprised four mQTLs detected under WW conditions: galactinol (*qGAOL-1*) and myo-inositol (*qMYO-1-1*), that are precursors shared with the raffinose family oligosaccharide (RFO) biosynthetic pathway, as well as two mQTLs not assigned to a named pathway, isomaltose (*qISM-1-1*) and pyroglutamic acid (*qPYR-1*). The most significant SNPs (msSNPs) of these mQTLs displayed a high minor allele frequency (MAF: 0.46-0.47). The genetic signal for two of them, *qMYO-1-1* (-log_10_ (*P*) = 9.8) and *qISM-1-1* (-log_10_ (*P*) = 5.5), was also detected under WL conditions, albeit with reduced significance (Table S10).

HDR1-2 (Chr 1: 33.19-33.20 Mbp) contained four mQTLs detected exclusively under WL conditions that mapped to two metabolic classes: tricarboxylic acid (TCA) cycle intermediates (isocitric acid, *qISC-1-2*; citric acid, *qCIT-1*) and stress-related amino acids and amino acids derivatives (proline, *qPRO-1*; tyramine, *qTYM-1*). The association signals from these mQTLs were of low-significance (-log_10_ (*P*) = 4.1-4.5) but featured high MAFs (0.41-0.47; Table S10).

HDR4 (Chr 4: 23.95-24.01 Mbp) harboured six WL-exclusive mQTLs of low-significance (-log_10_ (*P*) = 4.3-4.6) and medium-to-high MAFs (0.35-0.39). Three of them were mapped to branched-chain amino acids (BCAAs: leucine, *qLEU-4-2*; valine, *qVAL-4*; isoleucine, *qILE-4*), while the remaining three were associated with three stress-related compounds, threonine, (*qTHR-4)*, ornithine (*qORN-4*) and allantoin (*qALL-4*) (Table S10).

HDR11 (Chr 11: 4.54-4.74 Mbp) represented a dense cluster comprising 13 WL-exclusive mQTLs characterized by a very low MAF (0.06). All 13 were mapped to a diverse array of amino acids, including BCAAs (isoleucine, leucine, valine), sulphur-containing amino acids (methionine, cysteine), and polar amino acids (threonine, lysine, serine, arginine, glutamine, 3-hydroxyasparagine, pyroglutamic acid, and ornithine) (Table S10).

## Principal component-based proxies of metabolic variation in mQTL hubs and their correlation with grain yield

To evaluate the aggregate metabolic variation captured by each mQTL hub and its relationship with GY, region-specific PCAs were performed using only the metabolites that mapped specifically to each respective HDR. The PCA of the four HDRs showed that the first PC (PC1) consistently captured a dominant share of the total specific locus-specific metabolic variation: 53.8% for HDR1-1, 66.5% for HDR1-2, 83.9% for HDR4, and 81.3% for HDR11 (Fig. 3A-D; Table S11). Consequently, PC1 was used as a synthetic metabolic proxy for each mQTL hub.

**Fig. 3.**
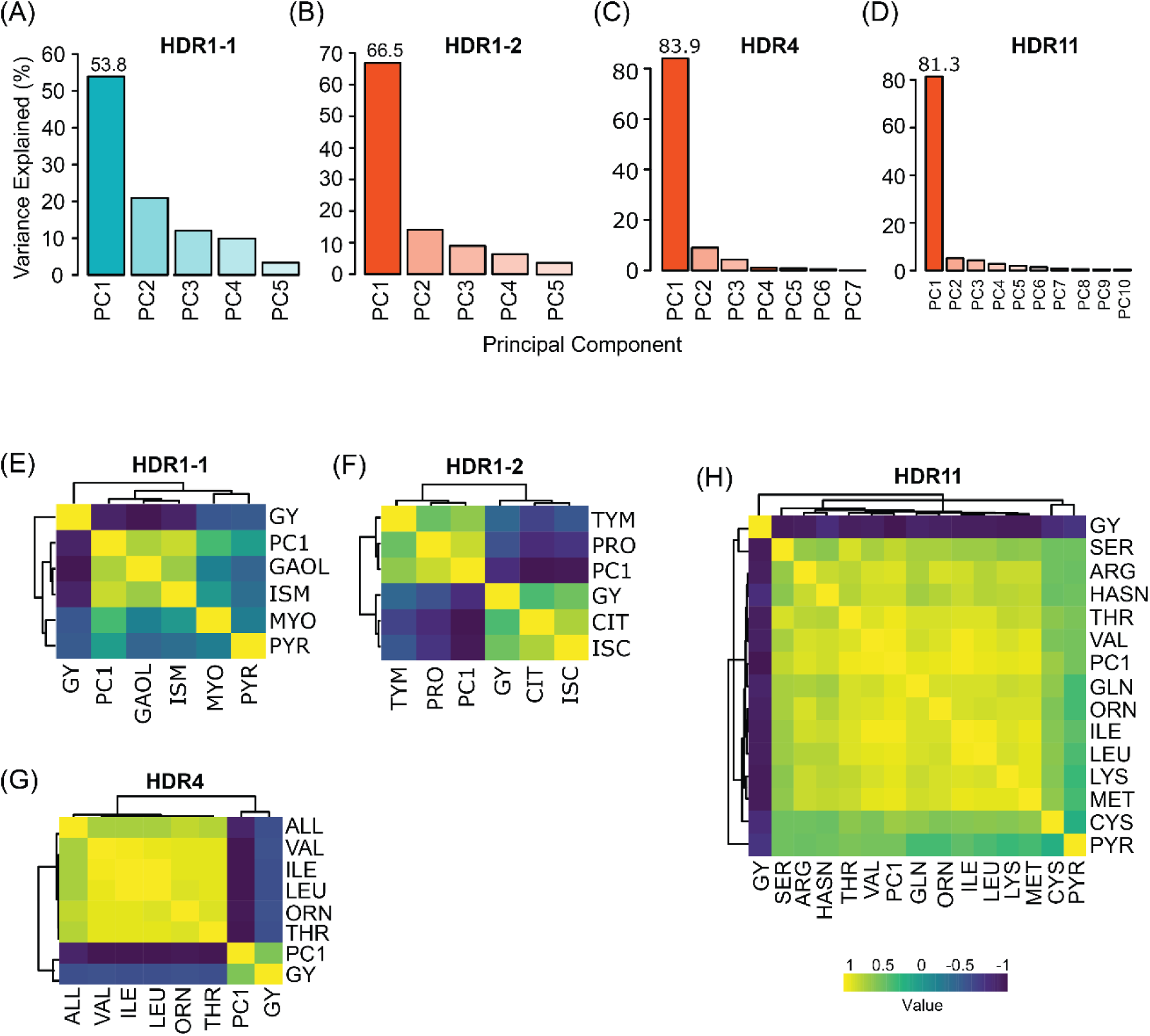
PCA-based metabolic proxies of four high-density regions (HDRs) and their correlation with grain yield. (A-D) Variance explained by the selected principal components of the metabolites mapping to each HDR: (A) HDR1-1 (myo-inositol, galactinol, isomaltose, pyroglutamic acid; PC1 = 53.8%), (B) HDR1-2 (citric acid, isocitric acid, proline, tyramine; PC1 = 66.5%), (C) HDR4 (allantoin, valine, isoleucine, leucine, ornithine, threonine; PC1 = 83.9%) and (D) HDR11 (serine, arginine, 3-hydroxyasparagine, threonine, valine, glutamine, ornithine, isoleucine, leucine, lysine, methionine, cysteine, pyroglutamic acid; PC1 = 81.3%). HDR1-1 values are derived from the well-watered (WW) treatment; HDR1-2, HDR4 and HDR11 values are derived from the water-limited (WL) treatment. (E-H) Spearman correlation heatmaps showing pairwise relationships among individual metabolites, PC1 proxy scores and grain yield (GY) for (E) HDR1-1, (F) HDR1-2, (G) HDR4 and (H) HDR11. The colour scale indicates Spearman *r_s_* values from -1 (dark purple) to 1 (yellow). Dendrograms reflect hierarchical clustering of traits by correlation similarity. Metabolite abbreviations: ALL, allantoin; ARG, arginine; CIT, citric acid; CYS, cysteine; GAOL, galactinol; GLN, glutamine; HASN, 3-hydroxyasparagine; ILE, isoleucine; ISC, isocitric acid; ISM, isomaltose; LEU, leucine; LYS, lysine; MET, methionine; MYO, myo-inositol; ORN, ornithine; PRO, proline; PYR, pyroglutamic acid; SER, serine; THR, threonine; TYM, tyramine; VAL, valine.

Spearman correlation analysis between the PC1 of each HDR and GY revealed a consistent pattern: significant correlations were detected exclusively under WL conditions. Specifically, PC1 of HDR1-1 was weakly and non-significantly correlated with GY under WW conditions (*r_s_* = -0.10, *P* = 0.12). In contrast, the proxies for the three WL- specific hubs were all significantly correlated with GY under WL conditions, showing negatively correlations for HDR1-2 (*r_s_* = -0.51, *P* < 0.001) and HDR11 (*r_s_* = -0.52, *P* < 0.001), and a positive correlation for HDR4 (*r_s_* = 0.52, *P* < 0.001) (Fig. 3E - H; Table S12-S13).

The intra-HDR correlations among metabolites, as well as between metabolites and GY, further elucidated these relationships. HDR4 and HDR11 exhibited strong positive collinearity among their constituent amino acids (*r_s_* > 0.6), consistent with their coordinated accumulation under stress. Notably, all of these amino acids were, in turn, negatively correlated with GY (HDR4: *r_s_* = -0.48 to -0.51; HDR11: *r_s_* = -0.32 to -0.51; all *P* < 0.001), mirroring the typical trend of amino acid accumulation under water limitation (Fig. 3G). In contrast, the metabolites within HDR1-1 maintained weak-to-strong positive correlations with one another (0.15 ≤ *r_s_* ≤ 0.89; Fig. 3E). HDR1-2 displayed a contrasting pattern: the two TCA cycle intermediates, citric and isocitric acid, were strongly positively correlated with each other (*r_s_* = 0.76) and with GY (0.44 ≤ *r_s_* ≤ 0.55) but negatively correlated with the stress-responsive amino acids and derivatives proline and tyramine (*r_s_* < -0.38). This reveals a potential metabolic antagonism within this locus between primary energy metabolism and nitrogen-based stress responses (Fig. 3F).

## PC1-based GWA mapping of mQTL hubs and haplotype analysis

To further investigate the genetic architecture of the four HDRs, we performed GWA mapping using the region-specific PC1s as synthetic phenotypes, followed by linkage disequilibrium (LD) characterization and haplotype analysis.

For HDR1-1, PC1-based GWA mapping identified a QTL on chromosome 1 with three co- localizing msSNPs (Chr1:6,593,333, Chr1:6,593,334 and Chr1:6,593,337; -log_10_ (*P*) = 7.7) (Fig. S4A; Table S14). The local Manhattan plot confirmed that the PC1-based msSNPs co-mapped with the individual mQTLs defining HDR1-1 (*qMYO-1-1, qGAOL-1, qISM-1-1, qPYR-1*). Local LD was examined across a ∼1-Mbp region encompassing ∼500 kbp upstream and downstream of the msSNPs. The pairwise LD estimates (*r^2^*) of the 207 SNPs in this region showed that all three msSNPs mapped to the same 39.14 kbp LD block (Fig. S4B; Table S15). Within this LD block, haplotype variation spanning 36.9 kb and containing 7 SNPs in strong LD (*r^2^* > 0.8) was identified (Fig. S4C), resolving the panel into two main haplotypes: HAP1 (n =144, 53.1%) and HAP2 (n = 116, 42.8%); two additional recombinant haplotypes were detected in the remaining 11 accessions (Fig. S3A). HAP1 carried the alternative alleles across the 7 SNP positions, while HAP2 carried only the reference alleles. After accounting for genetic relatedness of the accessions (by using a kinship-corrected mixed linear model), HAP2 was associated with significantly higher galactinol, isomaltose, and myo-inositol levels (Wald *χ*^2^ > 15.08, *P* < 0.001 for each) under WW conditions, whereas pyroglutamic acid levels did not differ between the two haplotypes (Wald *χ*^2^ = 1.02, *P* = 0.311; Fig. S5A-D). GY did not differ significantly between HAP1 and HAP2 accessions (Wald *χ*^2^ = 0.74, *P* = 0.391; Fig. S4D; Table S17), nor did GY loss (Wald *χ*^2^ = 0.04, *P* = 0.847; Fig. S4E, Table S17).

For HDR1-2, PC1-based GWA mapping identified a QTL with a single msSNP (Chr1:33,603,854; -log_10_ (*P*) = 6.0) (Fig. 4A). Although this SNP did not co-localize with the msSNPs of the individual mQTLs defining HDR1-2 (*qCIT-1, qISC-1-2, qPRO-1*, and *qTYM-1*), it was in close proximity to them (Fig. 4B). The LD block containing the PC1 msSNP spanned 42.4 kbp and enclosed six SNPs (*r^2^* > 0.6; Fig. S5C and Fig. S3B), resolving the panel into three main haplotypes: HAP1 (n = 141, 52%), HAP2 (n = 101, 37%) and HAP3 (n = 26, 9.6%). HAP1 carried only alternative alleles and HAP2 only reference alleles across the six SNPs, while HAP3 was a recombinant haplotype carrying a mixture of alternative and reference alleles. The values of the four metabolites mapped to HDR1-2 differed significantly among the accessions carrying the three haplotypes (Wald *χ*^2^ ≥ 9.17, *P* ≤ 0.034; Fig. S5E-H; Table S16). HAP1 was associated with higher proline and tyramine levels but lower citric and isocitric acid levels than HAP2 and HAP3 (*P_adj_* < 0.03), consistent with the metabolic antagonism at this locus described above (Fig. 3F). For isocitric acid, haplotype significantly affected metabolite abundance, with HAP1 showing lower levels than HAP2 and HAP3 (*P_adj_* ≤ 0.009), while HAP2 and HAP3 did not differ (*P_adj_* = 0.9999; Fig. S5F). Despite differential haplotype-based metabolic profiles, the three haplotypes did not differ significantly in GY under WL conditions (Wald *χ*^2^ = 2.29, *P* = 0.319; all pairwise *P_adj_* ≥ 0.29; Fig. 4D; Table S17). GY loss also did not differ significantly among haplotypes (Wald *χ*^2^ = 4.53, *P* = 0.104; all pairwise *P_adj_* ≥ 0.10; Table S17). Nevertheless, HAP2 showed a numerical trend toward higher GY and lower GY loss than HAP1 and HAP3, with the highest estimated marginal mean for GY (332.6 ± 11.2) and the lowest for GY loss (26.1 ± 2.0), (Table S17).

**Fig. 4.**
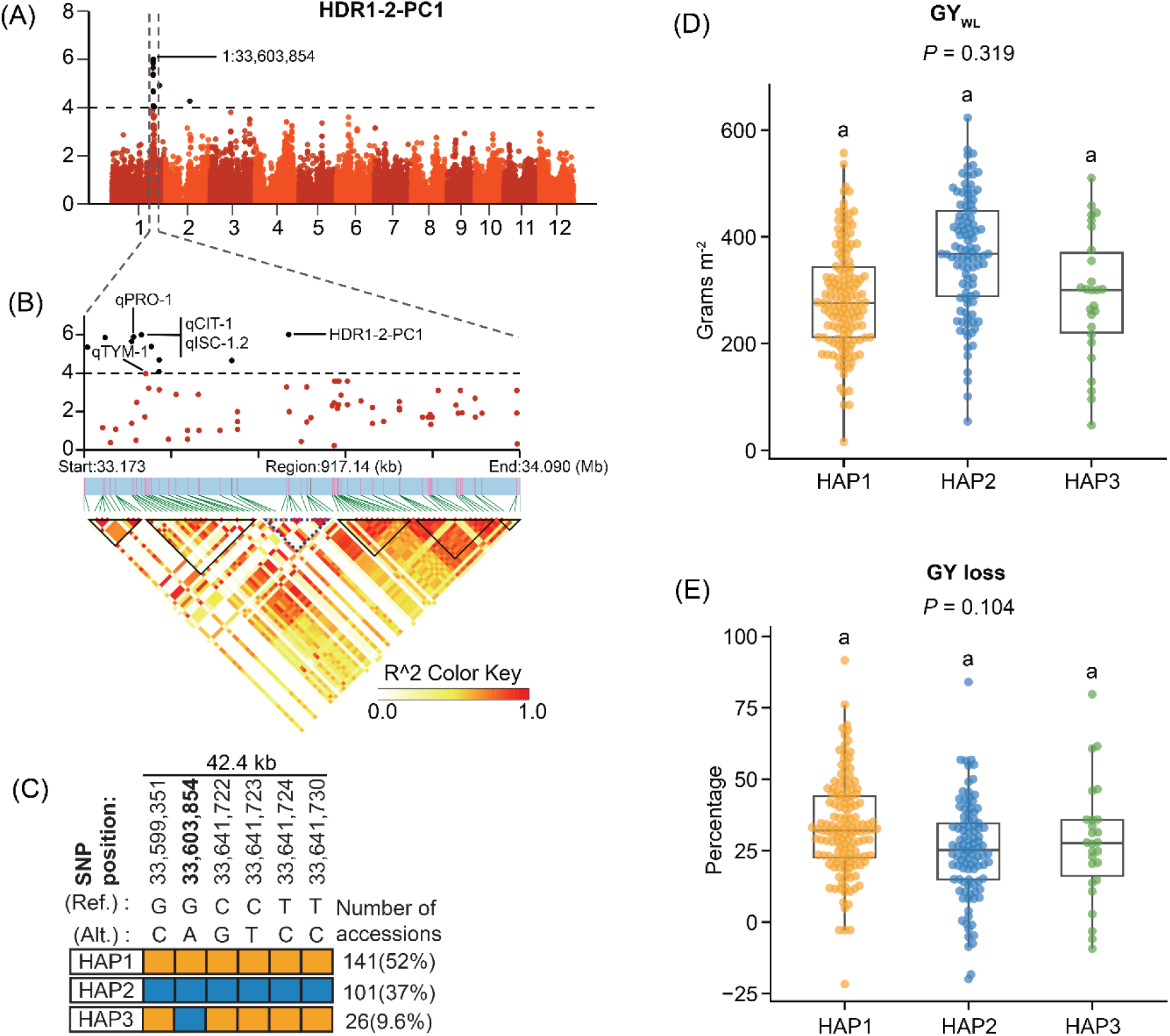
Haplotype analysis, grain yield and grain yield loss associations at HDR1-2 on chromosome 1. (A) Manhattan plot of GWA mapping for the HDR1-2 PC1 metabolic proxy under water-limited (WL) conditions. The most significant SNP (msSNP) on chromosome 1 is indicated. The dashed line represents the significant threshold (-log_10_(*P*) = 4.0). (B) Regional association plot (upper) and pairwise LD heatmap (lower, *r^2^* colour key) for the HDR1-2 local LD interval (Chr 1: 33.173-34.090 Mb). Individual metabolite QTLs lead SNPs are labelled. (C) Haplotype block structure at HDR1-2, comprising 6 SNPs spanning 42.4 kb; the msSNP (33,603,854) is in bold. (D) Grain yield (GY) comparisons between haplotypes. (E) GY loss comparison between haplotypes. Gold, blue and green jitters colours denote alternative, reference and recombinant allelic composition of HAP1, HAP2 and HAP3, respectively. *P-*values are from Wald chi-square test; different letters denote significant differences based on Tukey’s HSD pairwise contrasts (α = 0.05). Box plots show medians, interquartile ranges, and individual data points.

For HDR4, PC1-based GWA mapping identified a QTL with a single msSNP (Chr4:24,053,641; -log_10_ (*P*) = 4.91) that co-localized with the msSNPs of all six mQTLs of the HDR (Fig. 4A-B). The LD block containing the msSNP spanned 98 kbp and enclosed 20 SNPs (Fig. 5C and Fig. S3C), resolving the panel into two major haplotypes: HAP1 (n = 165, 60.9%), carrying only alternative alleles, and HAP2 (n = 91, 33.6%), carrying only reference alleles. HAP2 exhibited significantly lower levels of all six metabolites under WL conditions (Wald *χ*^2^ ≥ 19.80, *P* < 0.001; Fig. S6A-F; Table S16), together with significantly higher GY (Wald *χ*^2^ = 4.27, *P* = 0.039; Fig. 5D; Table S17) and lower GY loss (Wald *χ*^2^ = 5.31, *P* = 0.021; Fig. 5E; Table S17) consistent with the negative correlation between amino acid accumulation and GY at this locus (Fig. 3G).

**Fig. 5.**
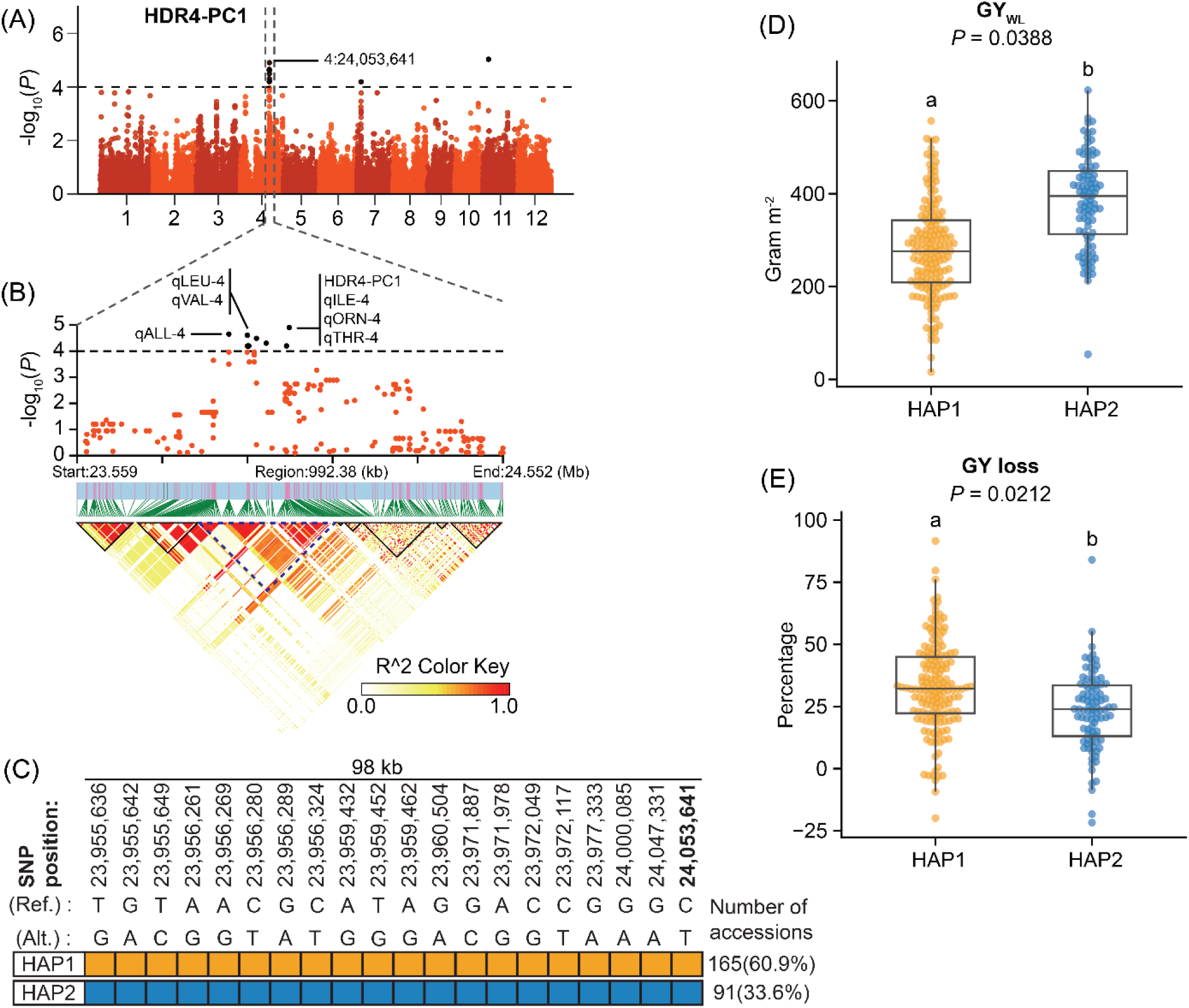
**Haplotype analysis and grain yield associations at HDR4 on chromosomes 4.**(A) Manhattan plot of GWA mapping for the HDR4 PC1 metabolic proxy under water-limited (WL) conditions. The most significant SNP (msSNP) on chromosome 4 is indicated. The dashed line represents the significance threshold (-log_10_(*P*) = 4.0). (B) Regional association plot (upper) and pairwise LD heatmap (lower; *r^2^*) for the HDR4 interval (Chr 4: 23.559-24.552 Mbp). Individual metabolite QTLs msSNPs are labelled. (C) Haplotype block structure at HDR4, comprising 20 SNPs spanning 98 kbp. The msSNP (24,053,641) is indicated in bold. (D) Grain yield (GY) comparisons between haplotypes under water-limited conditions. (E) GY loss comparisons between haplotypes. Reference and alternative alleles are shown; gold and blue jitters denote the alternative and reference allelic composition of HAP1 and HAP2, respectively. *P-*values are from Wald chi-square test; different letters denote significant differences based on Tukey’s HSD pairwise contrasts (α = 0.05). Box plots show medians, interquartile ranges, and individual data points.

For HDR11, three tightly linked msSNPs on chromosome 11 were identified via PC1- based GWA mapping (Chr11:4,640,885, Chr11:4,640,889, and Chr11:4,640,900; -log_10_ (*P*) = 5.04; Fig. S7A), all three of which co-localized with the msSNPs of the individual mQTLs of the hub (Fig. S7B). The LD block containing the PC1 msSNPs enclosed four SNPs spanning only 96 bp (Fig. S7C), resolving the panel into two haplotypes: a common HAP1 (n = 254, 93.7%), carrying only alternative alleles, and a rare HAP2 (n = 13, 4.8%), carrying only reference alleles. All 13 amino acids of this hub displayed significantly higher values in HAP2 than HAP1 (Wald *χ*^2^ ≥ 10.63, *P* ≤ 0.001 for all; Fig. S8A-M), while GY did not differ significantly between haplotypes (Wald χ^2^ = 1.33, *P* = 0.248; Fig. S7D; Table S17), nor did GY loss (Wald χ^2^ = 1.50, *P* = 0.220; Fig. S7E; Table S17).

## Multi-locus haplotype combinations at HDR1-2 and HDR4 reveal a significant effect on grain yield and drought-induced grain yield loss

Although the haplotype-based comparison at HDR1-2 did not reach statistical significance for GY or GY loss under WL conditions, accessions carrying HAP2 (the reference-like haplotype at this locus) exhibited a higher estimated marginal mean (emmean) GY (332.6 g m^-2^) and a lower emmean GY loss (26.1%) than those carrying HAP1 (32.2%) and HAP3 (27.7%) accessions. Because HDR1-2 and HDR4 showed a similar trend – the reference-like allele was associated with higher GY and lower GY loss at both loci – we evaluated the combined effect of these two stress-responsive HDRs. HDR1-1 was not included because it is WW-specific, and HDR11 because its reference allele was too rare in the panel (MAF = 0.06) to form groups of adequate size. The 26 accessions carrying the recombinant haplotype at HDR1-2 were excluded from this analysis to restrict the comparison to accessions carrying either the homozygous alternative (AA) or the reference (RR) haplotype at both loci.

The remaining accessions were classified into four groups based on their favourable (RR) or unfavourable (AA) haplotype allelic composition at HDR1-2/HDR4 loci (Table 1): Group I (AA/AA; n = 110, 44.9%), Group II (AA/RR; n = 31, 12.7%), Group III (RR/AA; n = 42, 17.1%), and Group IV (RR/RR; n = 62, 25.3%) (Table 1; Table S17).

**Table 1:** Multi-locus haplotypes combinations at HDR1-2 and HDR4 and their impact on grain yield under well-watered and water-limited conditions and grain yield loss.

| Group label | Group allelic composition | Alleles at HDRs |  | Count |  | Well-watered (WW) | Water-limited (WL) | GY loss (%)<sup>c</sup> |
| --- | --- | --- | --- | --- | --- | --- | --- | --- |
|  |  | HDR1-2<sup>a</sup> | HDR4<sup>a</sup> | No. of accessions | Freq. (%)<sup>b</sup> | emmeans GY (g m<sup>-2</sup>)<sup>c</sup> | emmeans GY (g m<sup>-2</sup>)<sup>c</sup> |  |
| I | AA/AA | AA | AA | 110 | 44.9 | 452.9 ± 11.8 | 291.9 ± 10.8 | 35.3 ± 1.9 |
| II | AA/RR | AA | RR | 31 | 12.7 | 442.2 ± 19.6 | 332.2 ± 17.9 | 24.6 ± 3.1 |
| III | RR/AA | RR | AA | 42 | 17.1 | 435.5 ± 16.3 | 318.6 ± 14.9 | 26.4 ± 2.6 |
| IV | RR/RR | RR | RR | 62 | 25.3 | 464.6 ± 17.8 | 354.5 ± 16.3 | 23.9 ± 2.8 |

<sup>a</sup>AA, homozygous alternative allele; RR, homozygous reference allele. <sup>b</sup>Frequency is expressed as a percentage of the total 245 accessions of the panel used in the modelling. <sup>c</sup>Values represent Estimated Marginal Means (EM-Means) ± Standard Error (SE) derived from mixed linear model.

Using a kinship-corrected mixed linear model, the overall haplotype-group effect on GY was not significant under WW conditions (Wald *χ*^2^ = 1.97, *P* = 0.580; Fig. S9, Table S17, but it was significant under WL conditions (Wald *χ*^2^ = 7.85, *P* = 0.049; Fig. 6A). Under WW conditions, no pairwise comparison among groups was significant (all *P_adj_* ≥ 0.61). Under WL conditions, Group IV (RR/RR; emmean GY = 354.5 ± 16.3 g m^-2^) showed significantly higher GY than Group I (AA/AA; 291.9 ± 10.8 g m^-2^; *P*-adj = 0.044). No other pairwise comparison under WL was significant (all *P_adj_* ≥ 0.24).

**Fig. 6.**
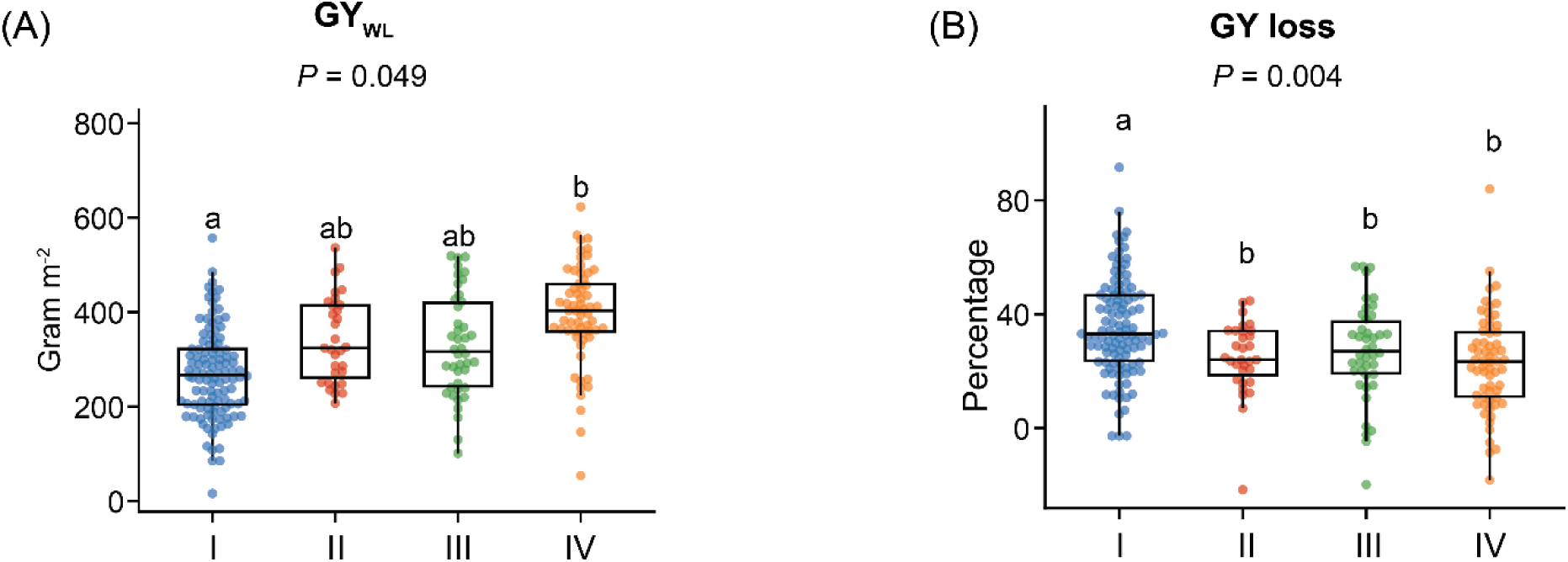
Multi-locus haplotype combinations at HDR1-2 and HDR4 and their impact on grain yield under water-limited conditions and grain yield loss. (A) Grain yield (GY, g m^-2^) under water-limited (GY_WL_) conditions for the four multi-locus haplotype groups (Table 1): Group I (AA/AA), Group II (AA/RR), Group III (RR/AA) and Group IV (RR/RR). (B) Percentage drought-induced grain yield loss for the same four groups. In both panels, box plots show the medians and interquartile ranges, overlaid with individual data points; different letters denote significant differences based on Tukey’s HSD groupings (α = 0.05) following a significant Wald chi-square test.

The overall haplotype-group effect onGY loss was significant (Wald *χ*^2^ = 12.84, *P* = 0.004; Fig. 6B; Table 1; Table S17). Group I accessions showed the greatest GY loss (emmean = 35.3 ± 1.9%), significantly exceeding Group II (24.6 ± 3.1%; *P_adj_* = 0.027), Group III (26.4 ± 2.6%; *P_adj_* = 0.048), and Group IV (24.0 ± 2.9%; *P_adj_* = 0.031). Groups II, III, and IV did not differ significantly from one another (all *P_adj_* ≥ 0.92).

Taken together, these results indicate that carrying the favourable reference haplotype allele at either HDR1-2 or HDR4 was associated with reduced drought-induced GY loss relative to the homozygous alternative allelic combination. However, carrying the reference allele at both loci conferred no additional benefit over carrying either allele alone suggesting a pattern inconsistent with a strictly additive, dose-dependent effect of the two loci. The effect on absolute GY was more limited: only the homozygous reference vs. homozygous alternative comparison (Group IV vs. Group I) was significant, and exclusively under WL conditions. The most robust signal from this locus combination is therefore the reduction in drought-induced GY loss associated with the favourable reference allele at either HDR1-2 or HDR4, rather than a cumulative increase in absolute GY.

## Haplotype-based anchoring of the *indica* mapping panel to the 3,000 Rice Genomes Project panel for detection of genetic variants at HDR1-2 and HDR4

To increase the genetic resolution of our SNP map (∼45K SNPs) and identify potential genetic variants underlying each HDR, haplotype-based anchoring was initially performed using the 3,000 Rice Genomes Project (3K RGP) accessions as a densely genotyped reference panel (∼29 million SNPs and ∼2.3 million insertions/deletions [INDELs]).

To anchor our low-density SNP map to the 3K RGP SNP map, we first assessed whether the mapping panel and the 3K RGP *indica* accessions were genome-wide genetically similar enough to share the more local haplotype variation at the detected HDRs. A PCA was conducted on a merged panel comprising the 271 accessions of our mapping panel and 1,785 *indica* or *indica*-admixed accessions from the 3K RGP (total = 2,056 accessions; PCA based on 3,954 shared, LD-pruned SNPs). PC1 and PC2 together explained 31.36% of the variation and showed extensive overlap between the mapping panel and the 3K RGP *indica* accessions, with no distinct separation between the two datasets (Fig. 7A), confirming that their suitability for haplotype-based anchoring. Furthermore, accessions separated broadly according to *indica* subpopulation and geographical origin, with the mapping panel distributed across the diversity space alongside the 3K RGP ind1, ind2 and ind3 groups rather than forming its own distinct cluster (Fig. S10). On this basis, the 500 3K RGP *indica* accessions genetically closest to the 271 accessions of our mapping panel (based on Euclidean distances in the top 10 PCs space of the SNP-based PCA), were selected as a reference subset for all downstream analyses, providing dense variant coverage from accessions closely related to our panel (Table S18 and S19). All subsequent variant analyses drew on the full genetic variant data (5,199,884 SNPs and 384,934 INDELs) available for the 500 3K RGP accessions, a dataset with a highly increased resolution than our GWA mapping SNP map (45,505 SNPs).

**Fig. 7.**
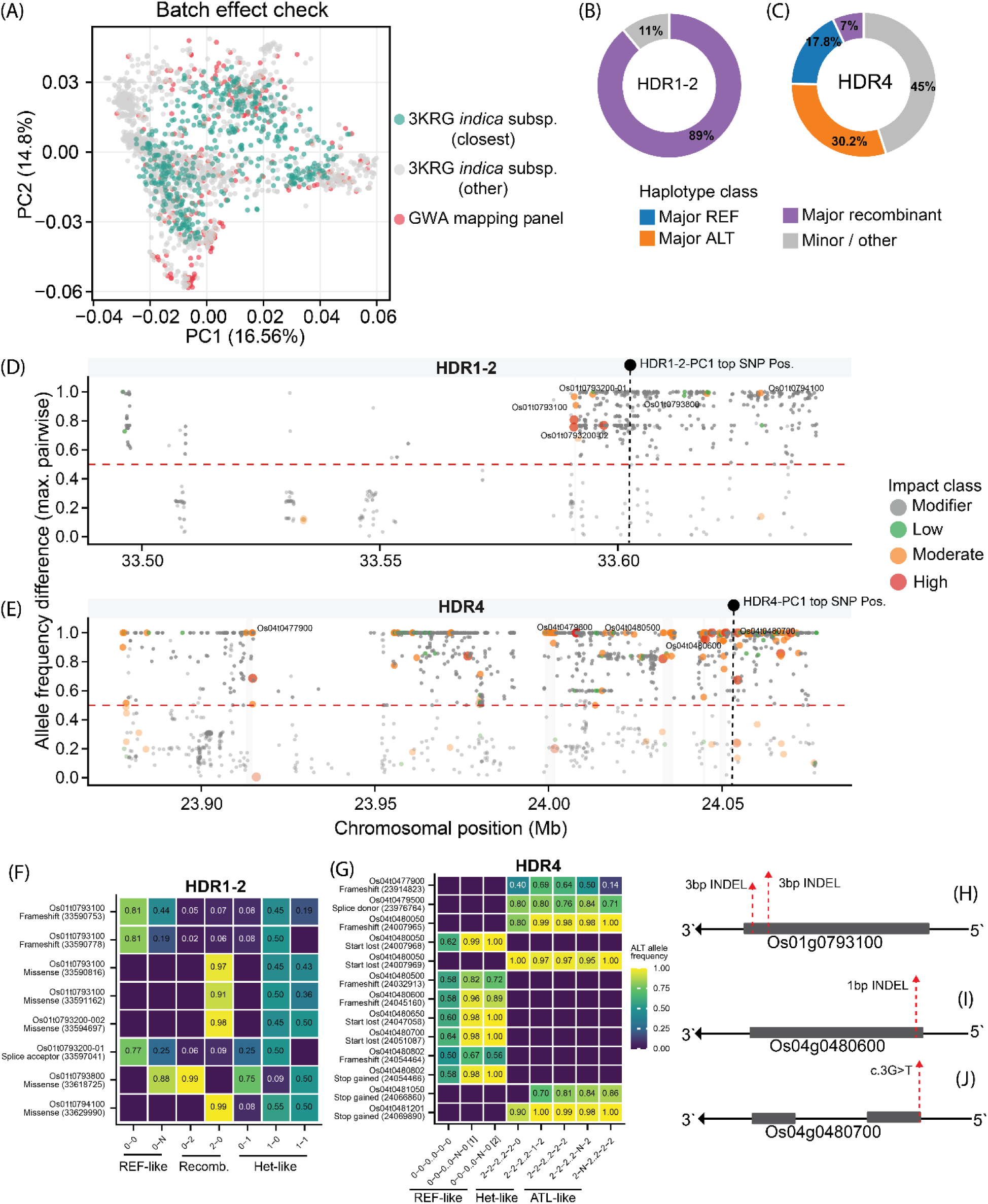
Haplotype-based HDR loci’s variant detection using the 3,000 Rice Genomes Project dataset as a reference panel. (A) Principal component analysis (PCA) score plot of 2,056 *indica* accessions based on the 3K RGP *indica* subspecies and our GWA mapping panel. Accessions are coloured based on their data source. (B, C) Donut charts showing haplotype class frequencies among the 500 closest 3K RGP *indica* reference accessions at (B) HDR1-2, and (C) HDR4. Major reference (REF; blue), major alternative (ALT; gold), major recombinant (purple), and minor/other (grey) haplotype classes are indicated. (D, E) Maximum pairwise allele frequency differences between major haplotype groups for all annotated variants within the haplotype blocks of (D) HDR1-2 and (E) HDR4, plotted against chromosomal position. Variants are coloured by SnpEff functional impact class: HIGH (red), MODERATE (orange), LOW (green), and MODIFIER (grey). The dashed red line indicates the 0.5 threshold for haplotype enrichment. The dashed vertical black line marks the HDR-PC1 msSNP position. Gene IDs for loci harbouring HIGH and MODERATE impact variants are labelled. (F) Heatmap of candidate genes with haplotype-enriched HIGH and MODERATE impact variants at HDR1-2. (G) Heatmap of candidate genes with haplotype-enriched, HIGH impact variants at HDR4. The ALT allele frequency is indicated within each heatmap box. (H, I, J) Schematic gene models for three priority candidate genes: (H) *Os01g0793100* (polyphenol oxidase) showing a 4-bp deletion and 4-bp insertion; (I) *Os04g0480600* (*CYP71S3*) showing a 1-bp insertion; (J) *Os04g0480700* (*CYP71S2*) showing a missense variant (c.3G>T). Grey boxes represent exons; arrows indicate the direction of transcription; red arrowheads mark the approximate position and type of each variant.

Haplotype-based anchoring was performed at the target HDRs (HDR1-2 and HDR4) by matching the GWA mapping SNPs that defined each haplotype block (Fig. 4C and Fig. 5C) against the 3K RGP genotypes; the SNPs shared between the two panels at each HDR are listed in Data S2. HDR1-1 and HDR11 were excluded from this set of analyses as they were not among our focal HDRs.

At HDR1-2 (2 of the 6 SNPs present), recombinant haplotypes predominated (n = 445; 89.0%), with minor haplotypes accounting for the remaining accessions (n = 55; 11.0%); notably, one of these minor haplotypes carried the reference allele at both SNP positions (n = 13; 2.6%), corresponding to the reference haplotype identified in the GWA mapping panel (HAP2, Fig. 7B; Fig. 4C). At HDR4 (18 of the 20 SNPs present), both haplotypes originally defined in the GWA mapping panel (HAP1, alternative, and HAP2, reference; Fig. 5C) were recovered among the 500 reference accessions: a major alternative haplotype (n = 151; 30.2%), a reference haplotype (n = 89; 17.8%) and a recombinant haplotype (n = 35; 7.0%) (Fig. 7C; Table S20 and S21).

Across the two anchored haplotype blocks, 2,340 variants were enriched between major haplotype groups (alternative-allele frequency difference ≥ 0.5): 487 at HDR1-2 and 1,853 at HDR4 (Table S21). Variants were classified by their predicted functional effect: HIGH and MODERATE impact variants alter the encoded protein, including frameshifts, premature stop codons, splice-site disruptions (HIGH), and missense substitutions (MODERATE). In contrast, MODIFIER variants fall in non-coding or regulatory regions (e.g. introns, UTRs, promoters), and LOW impact variants are synonymous. Of the 2,340 enriched variants, 141 were classified as HIGH or MODERATE impact, 129 as LOW impact, and 2,070 as MODIFIER.

At HDR1-2, the 42.4-kbp LD block (the same block reported in Fig. 4C) contained 487 enriched variants, of which eight were of HIGH or MODERATE impact, mostly located in two genes (Fig. 7D, F; Table S21). *Os01g0793100*, encoding a polyphenol oxidase (PPO), carried two frameshift mutations (p.Arg302fs, p.Ala295fs; HIGH) and two missense substitutions (p.His282Gln, p.Ala167Val; MODERATE; Fig. 6H). *Os01g0793200*, a pentatricopeptide-repeat protein gene, carried one missense and one splice-acceptor variant, and additional missense variants were found in *Os01g0793800* and *Os01g0794100* (Fig. 7F; Fig. S105A; S107A; Table S22). These variants were partitioned between haplotypes (shown as per-haplotype alternative-allele frequency in Fig. 7F). The two PPO frameshifts and the *Os01g0793200* splice-related HIGH-impact variant were present almost exclusively in the minor reference haplotype ("0-0"; n = 13, 2.6%; corresponding to HAP2 in Fig. 4C), where their alternative-allele frequency reached 0.81, compared with below 0.08 in each of the two recombinant haplotypes ("0-2" and "2- 0"). By contrast, the four MODERATE-impact missense variants were distributed across the two recombinant haplotypes: three (in *Os01g0793100*, *Os01g0793200* and *Os01g0794100*) were enriched in "2-0," and one (in *Os01g0793800*) was enriched in "0-2." This suggests that the divergence between the major haplotypes in this HDR is associated with missense variation across several genes, whereas the PPO frameshift disruptions are restricted to the rare reference haplotype.

At HDR4, within the 98-kbp LD block (the same block reported in Fig. 5C), 133 of the enriched harboured variants were of HIGH or MODERATE impact (Fig. 7G, I). The highest densities of haplotype-based differentiation fell within the tandem cytochrome P450 genes *CYP71S2* (*Os04g0480700*; 18 missense, 1 start-lost variant; Fig. 7J) and *CYP71S3* (*Os04g0480600*; 17 missense, 1 frameshift; Fig. 7I), followed by an LRR receptor-like protein gene (*Os04g0480500*; 16 missense, 1 frameshift), an F-box protein gene (*Os04g0479800*; 12 missense), a cysteine-proteinase precursor gene (*Os04g0477900*; 9 missense, 1 frameshift) and a glycoside hydrolase gene (*Os04g0480900*; 8 missense; Fig. 7G). The 13 HIGH-impact variants were distributed across both major haplotypes, with loss-of-function variants enriched in different genes: the reference-like haplotype carried loss-of-function variants concentrated mainly in the *CYP71S2/CYP71S3* cluster and flanking genes, while the alternative-like haplotype carried its own loss-of-function variants in upstream (*Os04g0479500*, *Os04g0480050*) and downstream (*Os04g0481050*, *Os04g0481201*) genes (Fig. S106). This bidirectional distribution of loss-of-function variants across both haplotypes suggests that each haplotype may disrupt a different subset of genes within this tandemly duplicated cluster (Fig. S106; S107B; Table S23).

## Discussion

Grain yield, especially under drought stress, is a low-heritability, complex endpoint trait controlled by many loci with minor effects that are highly environment-dependent (Blancon et al. 2024). This complex genetic architecture limits the resolution and power of conventional, yield-based mapping approaches. To circumvent these limitations, this study demonstrates that rice flag leaf primary metabolites, measured under field-based drought conditions, serves as genetically tractable intermediate phenotypes. Utilizing these intermediate metabolic proxies enables the identification of moderate-effect loci that are directly linked to grain yield stability under stress.

## Flag leaf metabolic components as genetically tractable intermediates physiologically connected to rice performance

We previously demonstrated, utilizing the same 271 *indica* rice accessions evaluated in this study, that the flag leaf metabolic profiles of drought-stressed plants predict grain yield (GY) and stress-induced GY loss more accurately than genomic markers alone (Melandri et al. 2022). The physiological basis for this higher prediction accuracy is clear: the flag leaf is the primary photosynthetic source for the developing grains in the panicle, contributing roughly half of the assimilates used for grain filling; thus, its metabolic profile is directly linked to yield formation (Li et al. 1998); Yoshida, 1981). Metabolite levels also integrate post-transcriptional and post-translational regulation, enzyme kinetics, allosteric control, substrate channelling, and environmental influences, positioning them closer to functional phenotypic outputs, such as GY, than genetic markers (Fiehn 2002; Matsuda et al. 2015). These characteristics support the role of flag leaf metabolites as intermediate phenotypes (endophenotypes) that link genotype to yield, especially under stress.

The present study dissects the genetic architecture of the rice flag leaf metabolic variation. While genome-wide association (GWA) mapping identified 180 metabolic quantitative trait loci (mQTLs) for 89 polar metabolites, it detected only four low-significance QTLs for GY (Fig. 2). This asymmetry reflects a fundamental difference in the genetic architecture of leaf metabolites versus GY. The latter is an endpoint trait shaped by hundreds-to-thousands of small-effect loci (Huang et al. 2010; Yano et al. 2016), whereas metabolite levels are governed by fewer loci with moderate-to-large effects (Wen et al. 2014; Chen et al. 2014; Zhu et al. 2018). Furthermore, metabolites were more heritable in our panel: 46.0% and 29.0% of metabolite traits exceeded the narrow-sense heritability (*h*^2^) of GY under water-limited (WL) and well-watered (WW) conditions, respectively. (Table S7).

Notably, 51.6% of the metabolic QTLs (mQTLs) were detected exclusively under WL conditions, indicating predominantly stress-dependent genetic control. Sampling flag leaves at the peak of the 14-day water-withholding period captured a metabolic state of stress characterized by reduced photosynthetic and tricarboxylic acid (TCA) cycle activities alongside the activation of stress-responsive pathways. Only under WL conditions the genetic variants that modulated these metabolic shifts produced phenotypic differences large enough to exceed the detection threshold. Consistent with this, sample scores on the first principal component (PC1) of the flag leaf metabolome correlated significantly with GY only under WL conditions (*r_s_* = 0.51, *P* < 0.001; Table S5), confirming that this stress-specific variation is intrinsically associated with yield performance.

## High-density regions of mQTLs as metabolic hubs linked to grain yield

Building on these observations, we prioritized high-density regions (HDRs) of mQTLs (defined as ≥4 overlapping mQTLs for distinct metabolites). We reasoned that such hubs were more likely to contain variants affecting multiple metabolic pathways than isolated mQTLs (Khahani et al. 2021; Daryani et al. 2024) and were therefore more likely to influence crop performance. Supporting this rationale, all four HDRs that were identified in the present study were significantly associated with GY (Fig. 3E - H). To connect these metabolic hubs more directly to GY, we performed a second round of GWA mapping using the region-specific PC1 score of each HDR’s constituent metabolites as a synthetic phenotype, followed by linkage disequilibrium characterization and haplotype analysis. GWA mapping utilizing PCA-derived synthetic phenotypes is a strategy previously applied in rice to increase the GWAS power for detection of genes comprehensively controlling multiple traits (Yano et al. 2019).

PC1-based mapping recovered a single associated locus per hub that co-localized with or fell within the interval defined by the individual mQTLs (Fig. 4-5; S4, and S7), indicating that each hub represents a coherent genetic unit. The PC1 proxies of all three WL-specific hubs were also significantly correlated with GY under water limitation (Fig. 3E-H), establishing a direct link between each hub’s collective metabolic variation and yield performance. Each hub’s constituent metabolites point to a distinct physiological process connected to grain yield under drought, as detailed below.

HDR1-1 was the only hub detected under WW conditions. It captured constitutive variation in galactinol and myo-inositol, both of which are related to the raffinose-family oligosaccharide (RFO) pathway (Elango et al. 2022), together with the starch-derived disaccharide isomaltose (Table S9). In contrast, the three WL-specific HDRs each captured a distinct facet of stress-induced metabolic reprogramming. The metabolites mapping to HDR1-2 showed opposite associations with GY: citric and isocitric acid correlated positively, whereas proline and tyramine correlated negatively (Fig. 3F). This pattern is consistent with a broader contrast under water-deficit stress. A sustained TCA cycle (citric and isocitric acid) is associated with yield stability (Obata and Fernie 2012; Melandri et al. 2021), while the accumulation of stress-responsive amino acids such as proline (Alvarez et al. 2022) and amino acid derivatives such as tyramine (Dos Santos et al. 2026) reflects the severity of the stress experienced and its negative effect on yield.

HDR4 was mapped for variation in the coordinated accumulation of branched-chain amino acids (BCAAs) together with threonine, ornithine, and allantoin. BCAAs accumulation is a hallmark of drought and carbohydrate starvation, conditions under whichprotein turnover releases free amino acids that can be catabolized as alternative respiratory substrates, feeding electrons into the mitochondrial electron-transport chain (Araújo et al. 2011; Hildebrandt et al. 2015; Shim et al. 2023). The negative association between BCAA levels and GY (Fig. 3G) is consistent with these amino acids marking a stress state of net protein degradation, and this locus reveals the presence of genetic variation underlying this response.

HDR11, associated with rare allelic variation, was mapped for a broad and generalized amino acid reprogramming, with 13 amino acids across multiple biosynthetic families associated with this hub (Fig. S8). Such a large and coordinated response is characteristic of large-scale proteolysis and senescence-like nitrogen remobilization under severe stress (Hildebrandt 2018), and the strong negative association of the rare allele with GY suggests it marks a metabolic-based susceptibility to drought stress present in only a few accessions of the panel.

By collapsing each hub’s multi-metabolite variation into a single genetically anchored axis tied to yield, this approach provided the basis for testing whether the favourable alleles at these hubs combine additively to affect grain yield.

## Favourable alleles of main haplotypes at two metabolic QTL hubs buffer drought-induced yield loss, but do not act additively

Whether the WL-specific hubs mapped in this study contributed independently to yield, and whether their effects can be combined, determines their usefulness for future breeding efforts aimed at improving rice yield stability under drought. This analysis was restricted to HDR1-2 and HDR4, the two hubs with allele frequencies that provided adequate group sizes for a two-locus contrast. After accounting for genetic relatedness among accessions, those carrying the favourable haplotype allele at both loci (Group IV) lost significantly less yield under WL stress than accessions carrying the alternative allele at both loci (Group I; 11.4% points less, Table 1; Table S17) and also exhibited significantly higher absolute GY under WL conditions (a difference of 62.5 g m^-2^, Table 1; Table S17). Notably, accessions carrying the favourable allele at only one locus (Groups II and III) exhibited a GY loss that was statistically indistinguishable from Group IV, yet significantly lower than Group I (Figure 6B). Concurrently, GY itself did not differ significantly among Groups II, III, and IV. This pattern is not consistent with a simple additive, dosage-dependent model, under which yield stability would be expected to improve progressively from one favourable allele to two. We note that this analysis compared haplotype-group means directly and did not formally test for a genotype-by-genotype interaction between HDR1-2 and HDR4; we therefore do not characterise the relationship between the two loci as either additive or epistatic. Instead, we describe the observed pattern as functionally redundant, in the sense that carrying either favourable allele appears sufficient to realise most of the benefit.

The clearest and most robust result from this analysis is the11.4%-point reduction in stress-induced GY loss associated with the favourable allele at either locus. This represents a direct gain in reproductive-stage yield stability, which is the most effective selection target when breeding for drought-prone rainfed systems (Venuprasad et al. 2007; Kumar et al. 2014). If this redundancy is confirmed in independent populations, it would simplify rather than complicate breeding deployment: selecting for the favourable allele at either HDR1-2 or HDR4 alone, rather than requiring both to be combined, may be sufficient to capture most of the yield-stability benefit. Together, these results indicate that metabolite-guided mapping can deliver yield-relevant markers that yield-based mapping alone did not detect in this panel. As with any comparison across haplotype groups in a structured diversity panel, validation in biparental or backcross populations, ideally with a factorial design capable of formally testing the HDR1-2-by-HDR4 interaction, is required before either locus is deployed in a marker-assisted selection scheme.

## Haplotype anchoring to the 3K Rice Genomes Project panel nominates candidate genes and variants underlying the metabolic hubs

A key methodological feature of this study is the use of HDR haplotypes as anchors linking our low-density genotyping-by-sequencing (GBS) SNP map to the whole-genome sequence (WGS) resource of the 3K RGP. Rather than attempting fine-scale mapping within our own sparsely genotyped panel, we used the haplotypes defining each hub to identify closely related WGS-sequenced 3K RGP accessions and mined those accessions for candidate variants. This approach borrows the logic of imputation-based association, wherein a dense reference panel resolves variation beyond the density of the study genotypes, an approach applied both in human genetics (McCarthy et al. 2016) and in plants (Cao et al. 2011). This strategy was applicable at two of the four hubs and yielded promising candidate variants at both hubs. The genes discussed below are hypothetical candidates for functional testing rather than validated causal loci.

At HDR1-2, the variant analysis identified a candidate gene that is mechanistically relevant to the mapped metabolites. *Os01g0793100*, which encodes a polyphenol oxidase (PPO), carried two frameshift mutations (p.Arg302fs and p.Ala295fs) enriched in the reference-like haplotype. Both mutations shift the reading frame upstream of the conserved copper-binding region and are predicted to yield a non-functional enzyme (Fig. 7D, F, H). Plant PPOs oxidize mono- and diphenolic substrates, and tyramine is a documented monophenolic PPO substrate (Kampatsikas et al. 2017). A simple substrate-oxidation model does not, however, account for the observed direction of this effect: the reference-like haplotype carries the disrupted PPO yet accumulates less tyramine, whereas the intact PPO haplotype accumulates more (Fig. 3F; Fig. S5). Consequently, a simple loss of PPO functional activity does not straightforwardly explain elevated tyramine. If PPO contributes to this phenotype, its effect is more likely to act indirectly, such as through negative feedback regulation of the substrate or by diverting it to alternative pathways. At the phenotypic level, the intact PPO haplotype (HAP1) also accumulated higher proline but lower levels of citric and isocitric acid, whereas the reference-like haplotype displayed the inverse profile and reached higher GY under water limitation. Because the PPO frameshifts co-occur with MODERATE-impact missense variants distributed across several other genes in the block (Fig. 7F), haplotype divergence likely reflects combined coding variation at multiple genes, with the PPO frameshifts representing the most prominent single change.

At HDR4, loss-of-function variants were partitioned between the two major haplotypes, concentrating in different genes within each lineage (Fig. 7E, G). The reference-like haplotype, which is higher-yielding and associated with lower amino acid levels, carried disrupting variants in the tandem cytochrome P450 pair *CYP71S3* (*Os04g0480600*; frameshift p.Leu18fs; Fig. 6J) and *CYP71S2* (*Os04g0480700*; start-lost; Fig. 7J), together with a frameshift in the adjacent LRR receptor-like protein gene *Os04g0480500* (p.Ala121fs). Conversely, the alternative-like haplotype carried stop-gained and frameshift variants in flanking genes (*Os04g0481050*, p.Glu330*; *Os04g0481201*, p.Arg138*; *Os04g0480050*, p.Cys2fs). Because each lineage independently carries loss-of-function variants in different gene members of the region, the observed pattern is consistent with the divergent resolution of a tandemly duplicated gene array, a common fate of duplicate genes (Lynch and Conery 2000; Taylor et al. 2001). *CYP71* group P450s act largely in amino acid-derived specialized metabolism, as exemplified by CYP71P1, which hydroxylates tryptamine to the antioxidant serotonin (Fujiwara et al. 2010), thereby channelling amino acid-derived substrates into stress-protective metabolites. *CYP71S2* and *CYP71S3* are characterized as aldoxime-metabolizing enzymes acting on indole-3- acetaldoxime (Nelson and Werck-Reichhart 2011; Irmisch et al. 2014). However, the amino acids mapped to HDR4 (branched-chain amino acids, threonine, and ornithine) are not established substrates of characterized *CYP71* enzymes. We therefore hypothesize that the differential disruption of this P450 pair could redirect metabolic flux through amino acid-derived metabolism, contributing to the difference in amino acid accumulation between the haplotype groups (Fig. S6). Another link to precursor supply is offered by the co-localization of *Os04g0479200*, encoding an isocitrate dehydrogenase that generates the 2-oxoglutarate skeleton for glutamate, the amino donor for the amino acids accumulating at this locus.

## Conclusions

In summary, flag leaf metabolite-based GWA mapping resolved genetic loci linked to grain yield under drought that conventional yield-based mapping could not detect in this panel. These loci include two hubs, HDR1-2 and HDR4, whose favourable alleles are each independently associated with reduced yield loss and higher yield under water limitation, with no additional benefit gained from combining both. Together, metabolite-guided GWA mapping framework deployed in this study offers directly actionable markers for improving drought resilience in *indica* rice breeding. Furthermore, it provides a transferable strategy for dissecting complex stress-tolerance traits in other crops where yield-based mapping alone lacks the resolution to identify candidate loci.

## Data availability

All supplementary data (Data S1 - S5) and files associated with this manuscript is available via this link: https://doi.org/10.6084/m9.figshare.32903357

## Acknowledgements Funding

This work was supported by Giovanni Melandri’s startup funds from the School of Plant Sciences, College of Agriculture, Life & Environmental Sciences of the University of Arizona.

## Author information

Giovanni Melandri ORCID: 0000-0002-0877-5009 Chosen Ekene Obih ORCID: 0000-0002-9387-0632

## Authors and Affiliations

School of Plant Sciences, University of Arizona, Tucson, AZ, USA Chosen Ekene Obih; Giovanni Melandri

Genetics Graduate Interdisciplinary Program, University of Arizona, Tucson, AZ, USA Zachary Dickens

Center for Agroecosystem Research in the Desert (ARID), University of Arizona, Tucson, AZ, USA

Giovanni Melandri

## Contributions

GM and CEO conceived the research, GM conducted the field and wet-lab experiments, CEO and ZD analysed the data. CEO and ZD drafted the manuscript. CEO and GM finalized the manuscript. All the authors approved the manuscript.

## Supplementary information

Supplementary file1 (Table S1 - S13) Supplementary file2 (Table S14 - S23) Supplementary file3 (Fig. S1 - S104) Supplementary file4 (Fig. S105 - S107)

