## Supplementary file3 (Fig. S1 - S104) for "GWAS of flag leaf primary metabolites uncovers metabolic QTL hubs linked to grain yield stability under drought in field-grown *indica* rice"

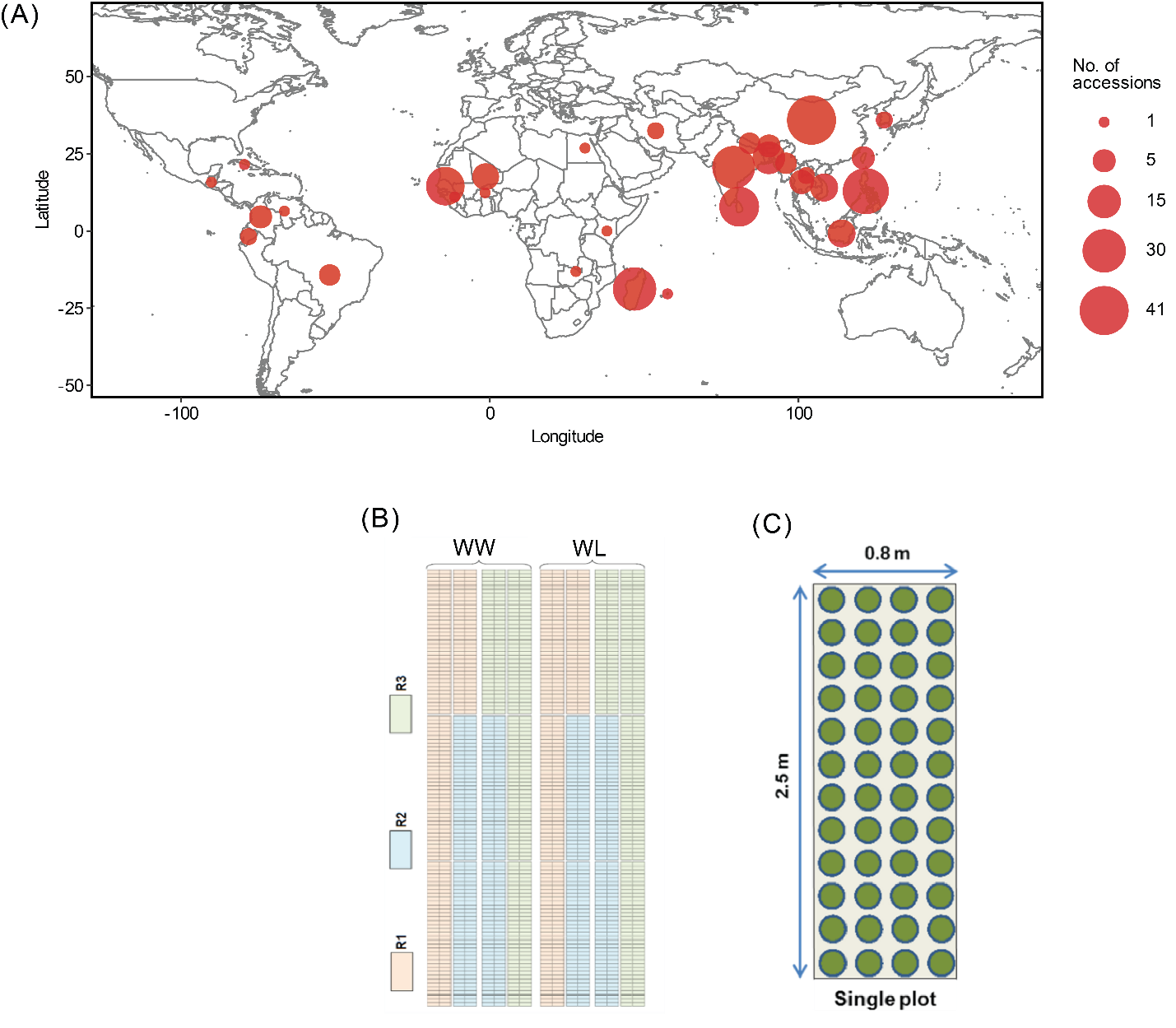

**Supplementary Fig. S1. Geographical distribution of the *indica* panel and experimental design of the field trial**

(A) Global distribution of the origin of the 271 accessions of the *indica* panel. Red circles are located on the countries of origin; sizes vary based on number of accessions collected from that country. (B) Experimental design showing the well-watered (WW) and water-limited (WL) treatment fields and the three replicated blocks (R1, R2 and R3) of the entire panel in each field. (C) Design and size of a single accession plot comprising 48 plants that correspond to each rectangle visible in the experimental design.

(A)	(B)

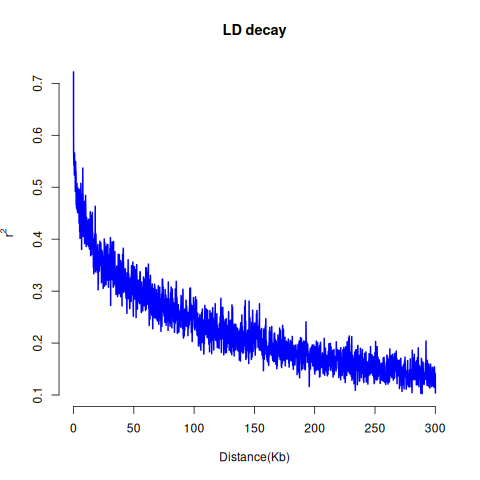

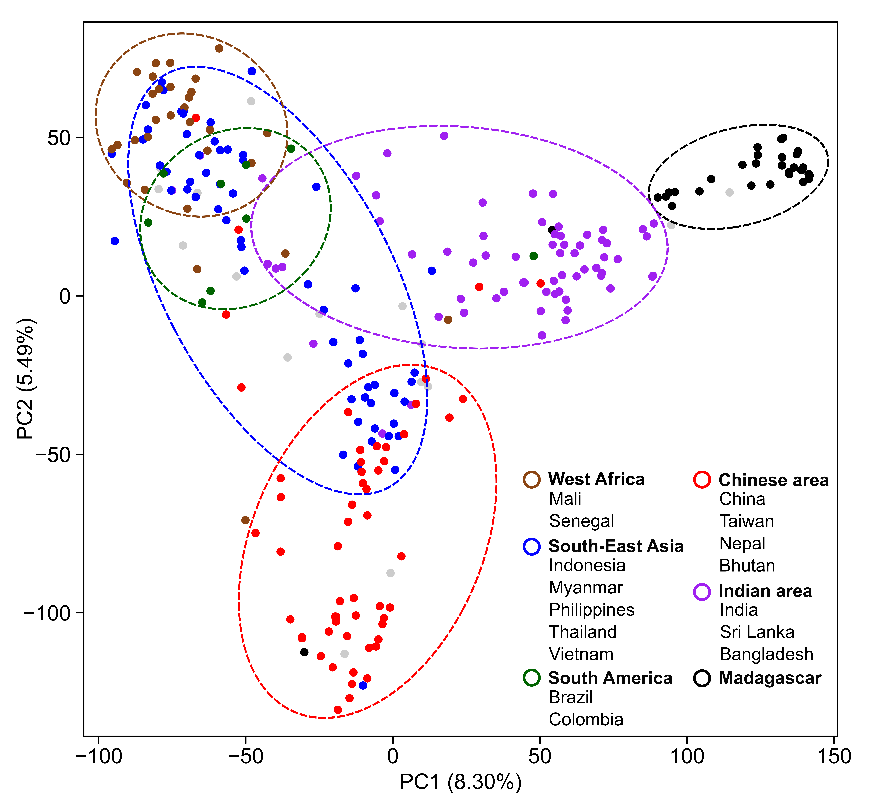

**Supplementary Fig. S2. SNP-based principal component analysis (PCA) score plot and genome-wide linkage disequilibrium (LD) decay plot**

(A) PC1-and PC2 score plot of the 271 accessions of the *indica* panel based on the 45,505 SNPs. Accessions are coloured based on their region of origin. (B) Genome-wide LD decay plot of the *indica* panel based on the 45,505 SNPs. The distance where the average pairwise LD (*r^2^*) declined to 0.2 (indicated by dashed red lines), at approximately 100 kb was used to as one of the criteria to define a quantitative trait locus (QTL).

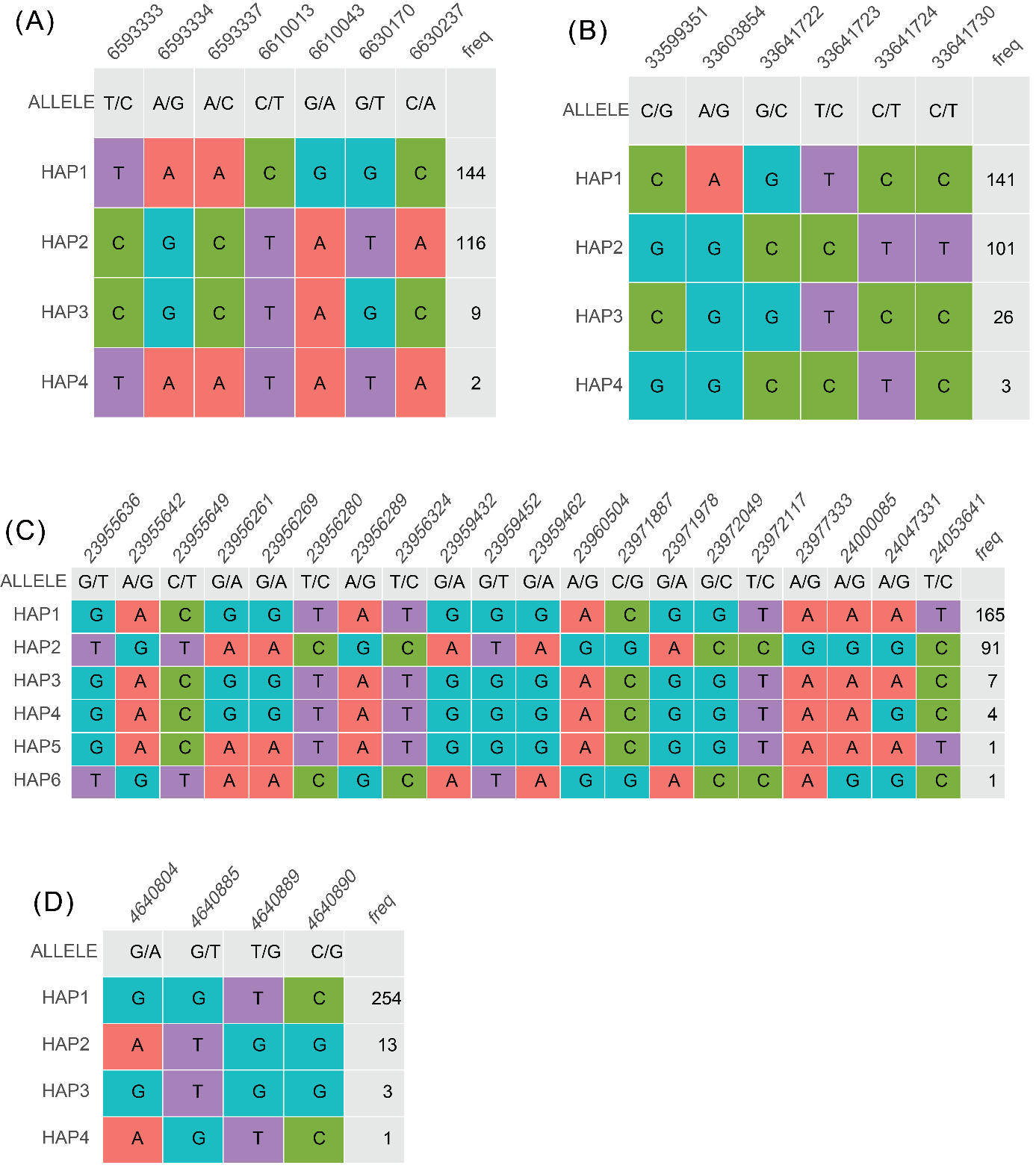

**Supplementary Fig. S3. Haplotype blocks of the HDRs for the 271 accessions**

(A) Haplotype block structure at HDR1-1. (B) Haplotype block structure at HDR1-2. (C) Haplotype block structure at HDR4. (D) Haplotype block structure at HDR11. Freq. represents the number of accessions in that haplotype.

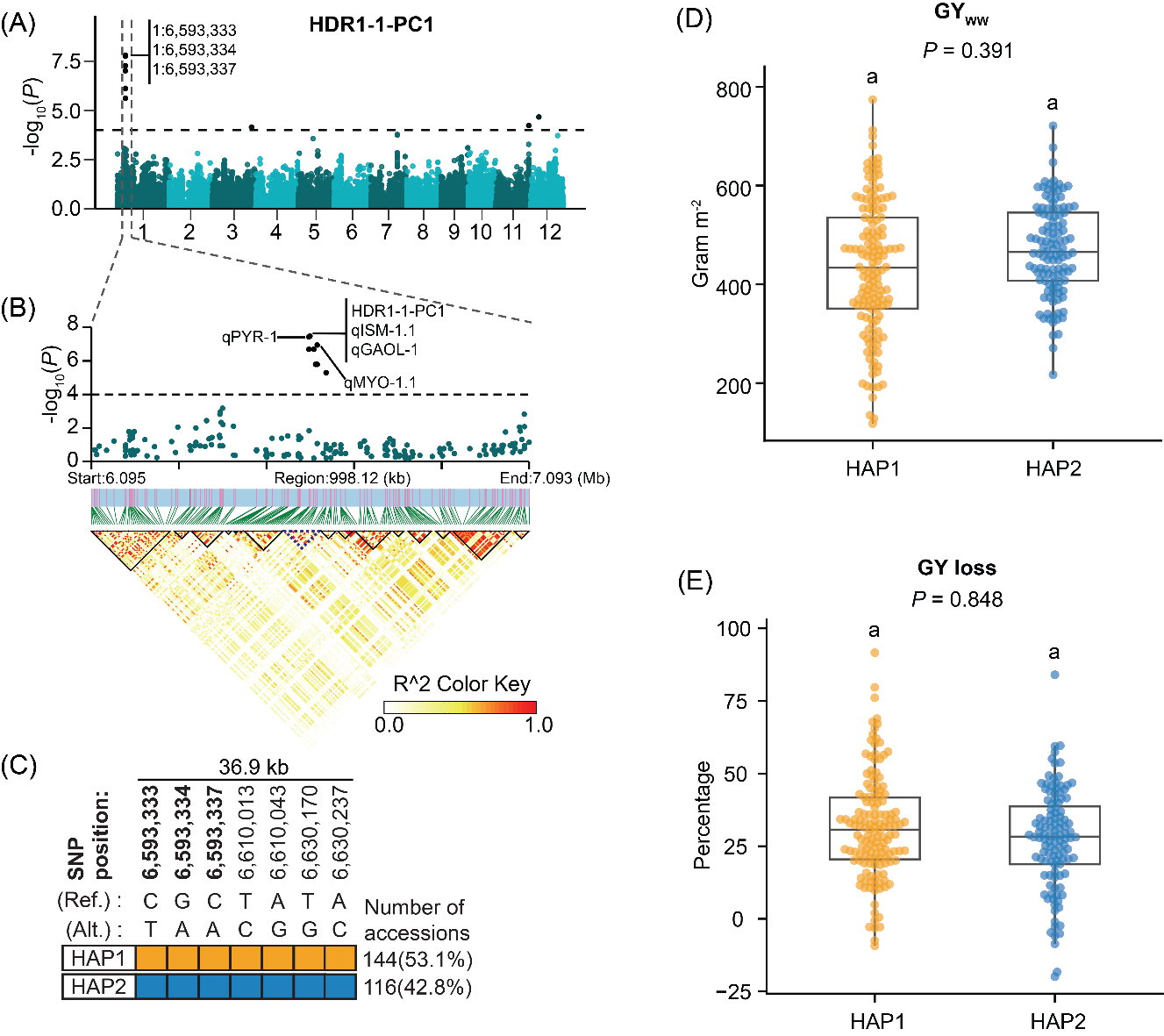

**Supplementary Fig. S4. Haplotype analysis, grain yield and grain yield loss associations at HDR1-1 on chromosome 1.**

(A) Manhattan plot of GWA mapping for the HDR1-1 PC1 metabolic proxy under well-watered (WW) conditions. The most significant SNP (msSNPs) on chromosome 1 are indicated. The dashed line represents the significant threshold (-log_10_(*P*) = 4.0). (B) Regional association plot (upper) and pairwise LD heatmap (lower, *r^2^* colour key) for the HDR1-1 local LD interval (Chr 1: 6.095-7.093 Mb). Individual metabolite QTLs msSNPs are labelled. (C) Haplotype block structure at HDR1-1, comprising 7 SNPs spanning 36.9 kb. The most significant SNPs (6,593,333; 6,593,334; 6,593,337) are indicated in bold. (D) Grain yield (GY) comparisons between haplotypes under well-watered conditions. (E) GY loss comparison between haplotypes. Reference and alternative alleles are shown; gold and blue jitters denote alternative and reference allelic composition of HAP1 and HAP2, respectively. *P*-values are from Wald chi-square test; different letters denote Tukey-HSD pairwise contrasts (α = 0.05). Box plots show medians, interquartile ranges and individual data points.

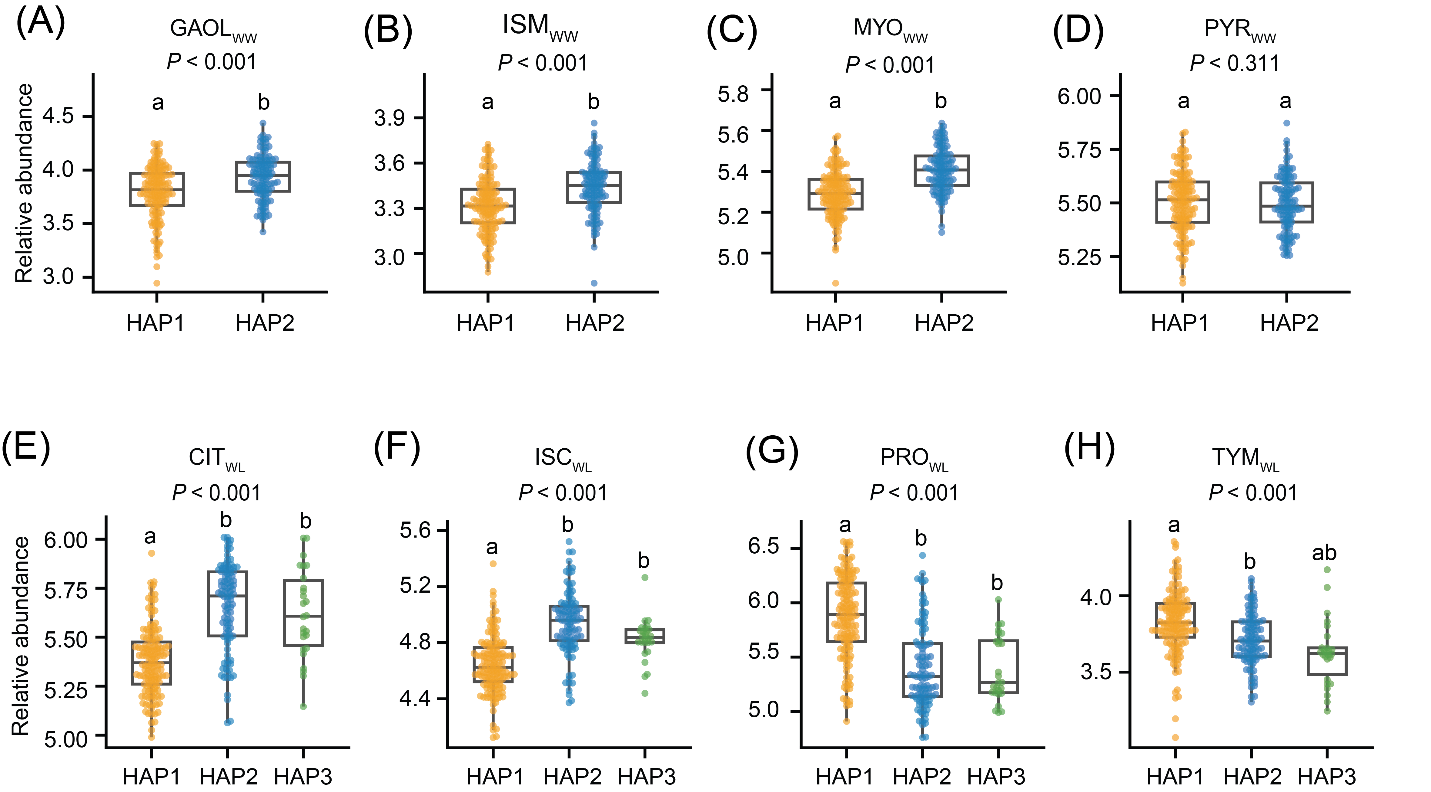

**Supplementary Fig. S5. Haplotype-trait associations at HDR1-1 and HDR1-2 loci on chromosome 1.**

Haplotype-trait comparisons at HDR1-1 for (A) galactinol, (B) isomaltose, (C) myo-inositol, and (D) pyroglutamic acid, all under well-watered (WW) condition for HAP1 (n = 144) and HAP2 (n = 116). (E-H) Haplotype-trait comparisons at HDR1-2 for (E) citric acid, (F) isocitric acid, (G) proline and (H) tyramine, all under water-limited (WL) condition for HAP1 (n = 141), HAP2 (n = 101) and HAP3 (n = 26); colours correspond to HAP1 (gold), HAP2 (blue) and HAP3 (green). *P-*values are from Wald chi-square test; different letters denote significant differences based on Tukey's HSD pairwise contrasts (α = 0.05). Box plots show medians, interquartile ranges, and individual data points. Box plots show medians, interquartile ranges and individual data points.

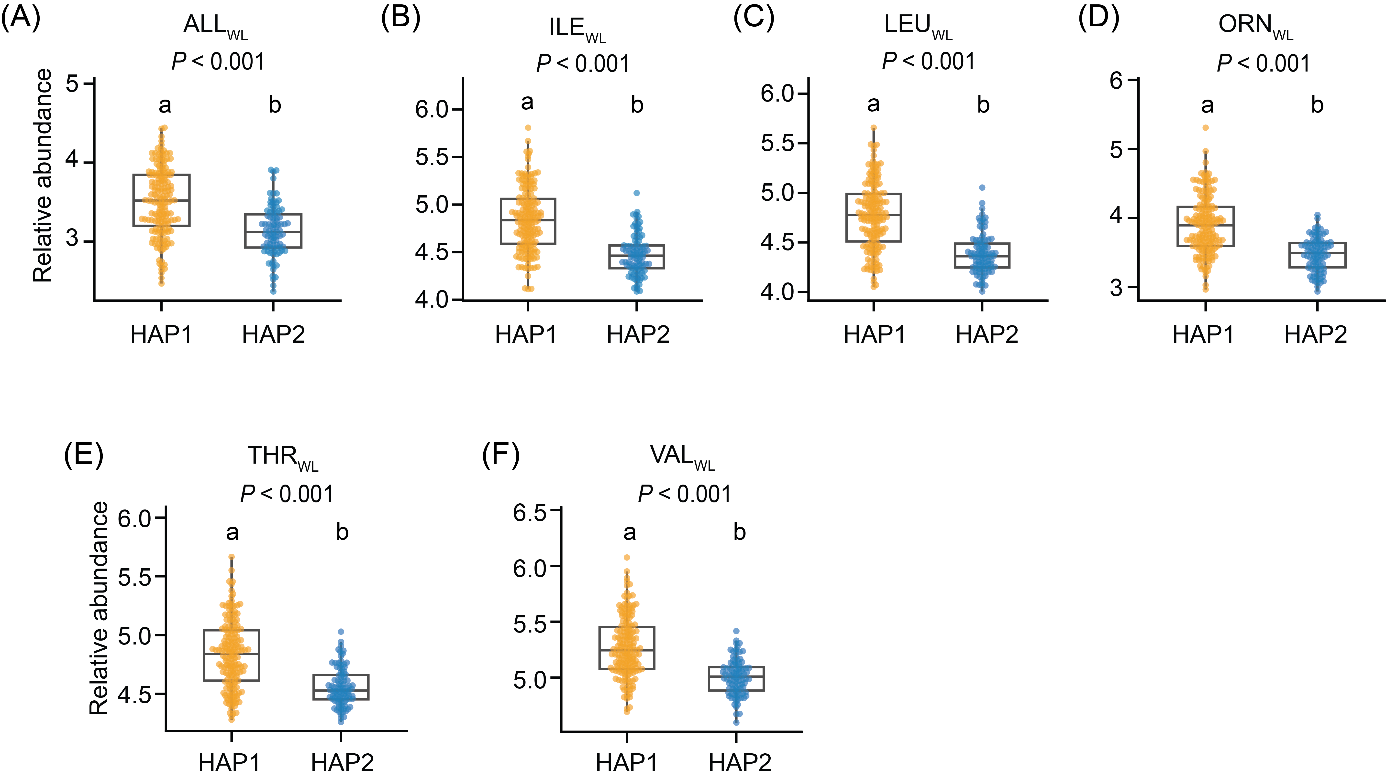

**Supplementary Fig. S6. Haplotype-trait associations at HDR4 loci on chromosome 4.**

(A) allantoin, (B) isoleucine, (C) leucine, (D) ornithine, (E) threonine and (F) valine, all under water-limited (WL) condition for HAP1 (n = 165) and HAP2 (n = 91). Colours correspond to HAP1 (gold) and HAP2 (blue). *P-*values are from Wald chi-square test; different letters denote significant differences based on Tukey's HSD pairwise contrasts (α = 0.05). Box plots show medians, interquartile ranges, and individual data points.

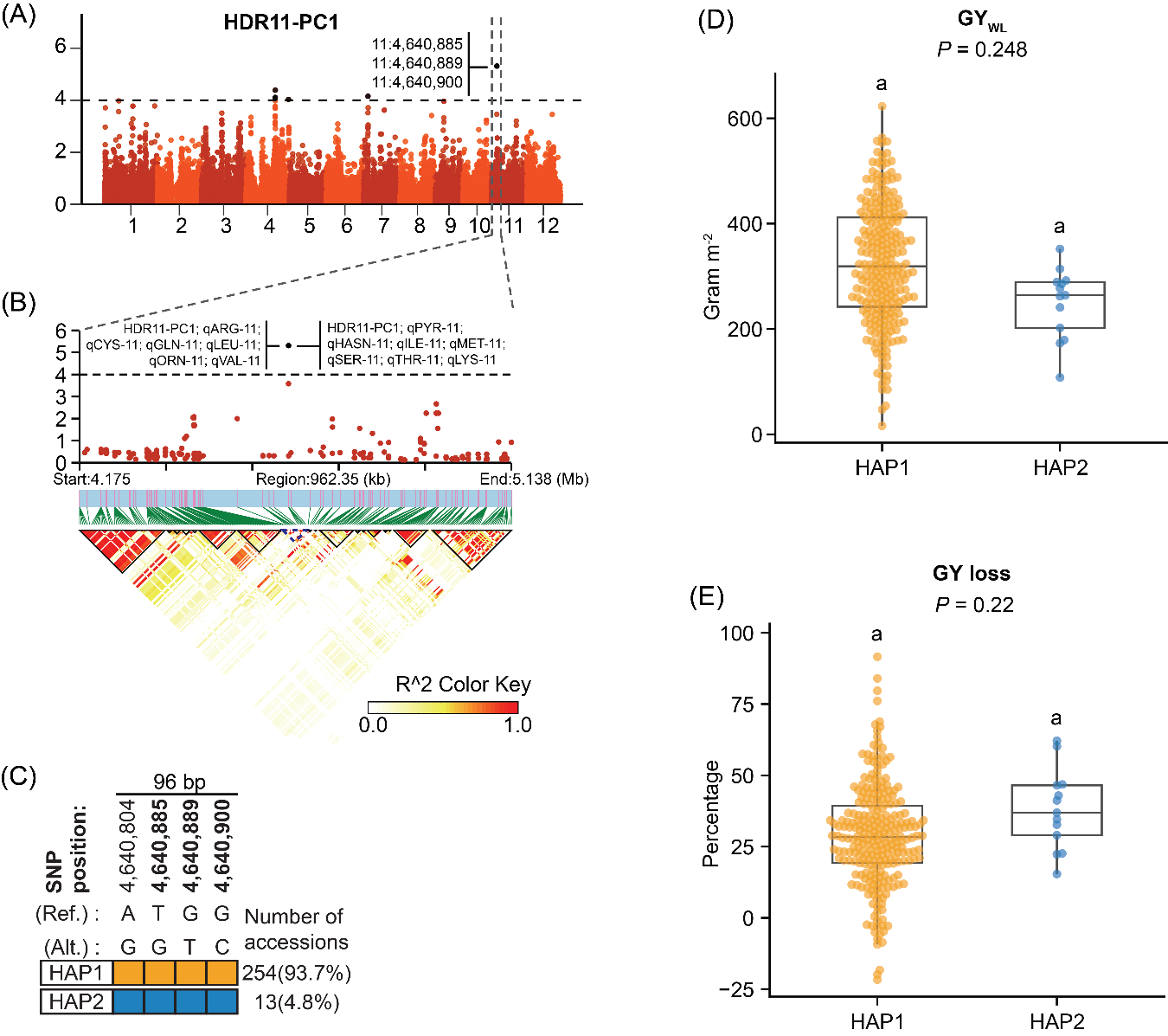

**Supplementary Fig. S7. Haplotype analysis, grain yield and grain yield loss associations at HDR11 on chromosome 11.**

(A) Manhattan plot of GWA mapping for the HDR11 PC1 metabolic proxy under water-limited (WL) conditions. The most significant SNPs (msSNPs) on chromosome 1 are indicated. The dashed line represents the significant threshold (-log_10_(*P*) = 4.0). (B) Regional association plot (upper) and pairwise LD heatmap (lower, *r^2^* colour key) for the HDR11 local LD interval (Chr 11: 4.175-5.138 Mb). Individual metabolite QTLs msSNPs are labelled. (C) Haplotype block structure at HDR11, comprising 4 SNPs spanning 96 bp; the msSNPs (4,640,885; 4,640,889 and 4,640,900) are in bold. (D) Grain yield (GY) comparisons between haplotypes under water-limited conditions. (E) GY loss comparison between haplotypes. Reference and alternative alleles are shown; gold and blue jitters denote alternative and reference allelic composition of HAP1 and HAP2, respectively. *P-*values are from Wald chi-square test; different letters denote significant differences based on Tukey's HSD pairwise contrasts (α = 0.05). Box plots show medians, interquartile ranges, and individual data points.

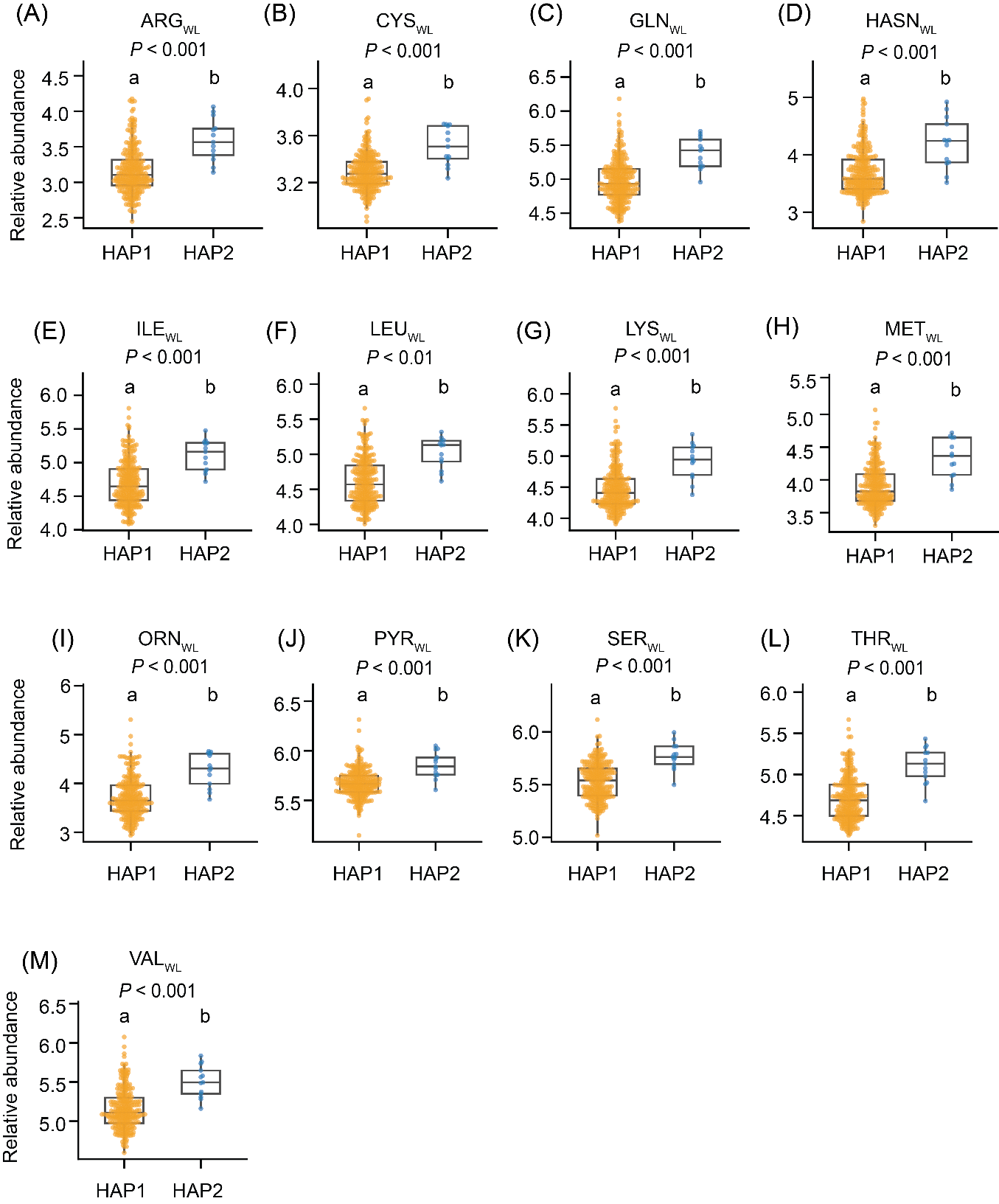

**Supplementary Fig. S8. Haplotype-trait associations at HDR11 loci on chromosome 11.**

(A) arginine, (B) cysteine, (C) glutamine, (D) 3-hydroxyasparagine, (E) isoleucine, (F) leucine, (G) lysine, (H) methionine, (I) ornithine, (J) pyroglutamic acid, (K) serine, (L) threonine and (M) valine, all under water-limited (WL) conditions for HAP1 (n = 254) and HAP2 (n = 13). *P-*values are from Wald chi-square test; different letters denote significant differences based on Tukey's HSD pairwise contrasts (α = 0.05). Box plots show medians, interquartile ranges, and individual data points; colours correspond to HAP1 (gold) and HAP2 (blue).

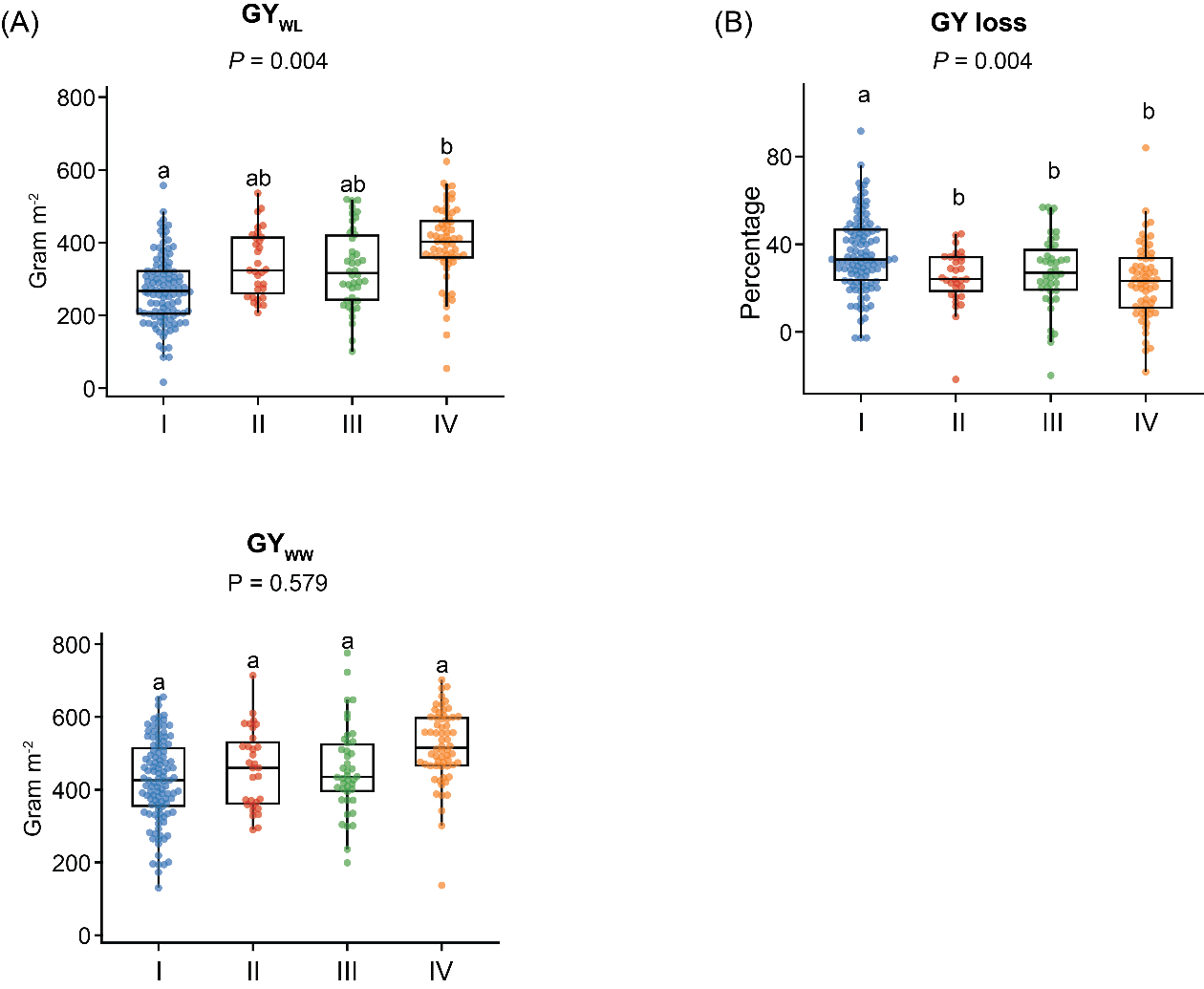

**Supplementary Fig. S9. Effect of multi-locus allele combinations at HDR1-2 and HDR4 on grain yield under well-watered (GY_WW_) conditions.**

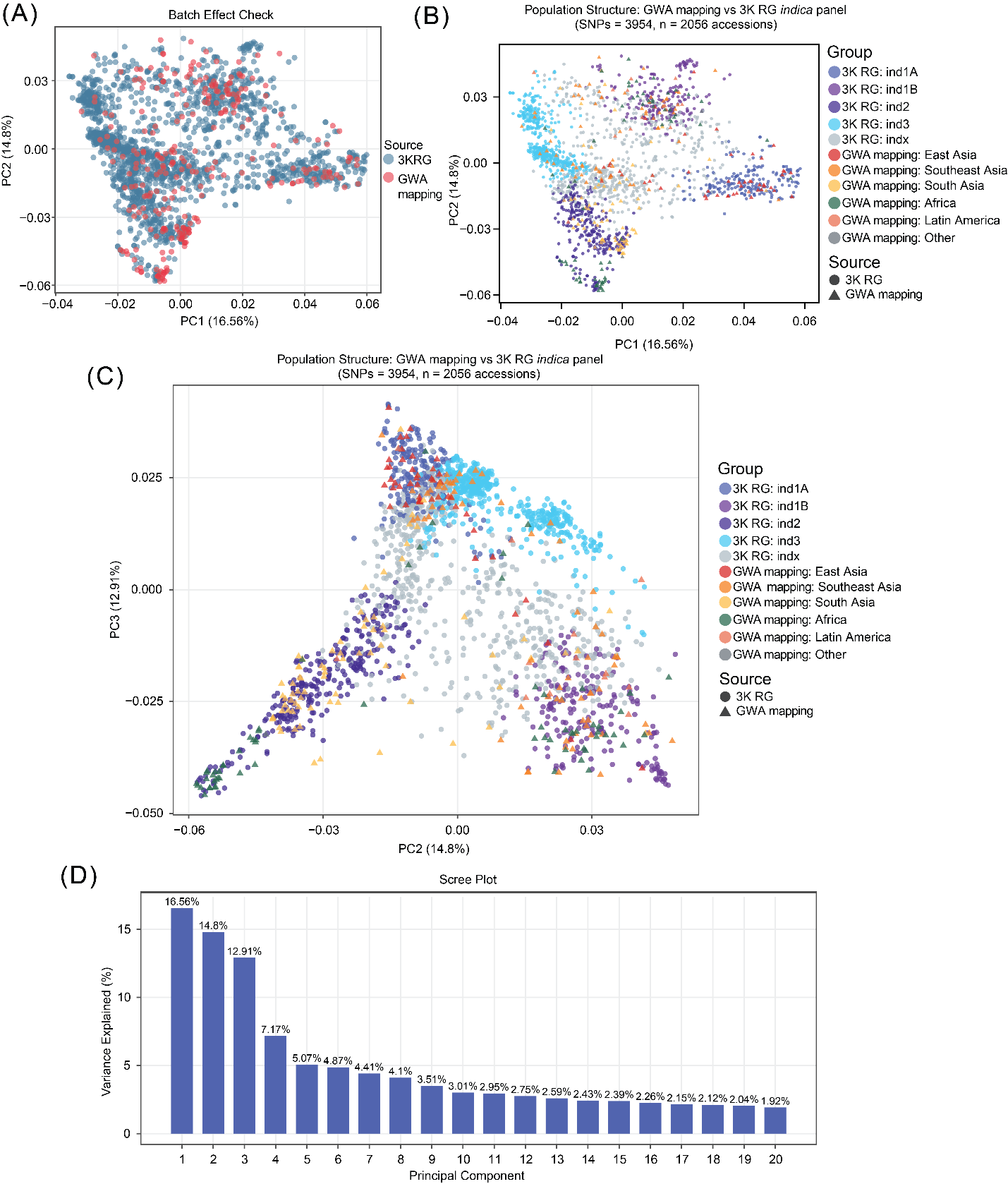

**Supplementary Fig. 10. Population structure of the 3KRG *indica* panel and the GWA mapping *indica* panel**

(A) Top 500 closest 3KRG indica accessions to the GWAS mapping panel. Accessions are coloured based on sources and mean distance score. (B) PCA of a merged dataset of 271 GWA mapping *indica* panel accessions and 1,785 3K RGP *indica* accessions (n = 2,056; 3,954 LD-pruned SNPs). PC1 versus PC2 is shown. Accessions are coloured by *indica* subpopulation (3K RGP: ind1A, ind1B, ind2, ind3, indx) or geographic region (GWA mapping *indica* panel) and shaped by data source (circle, 3K RGP; triangle, GWA mapping panel). (C) Population structure of the 3KRG *indica* panel and the GWA mapping panel, PC2 versus PC3, is shown. Accessions as coloured and shaped as in Fig.7 B. (C) Scree plot of the first 20 principal components (PCs) of the PCA of the 3KRG *indica* panel and GWA mapping panel.

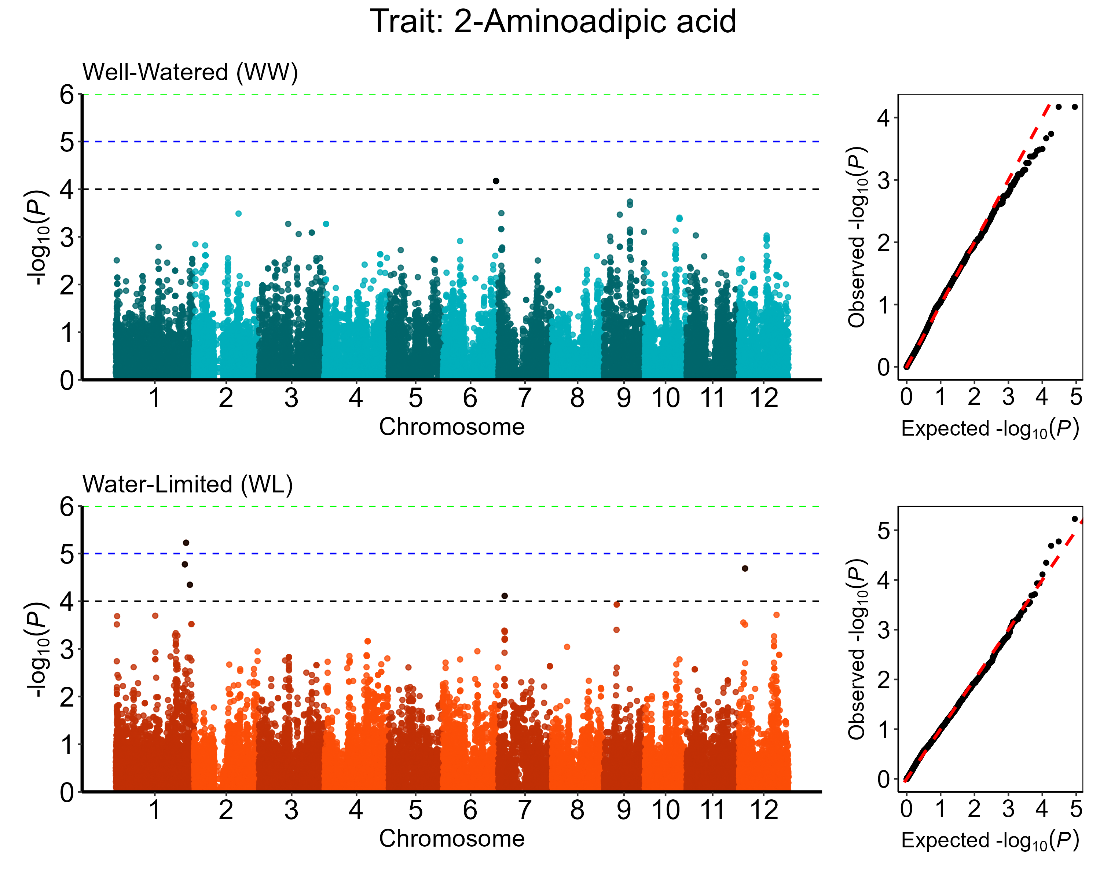

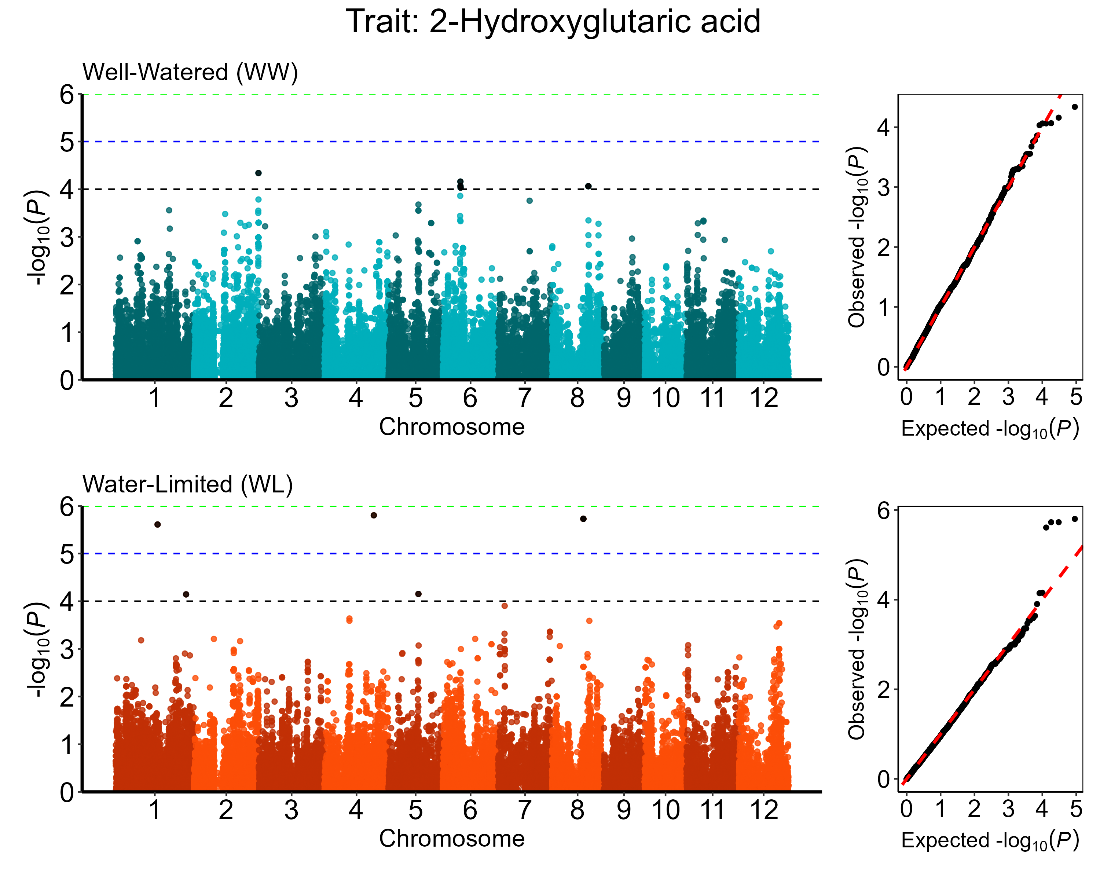

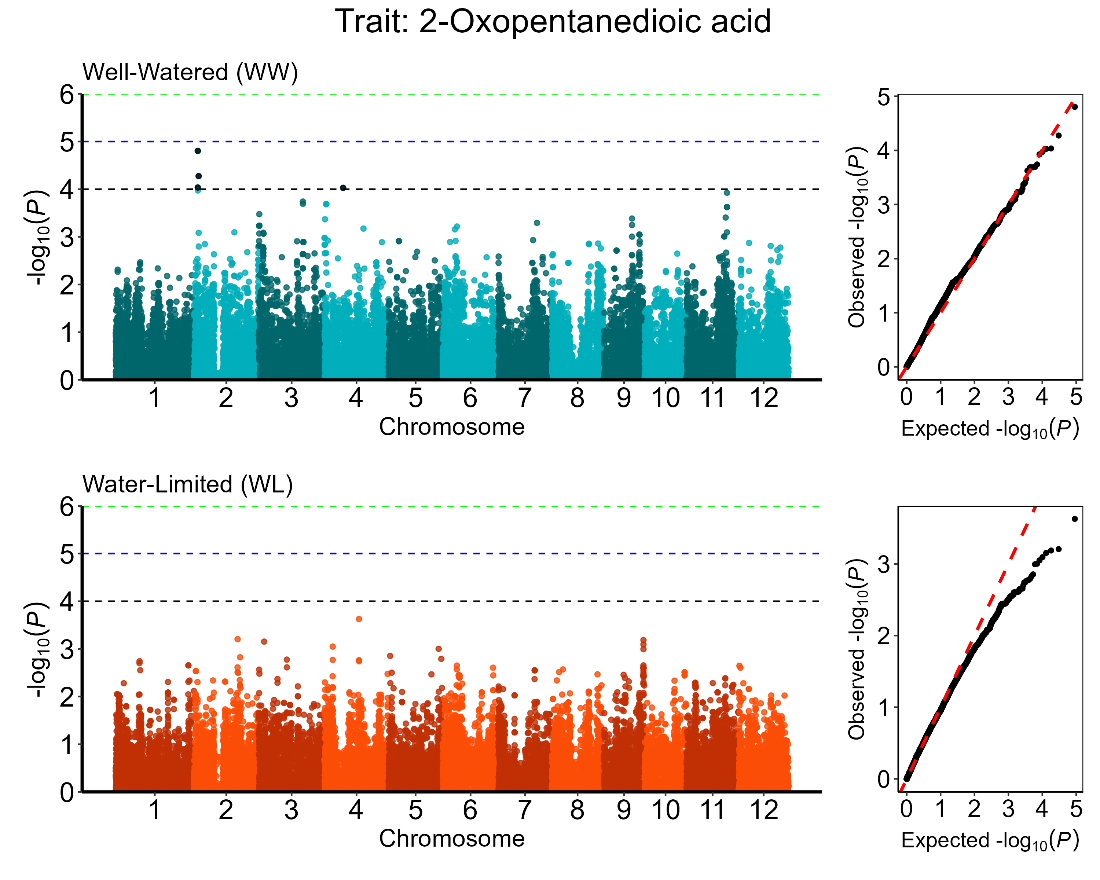

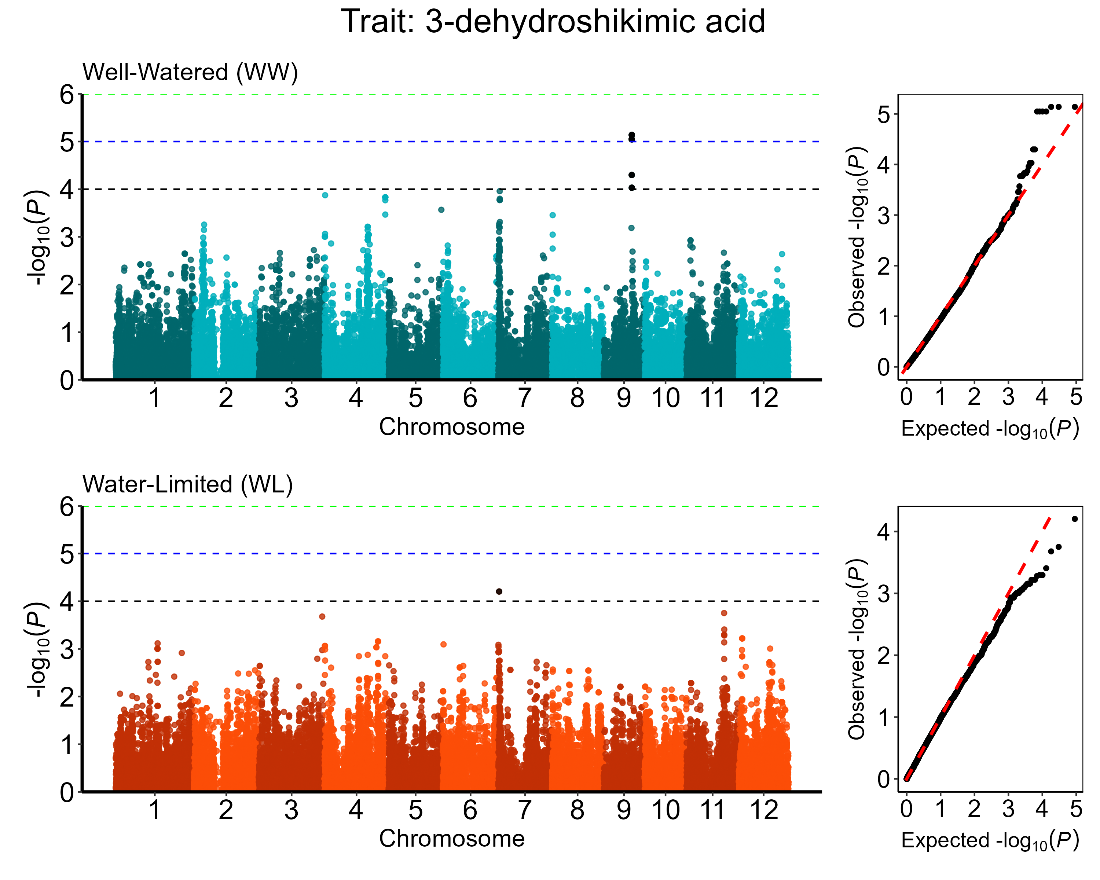

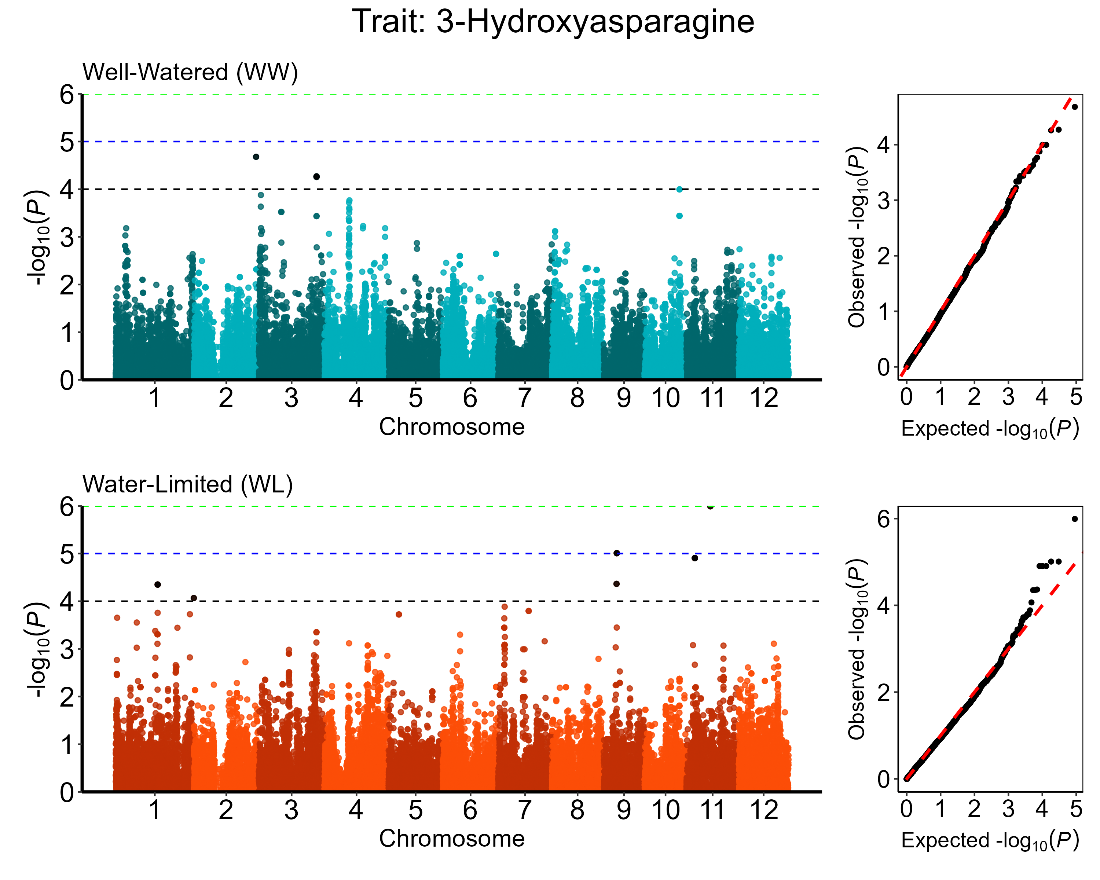

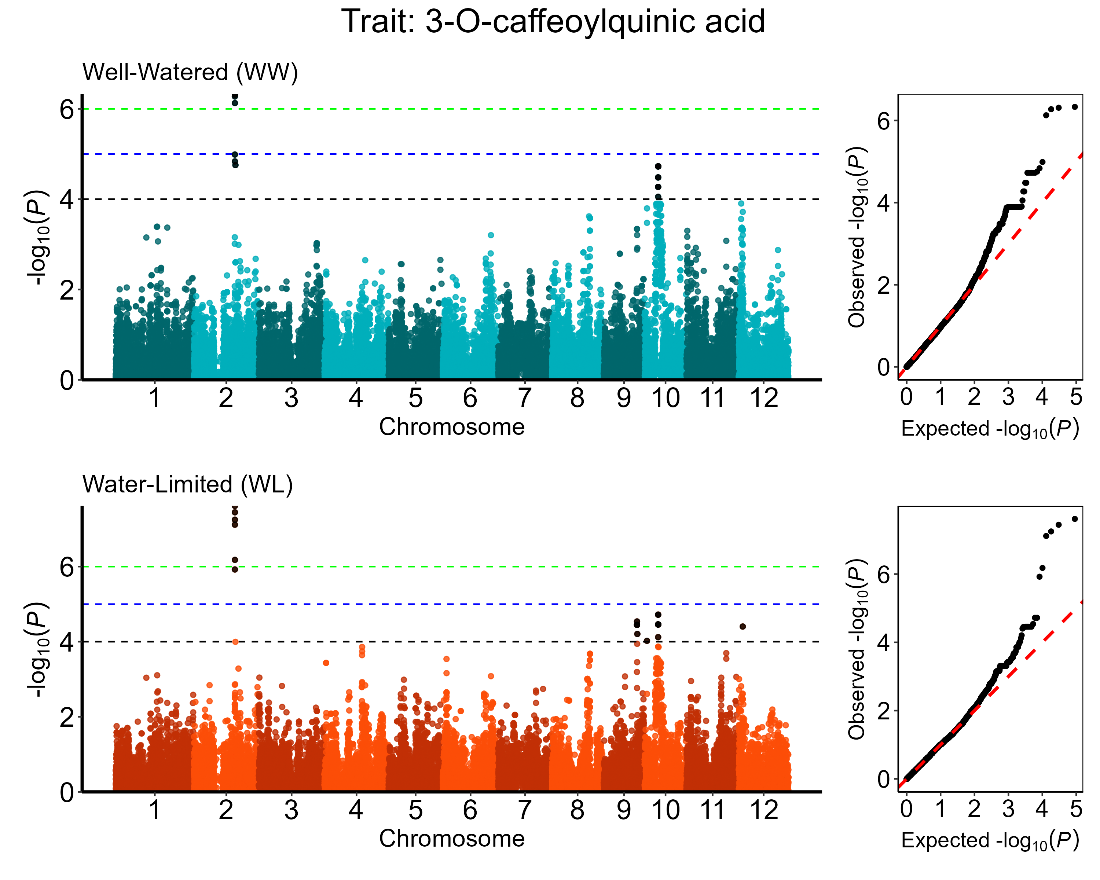

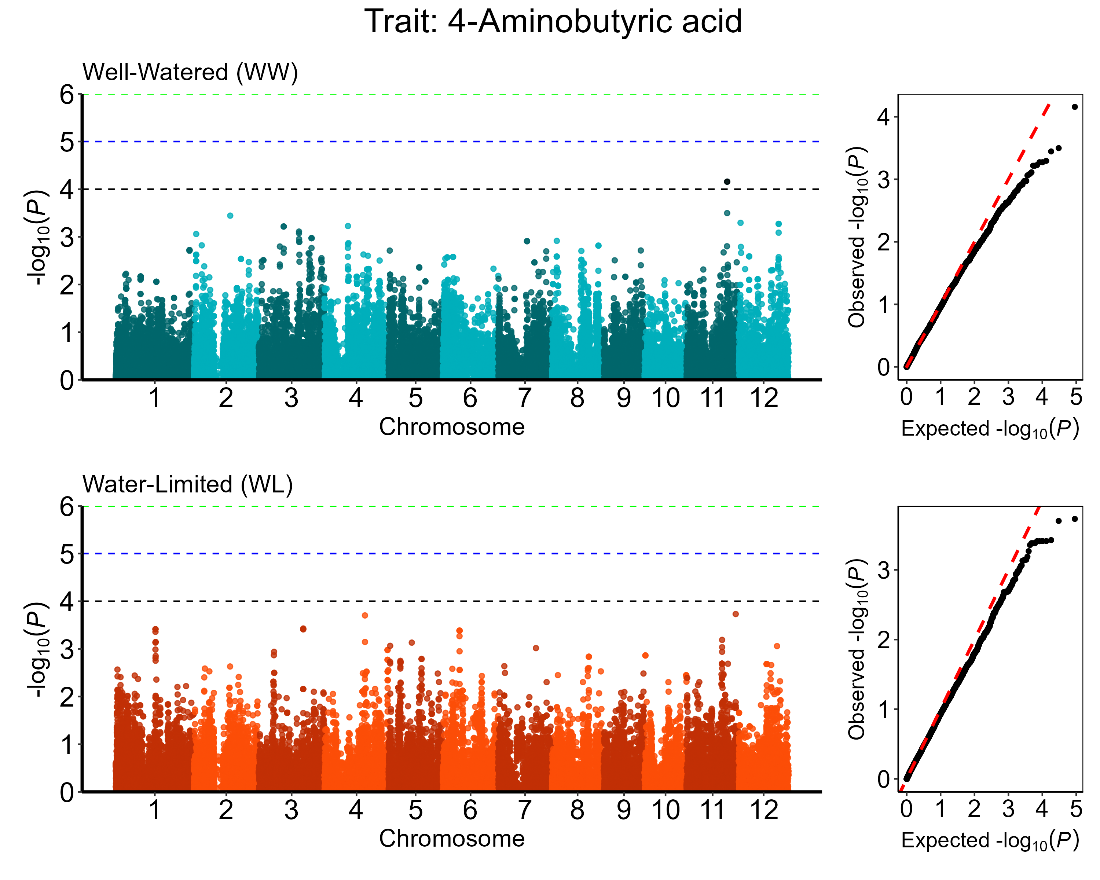

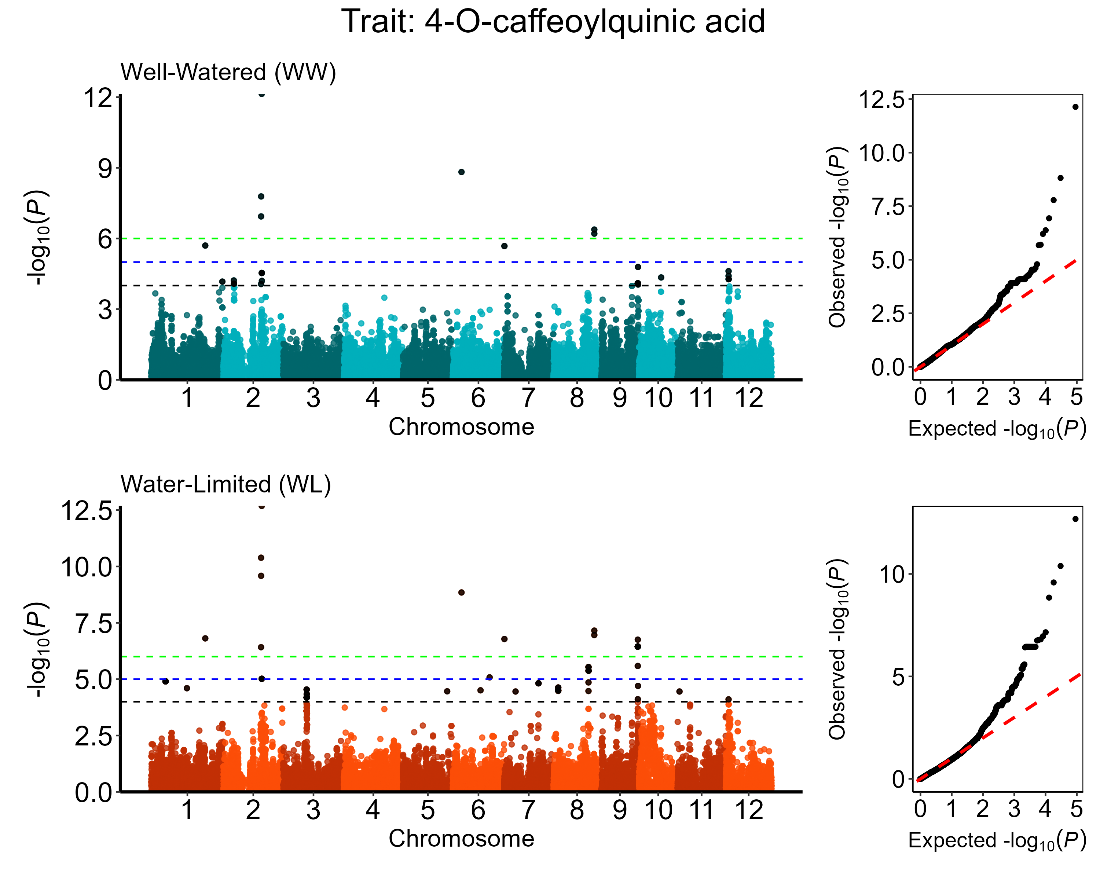

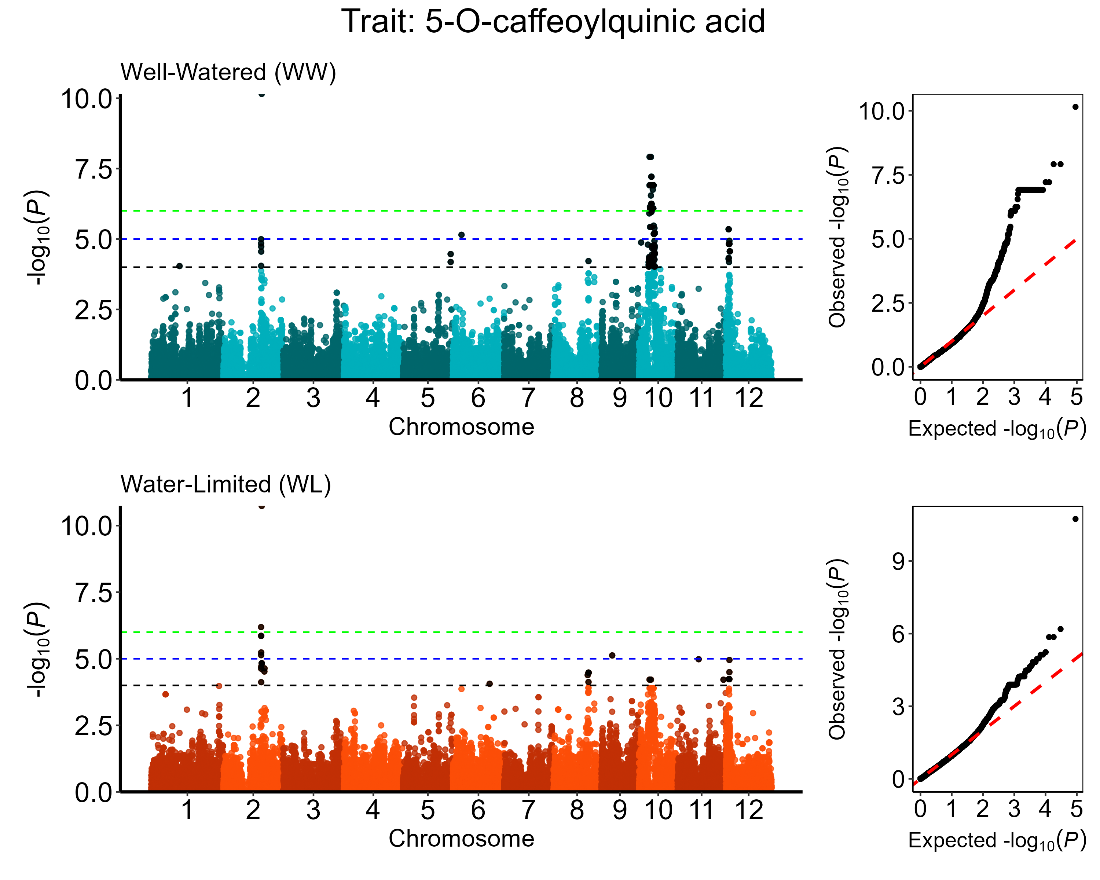

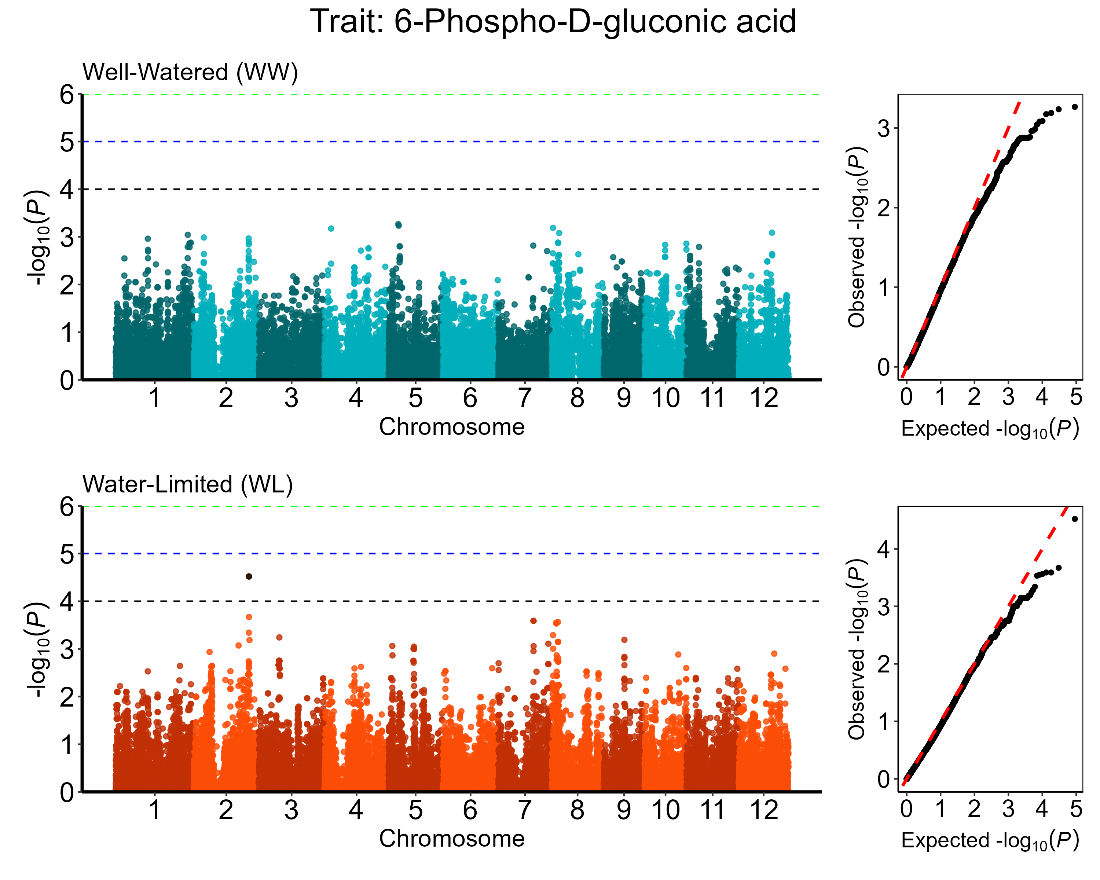

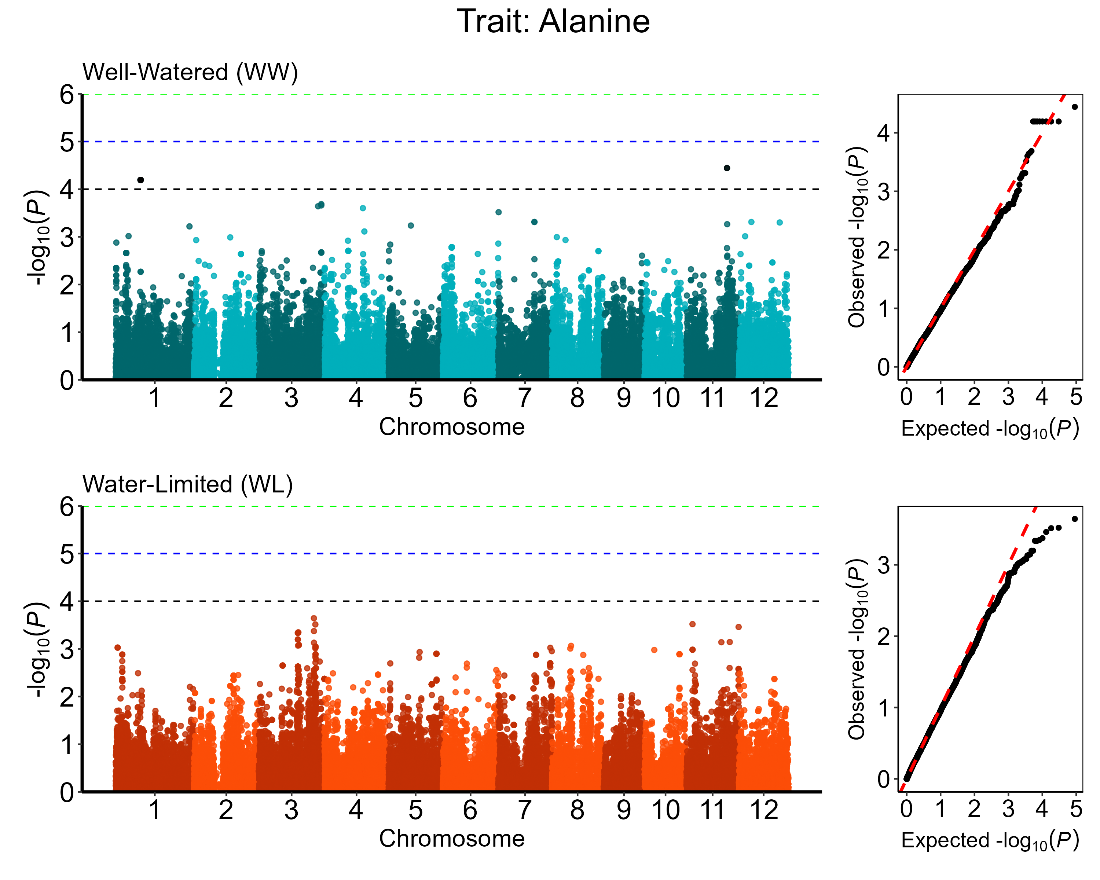

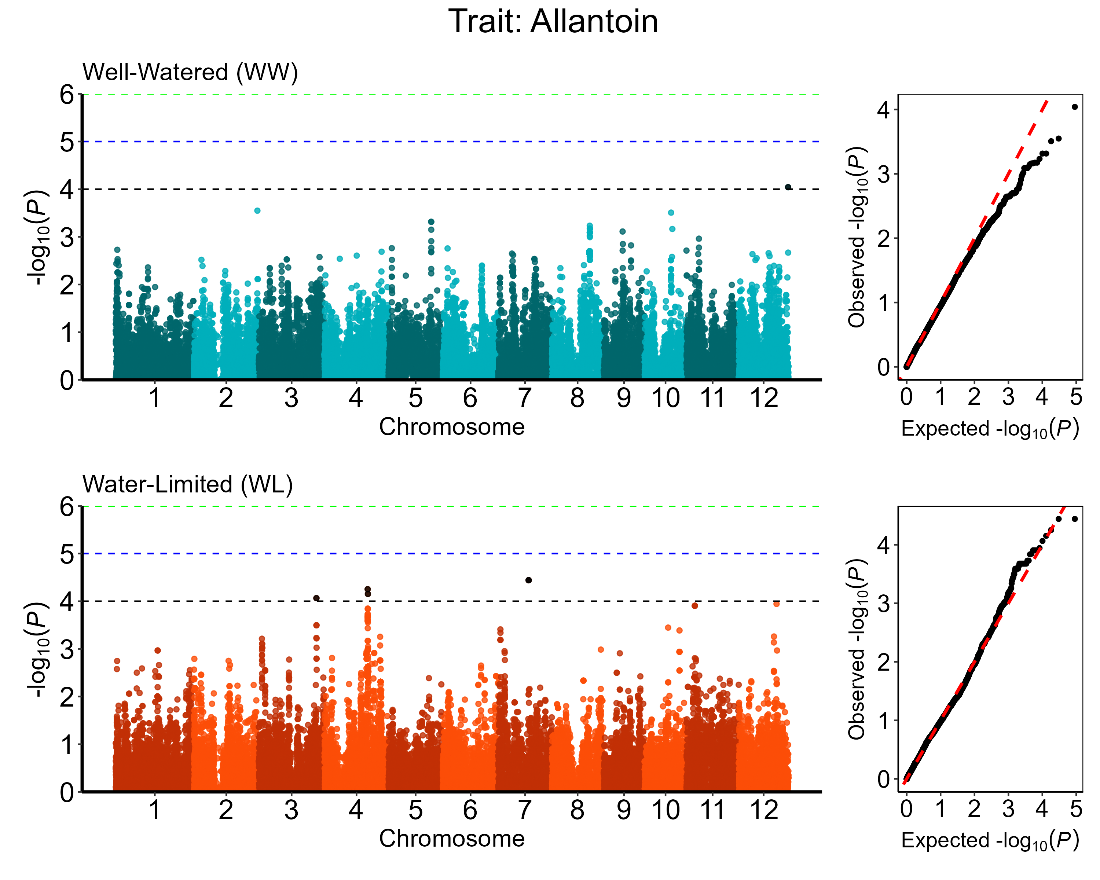

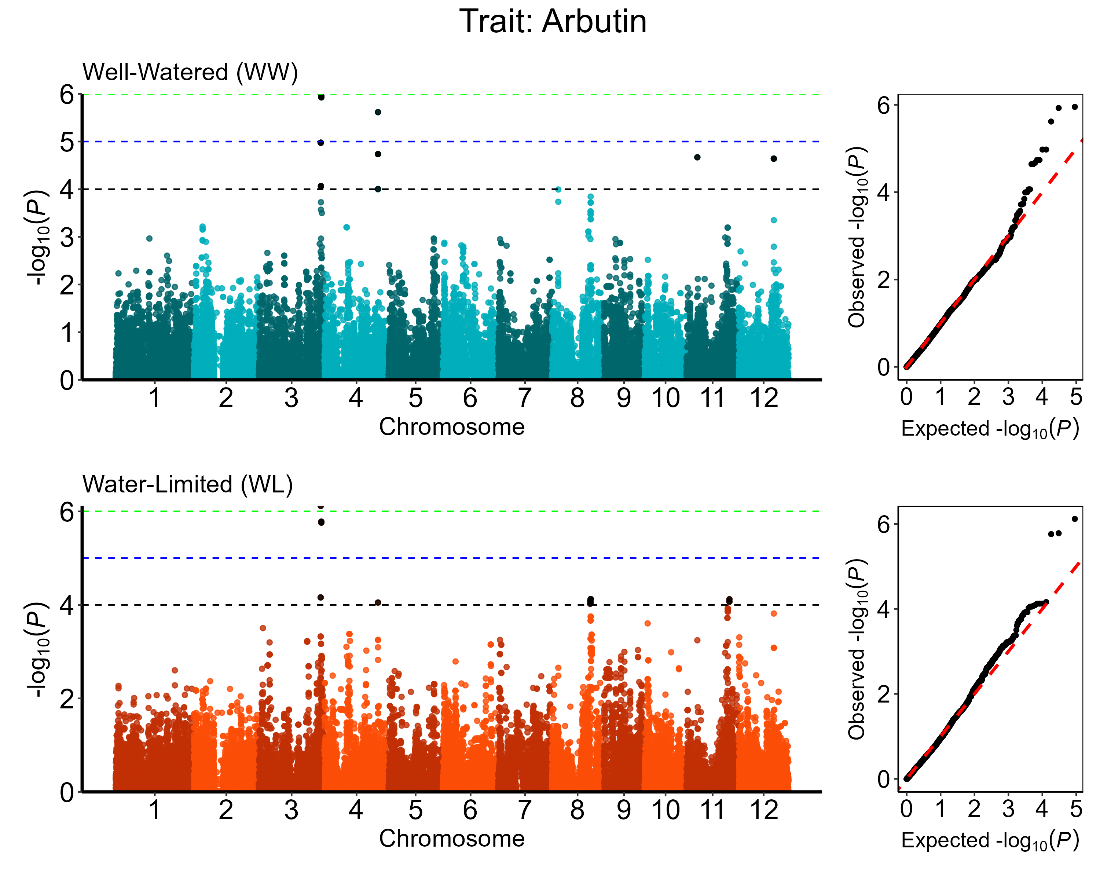

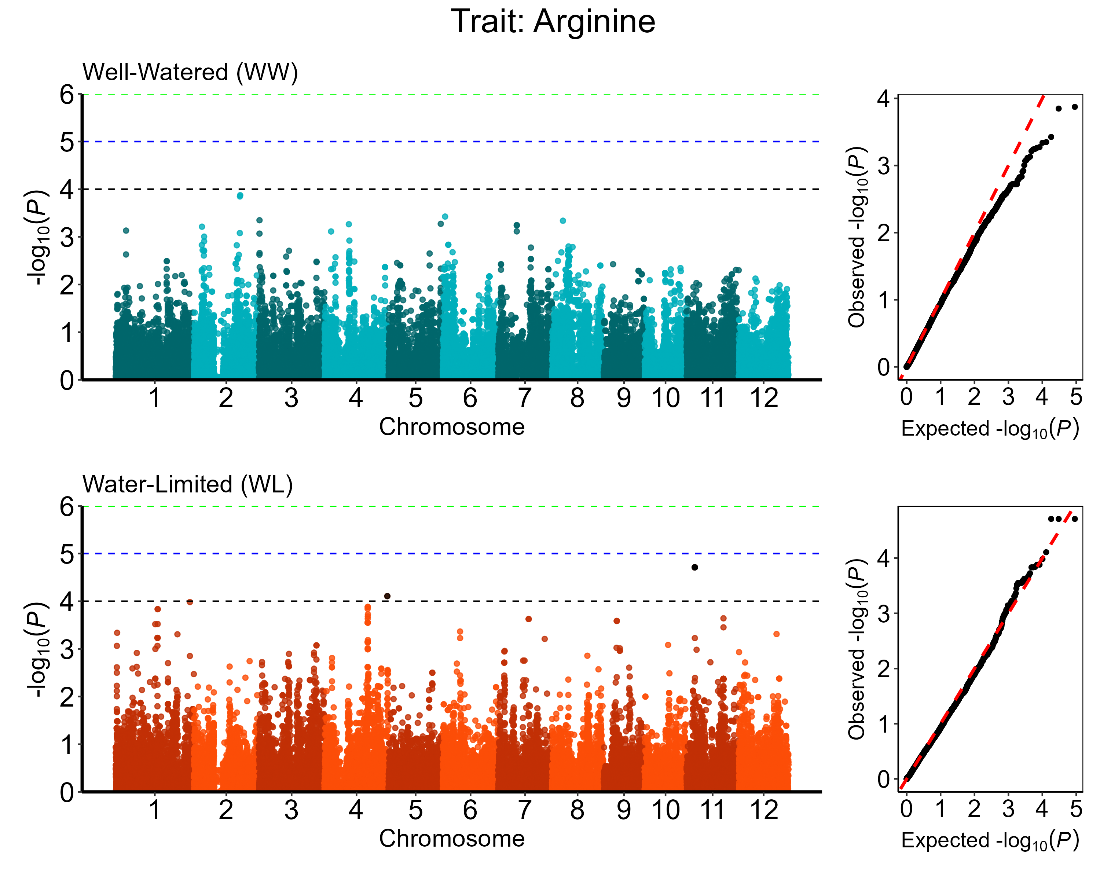

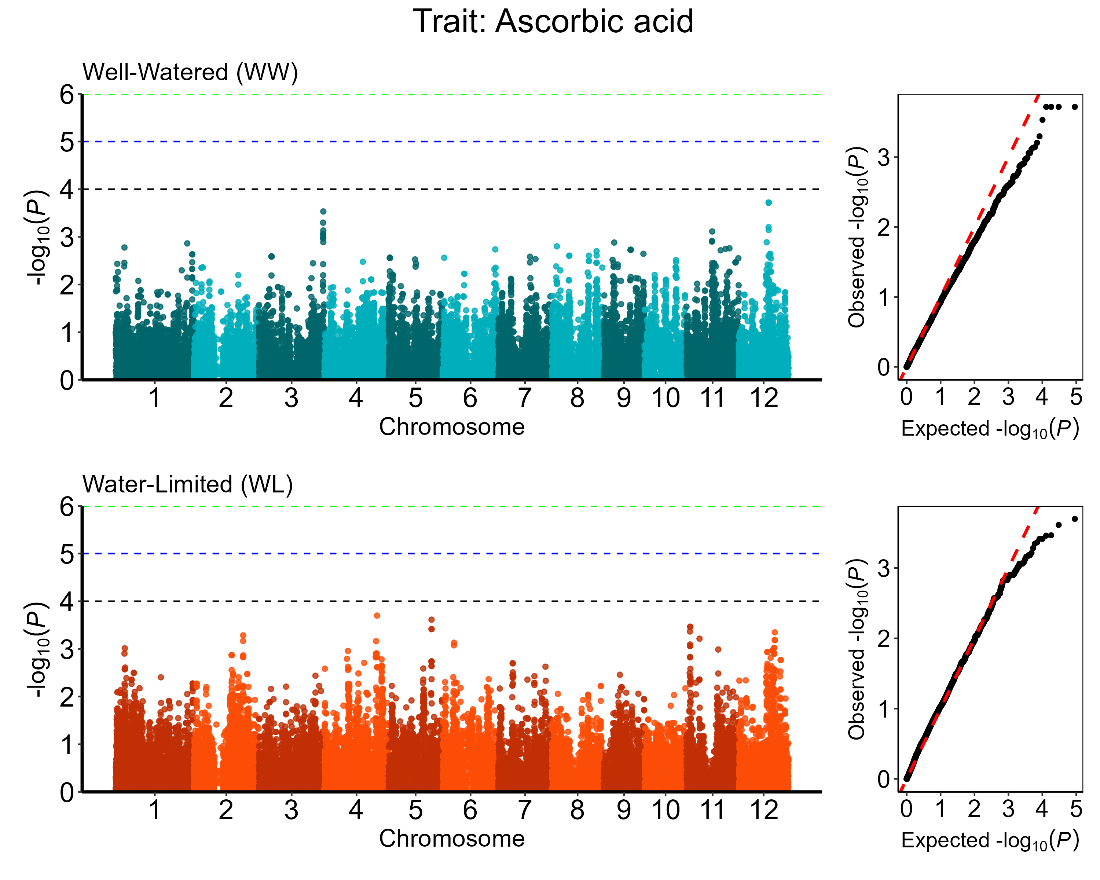

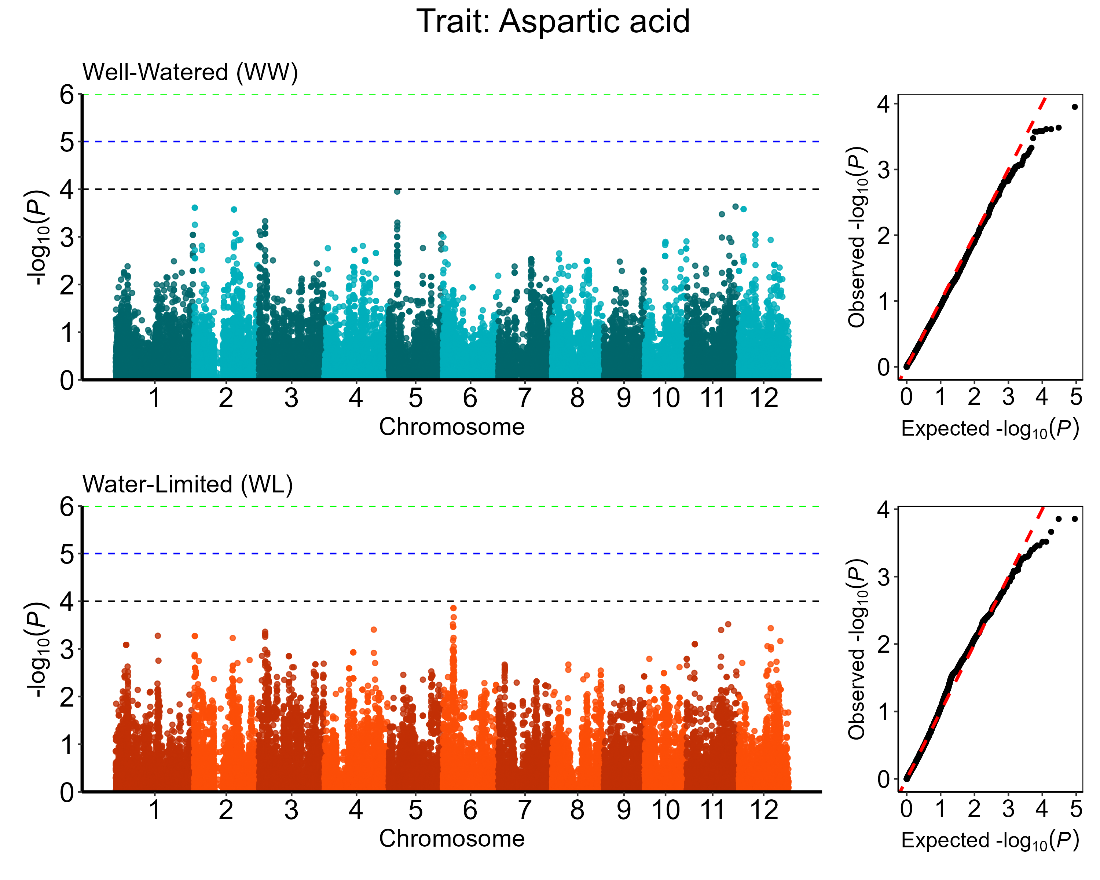

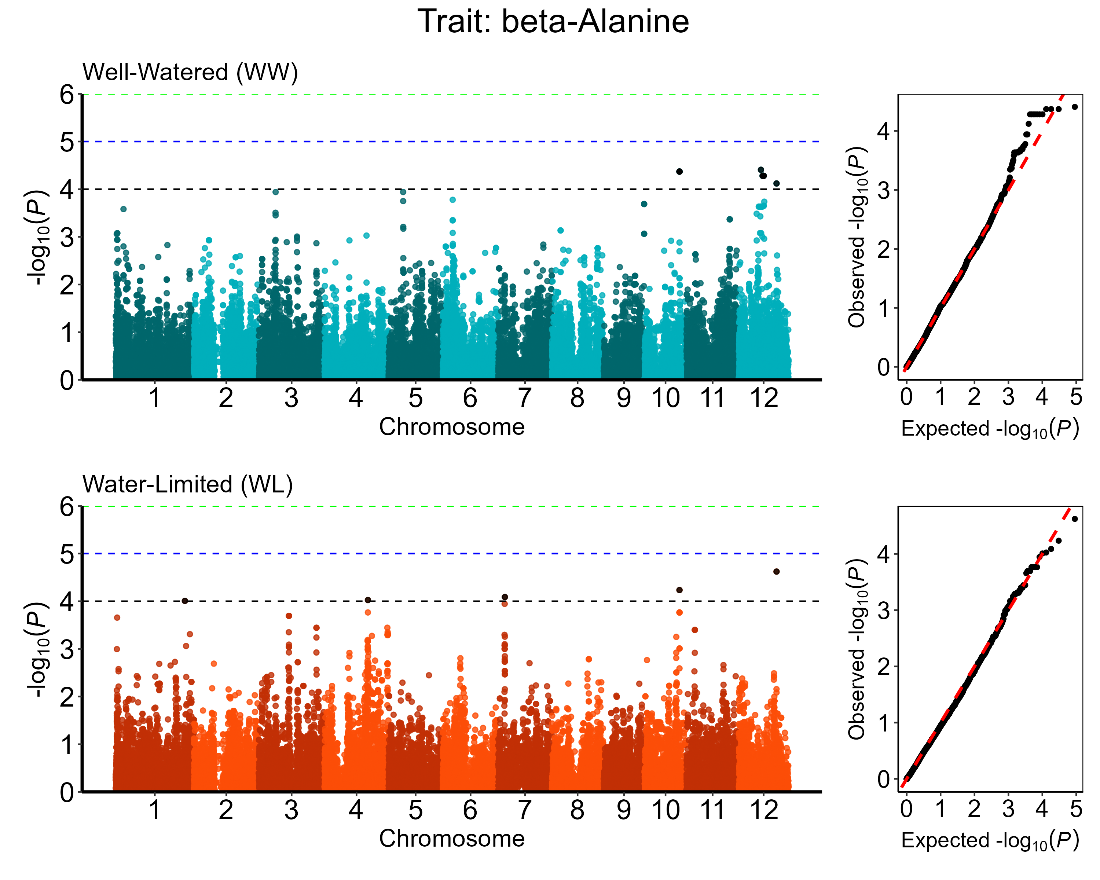

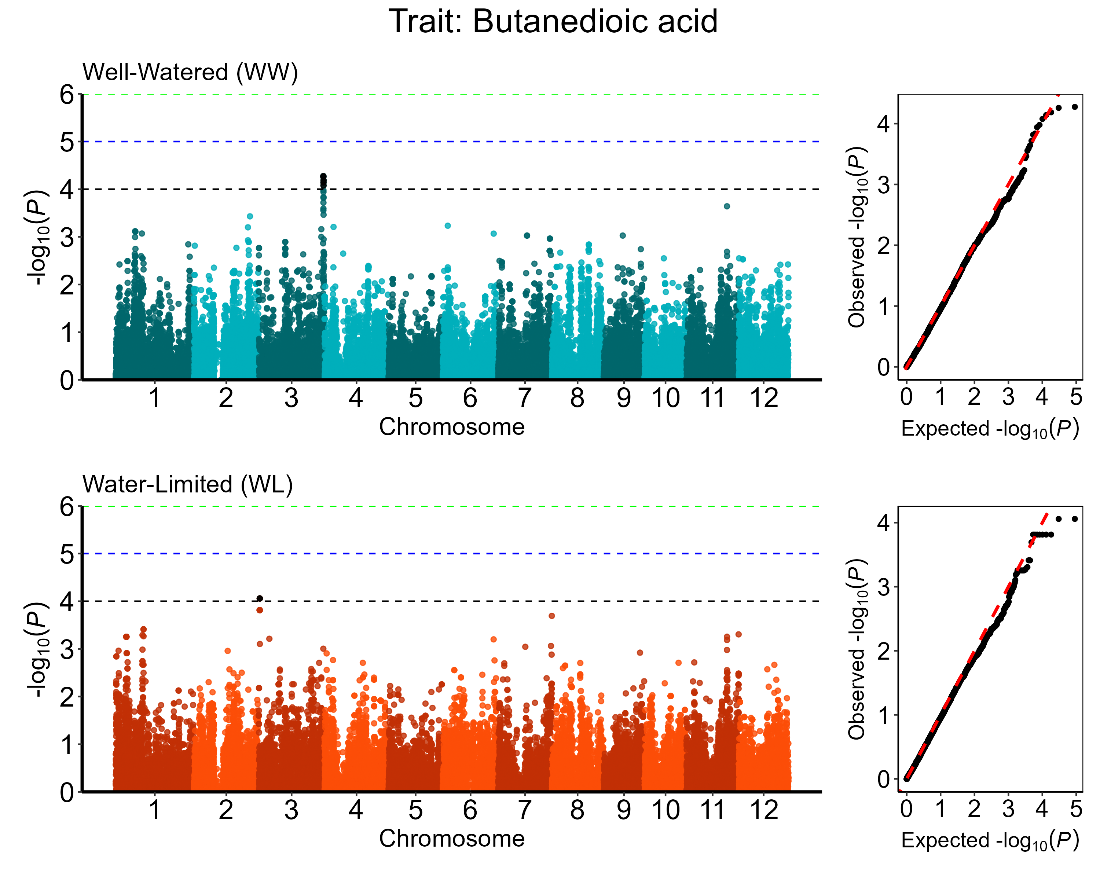

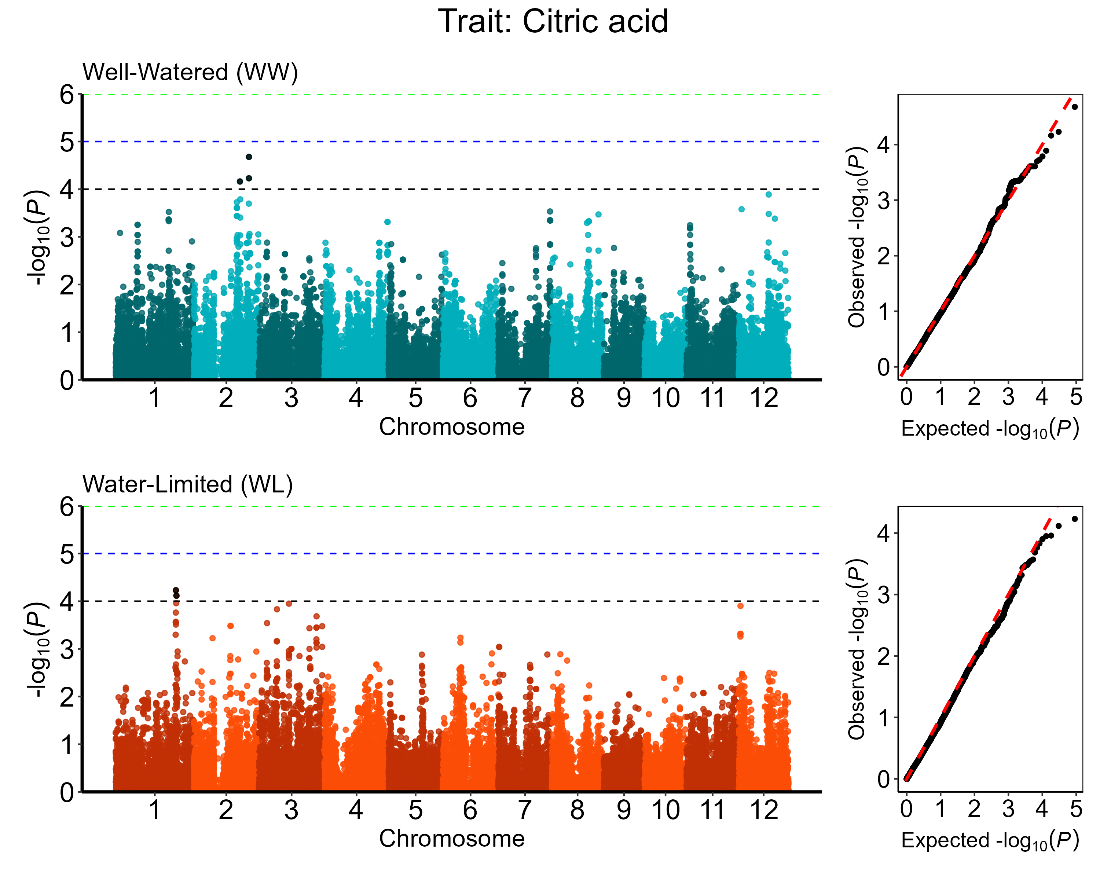

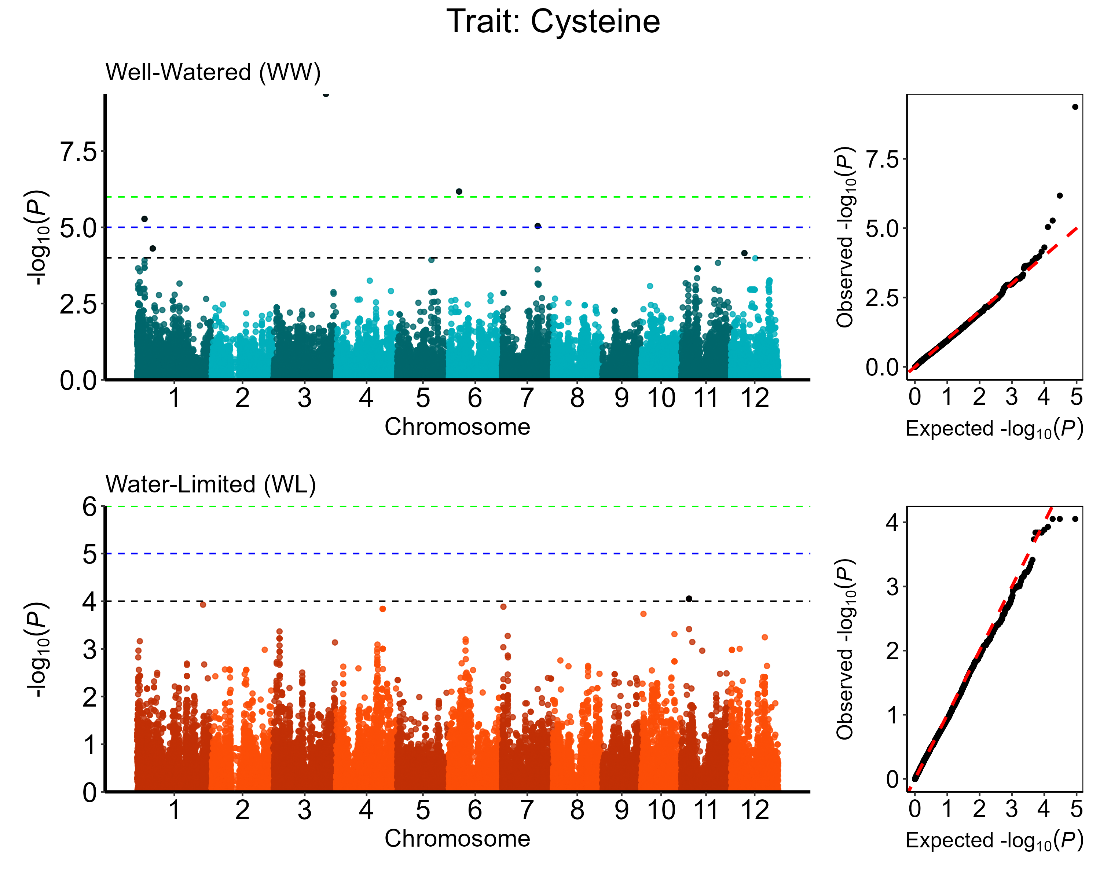

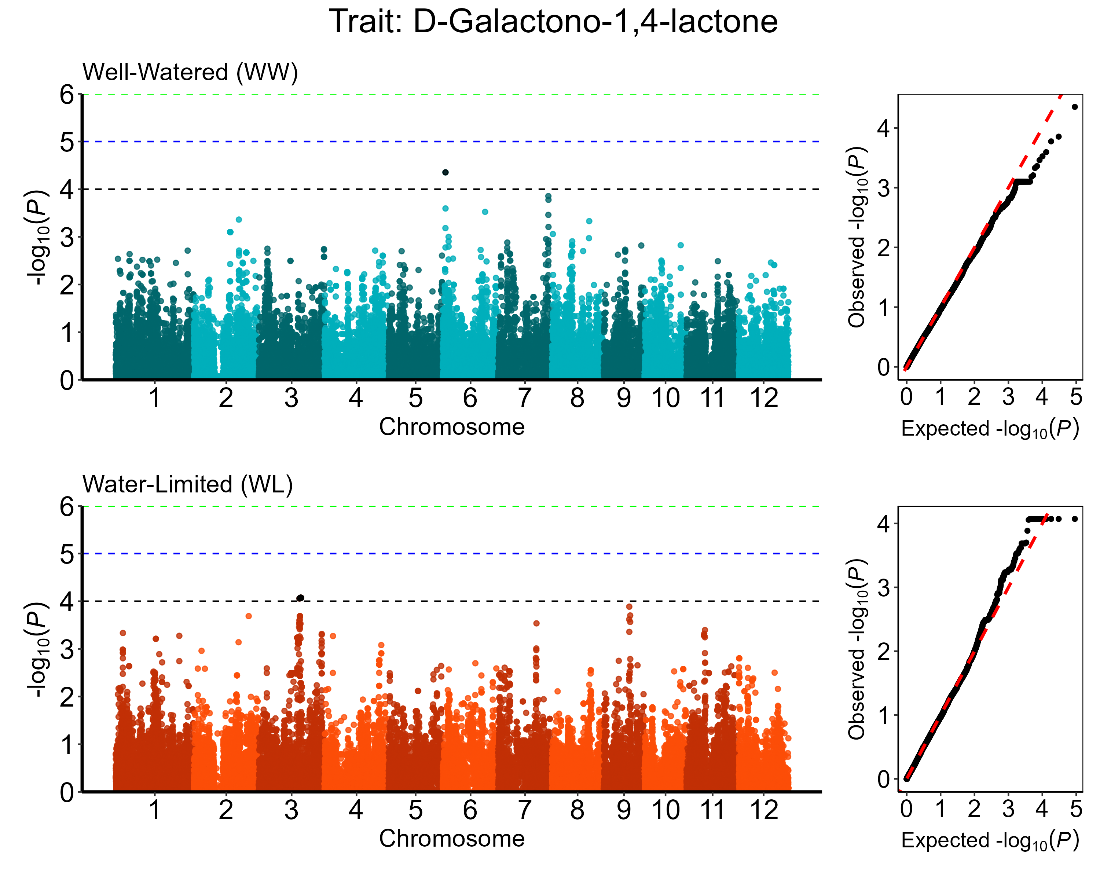

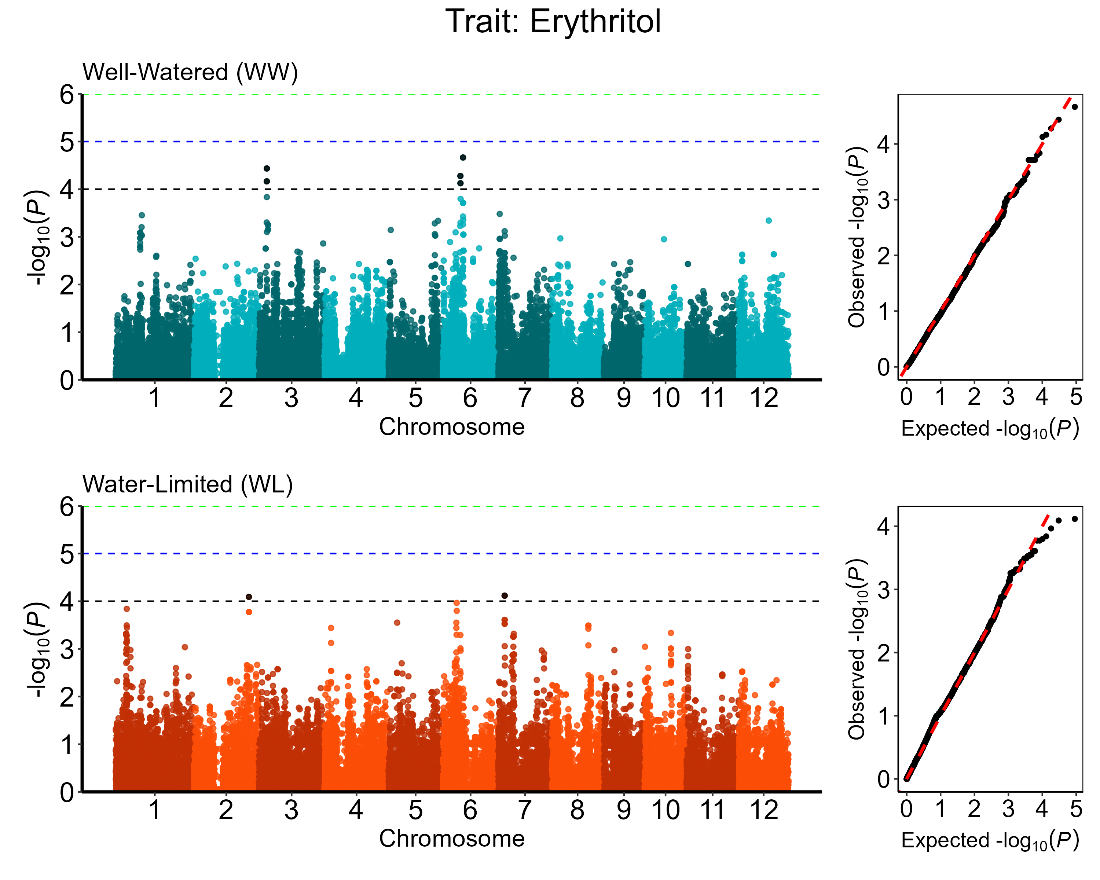

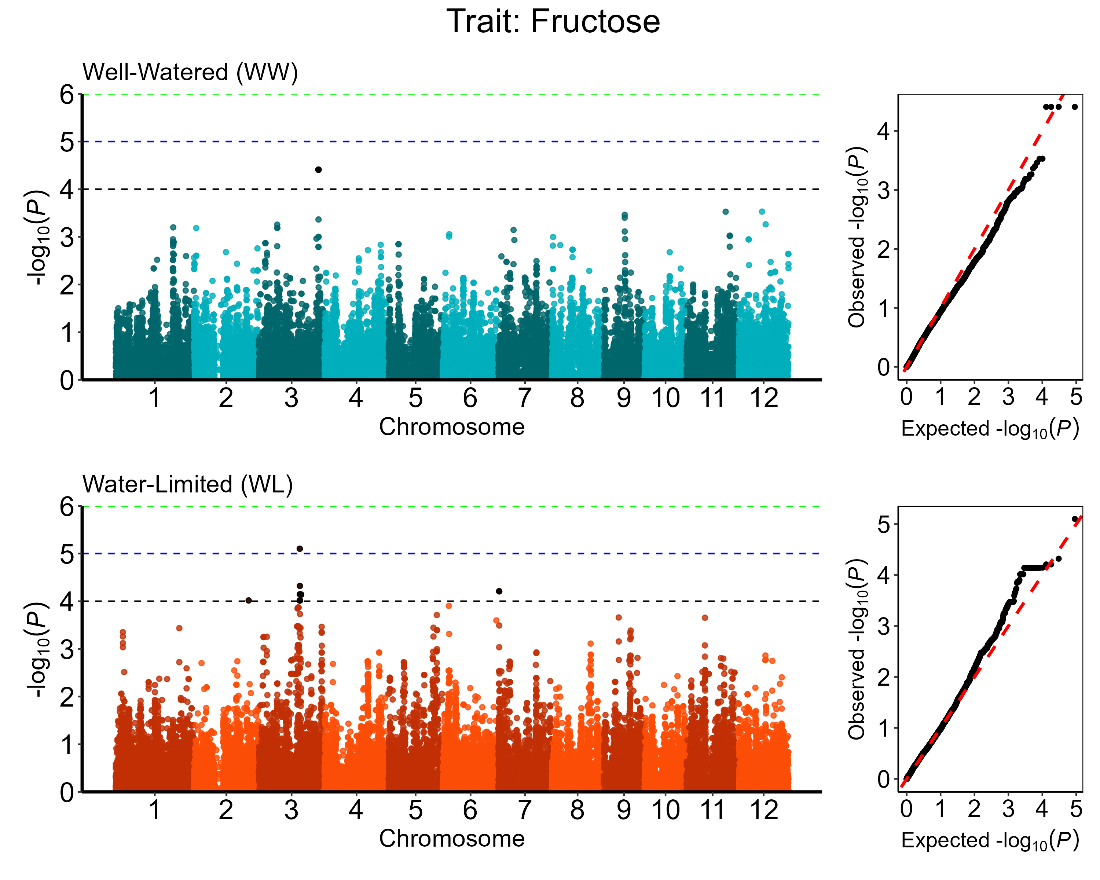

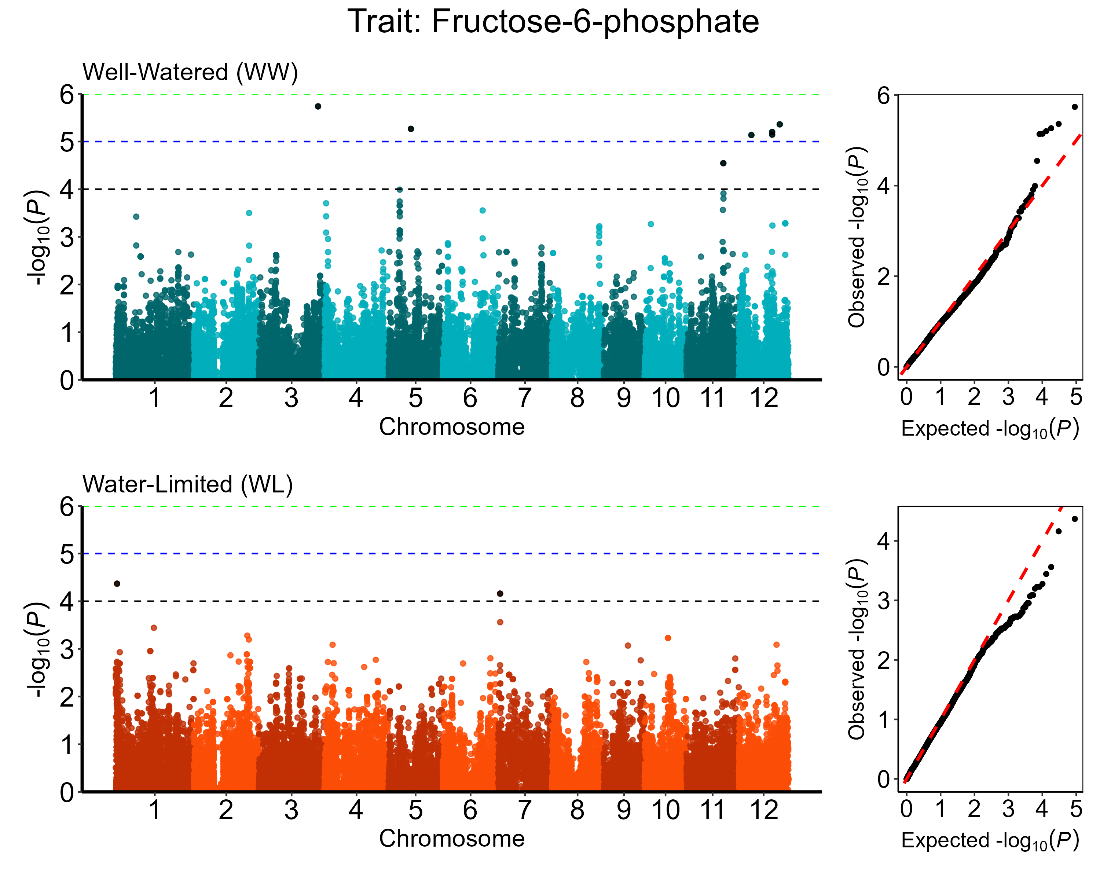

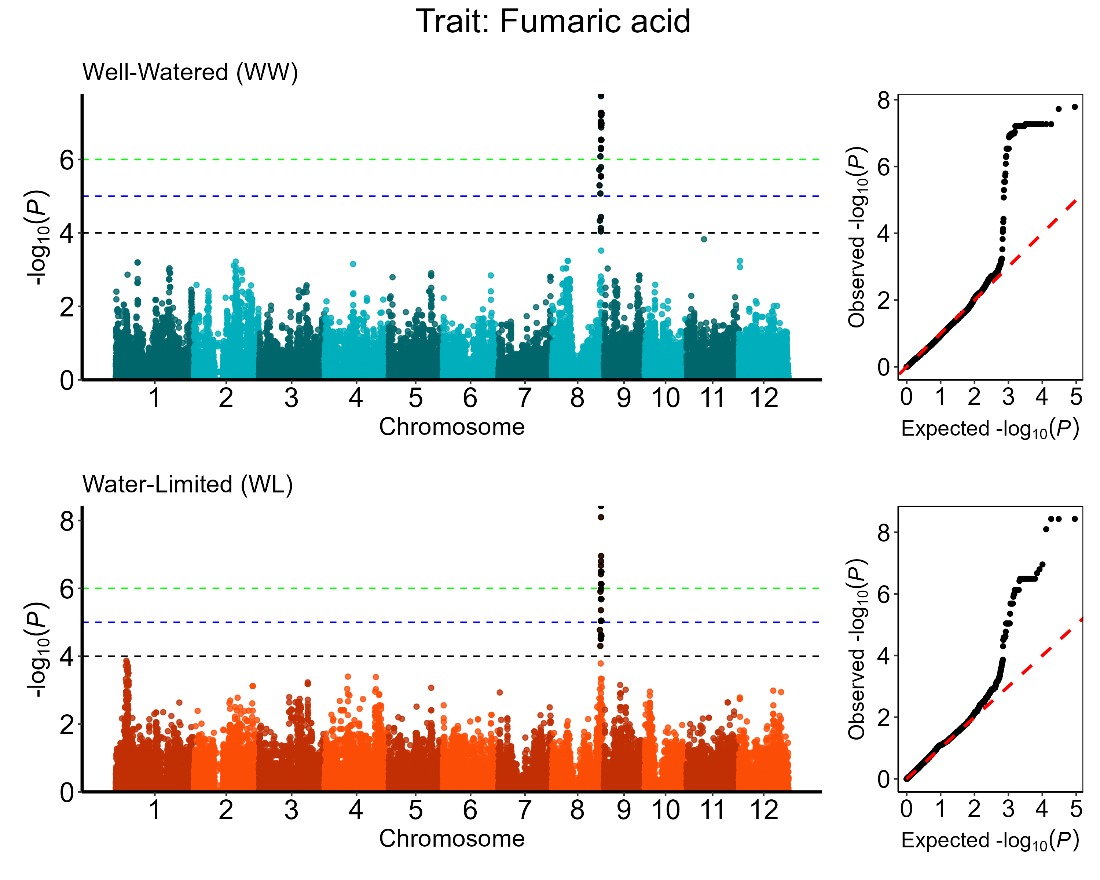

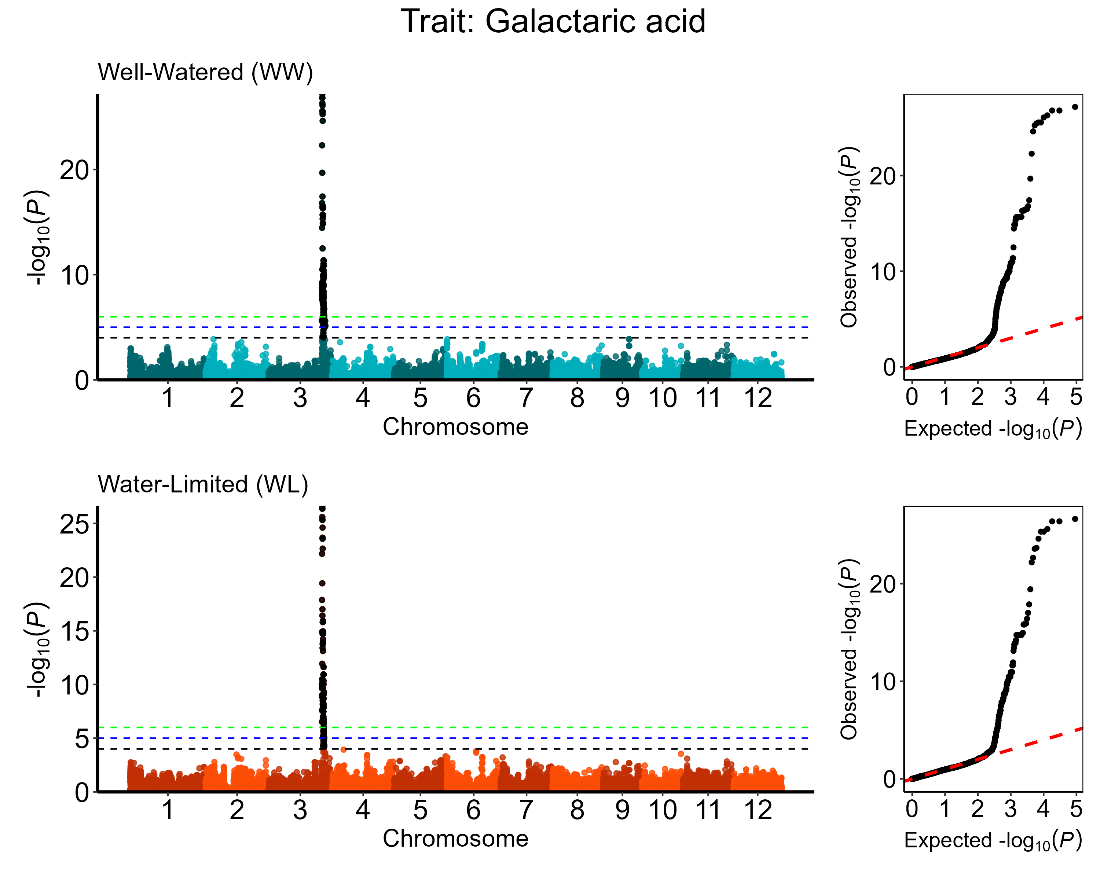

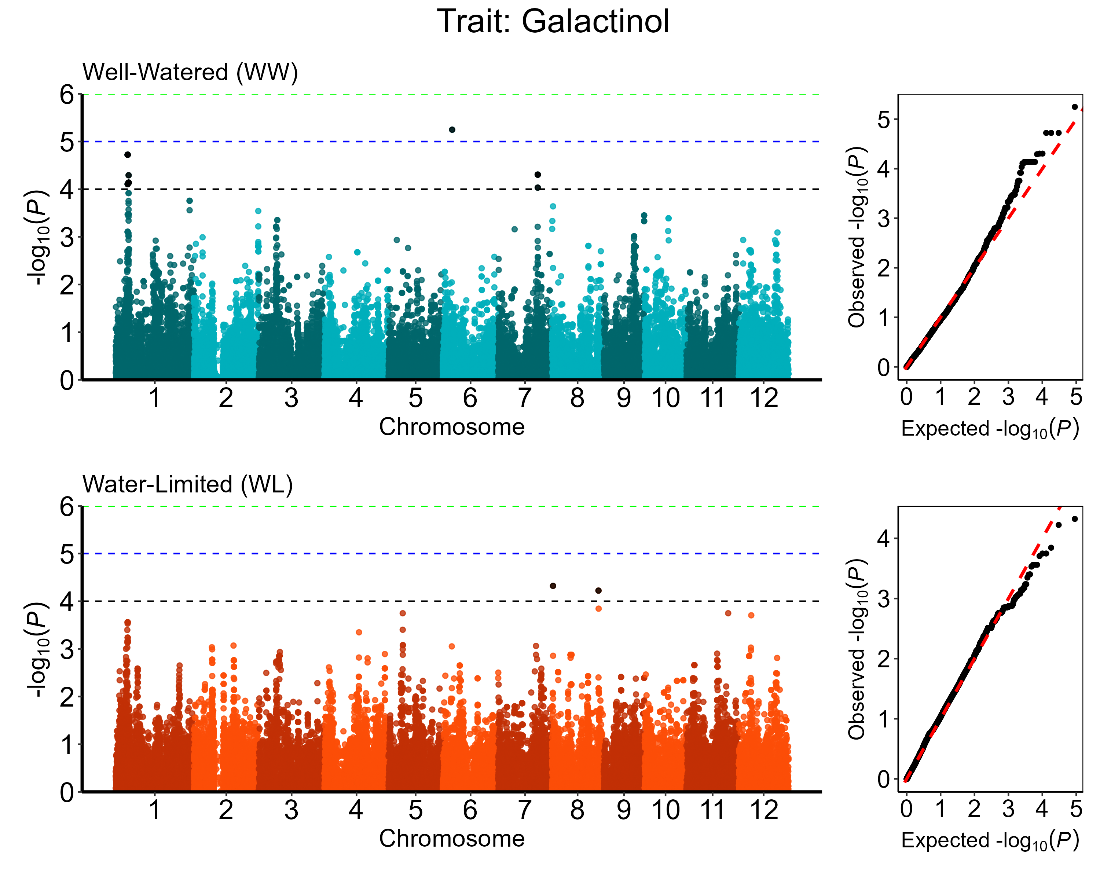

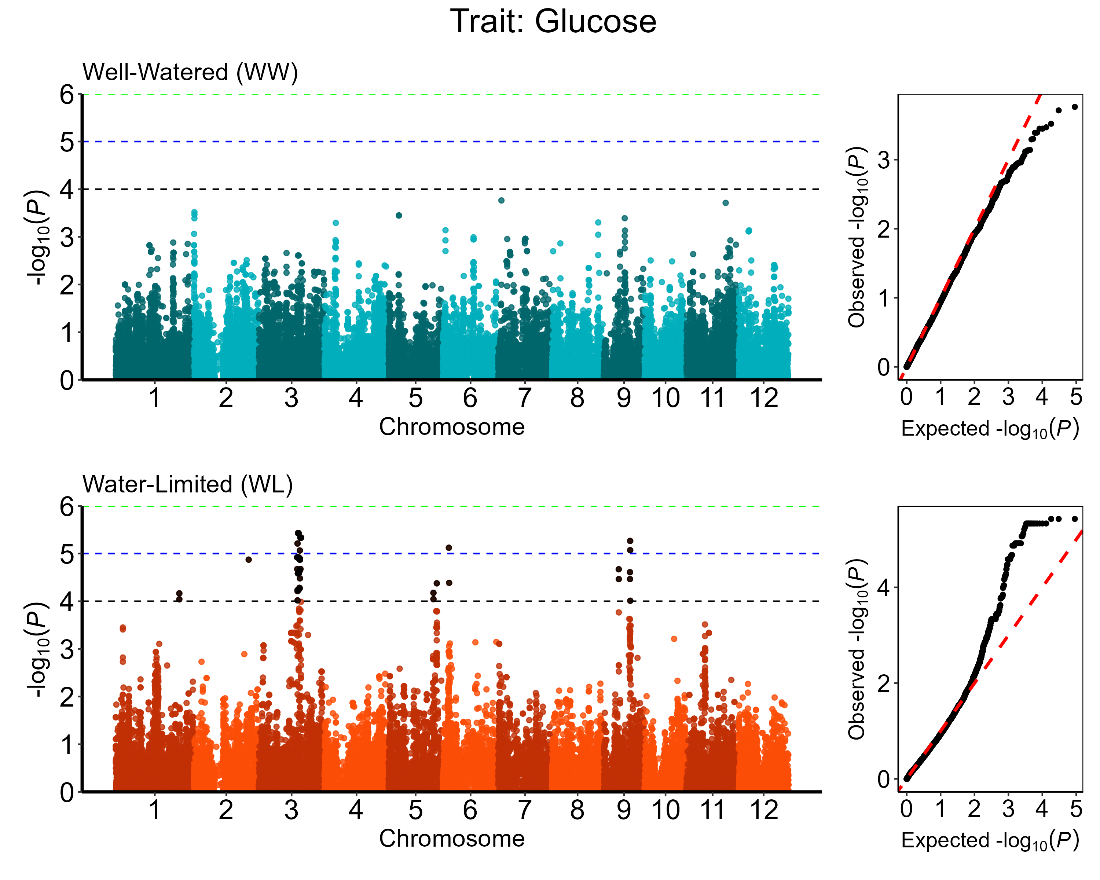

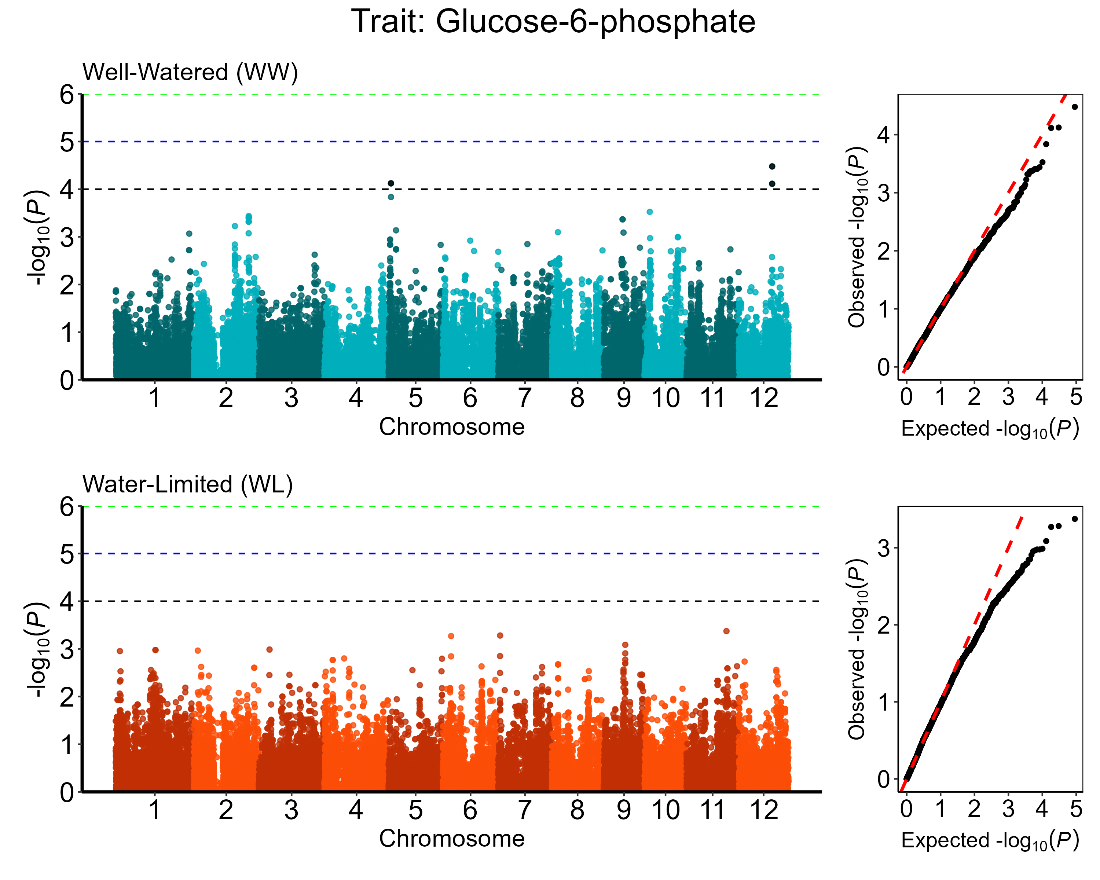

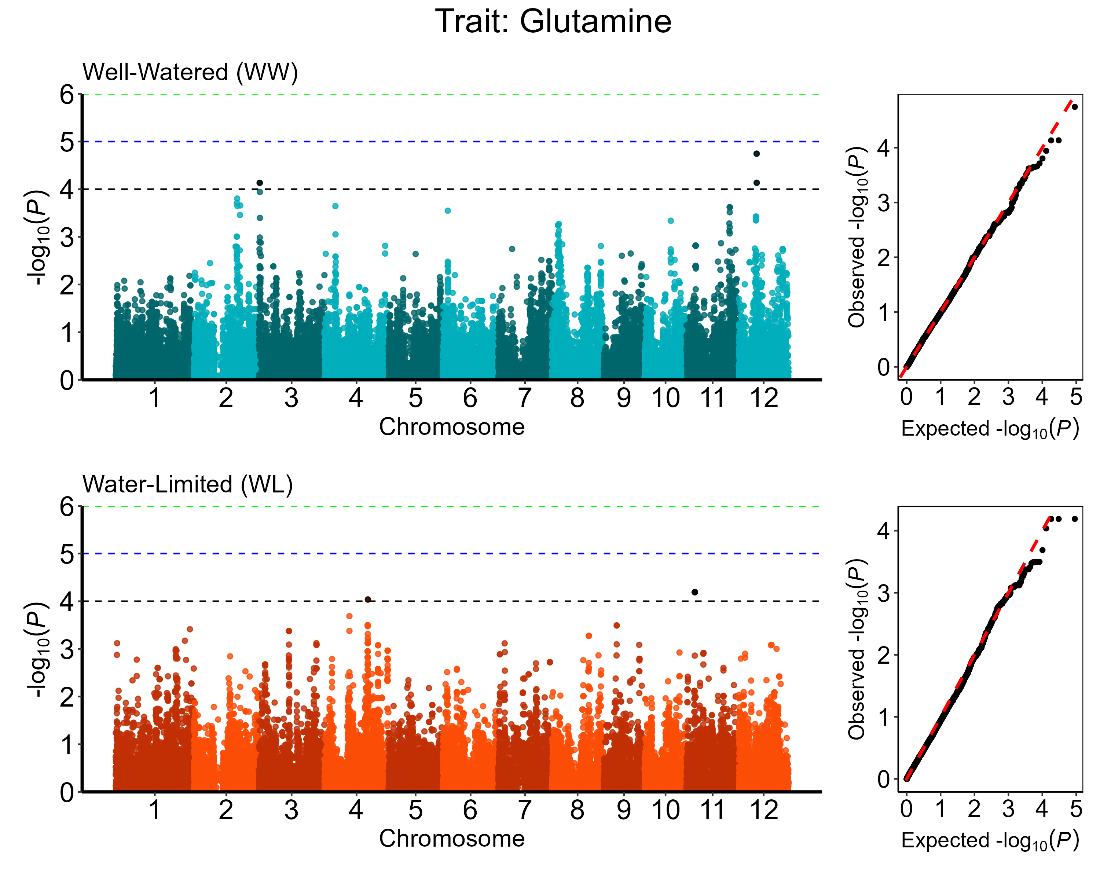

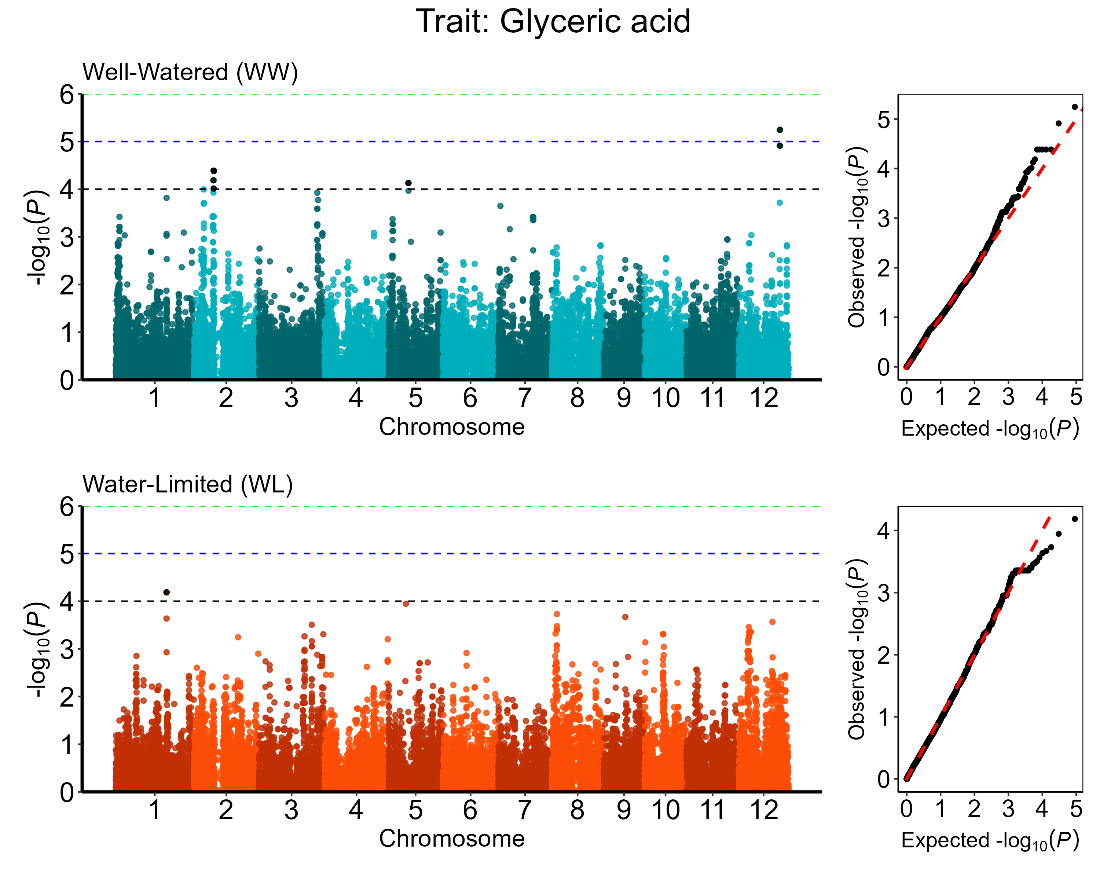

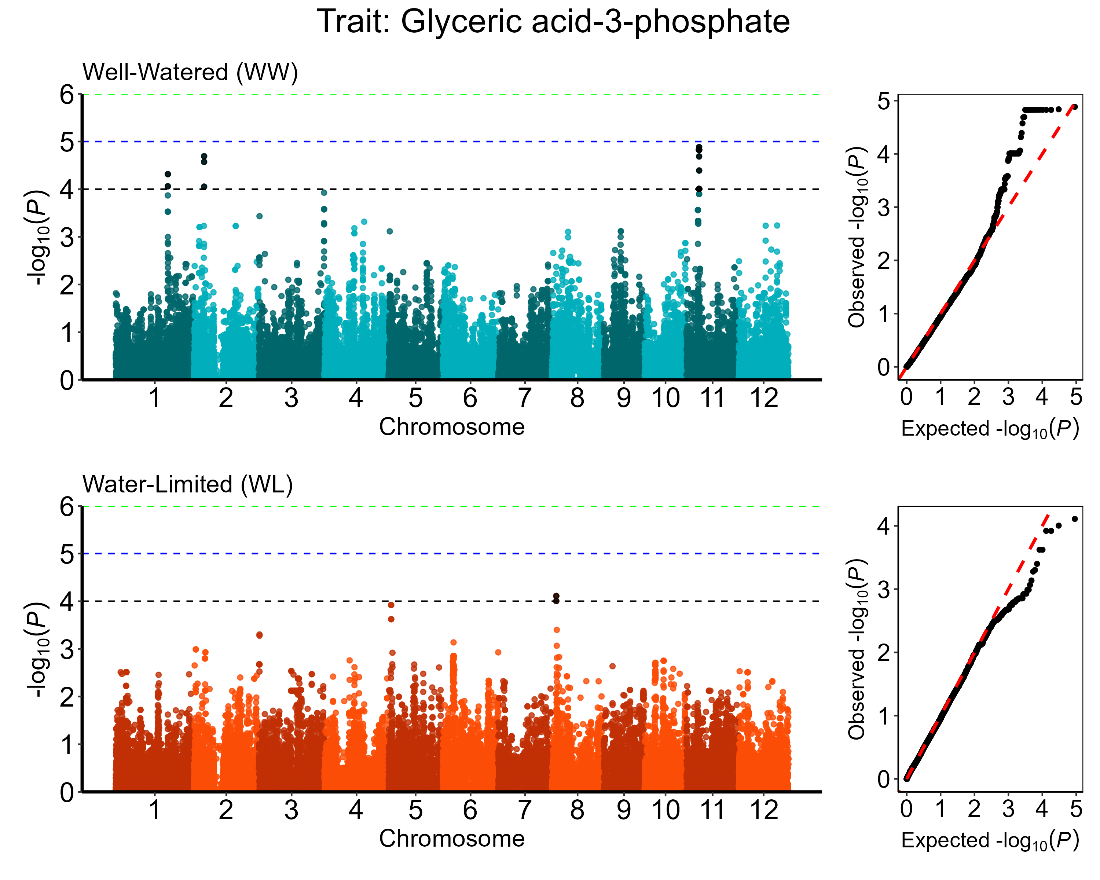

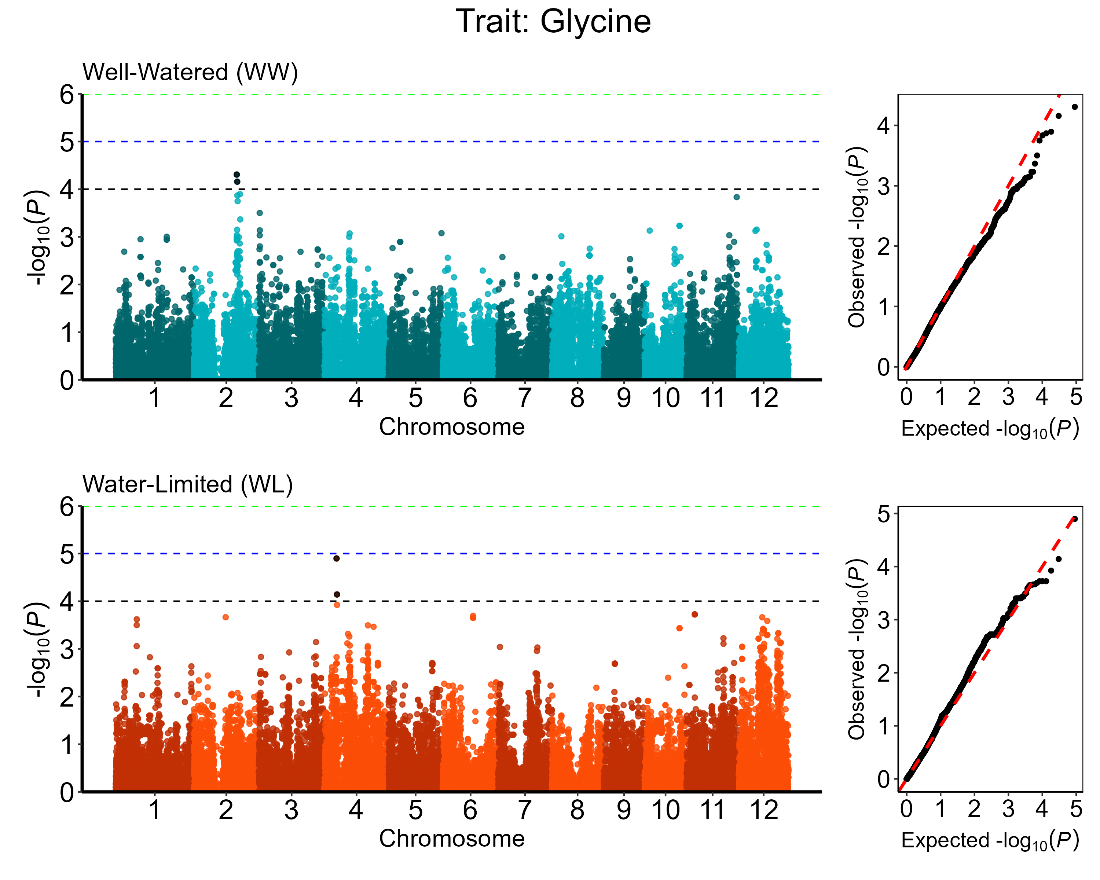

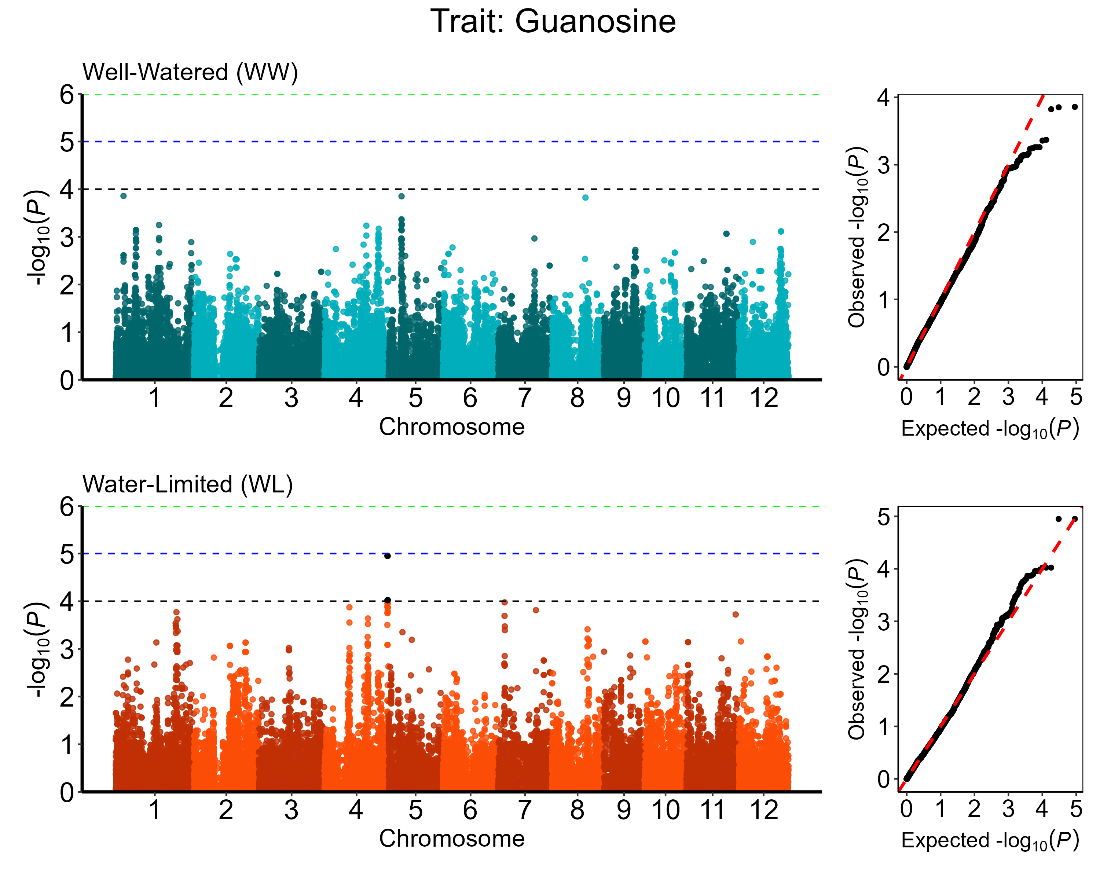

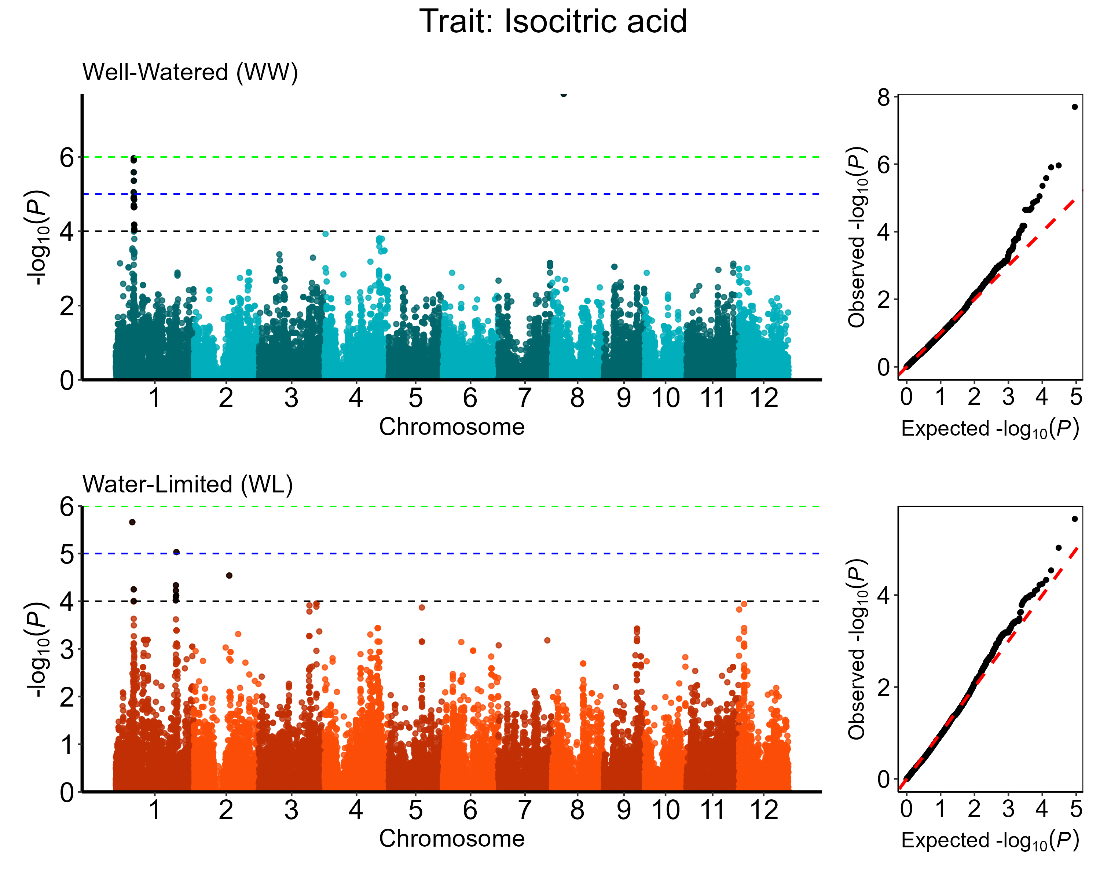

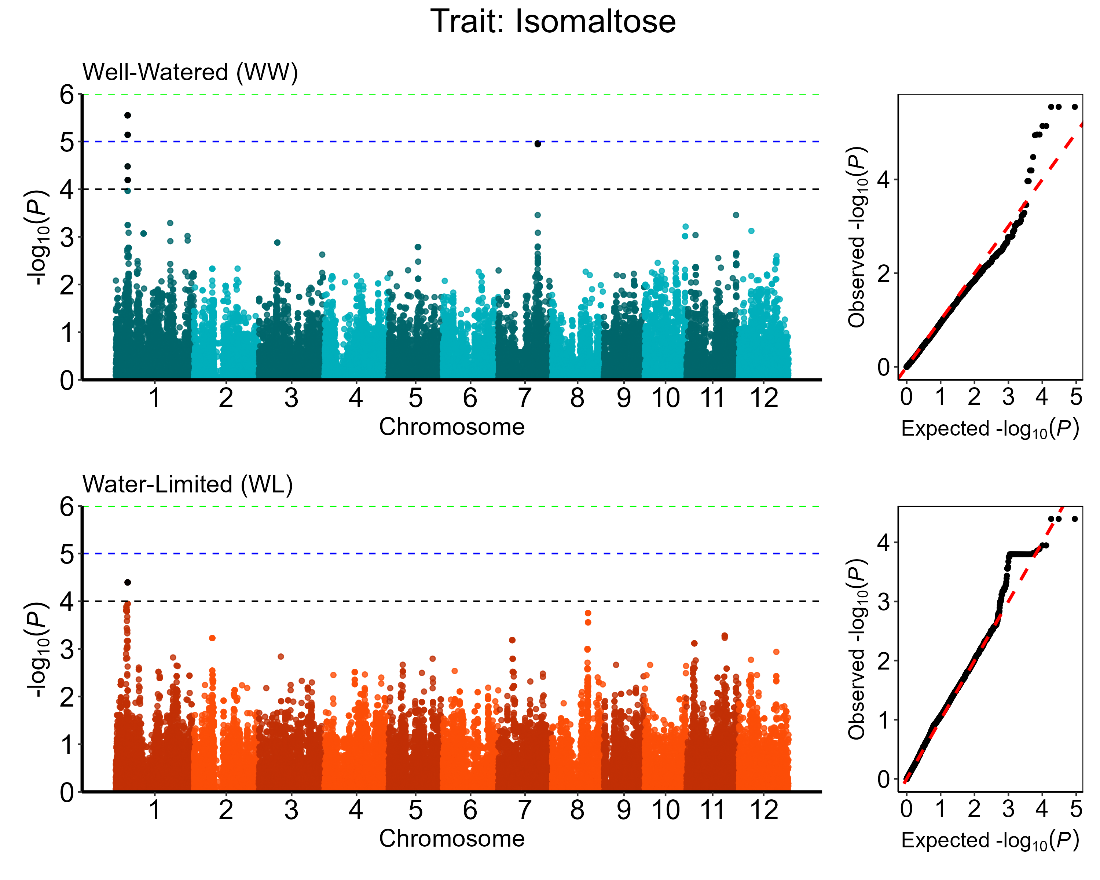

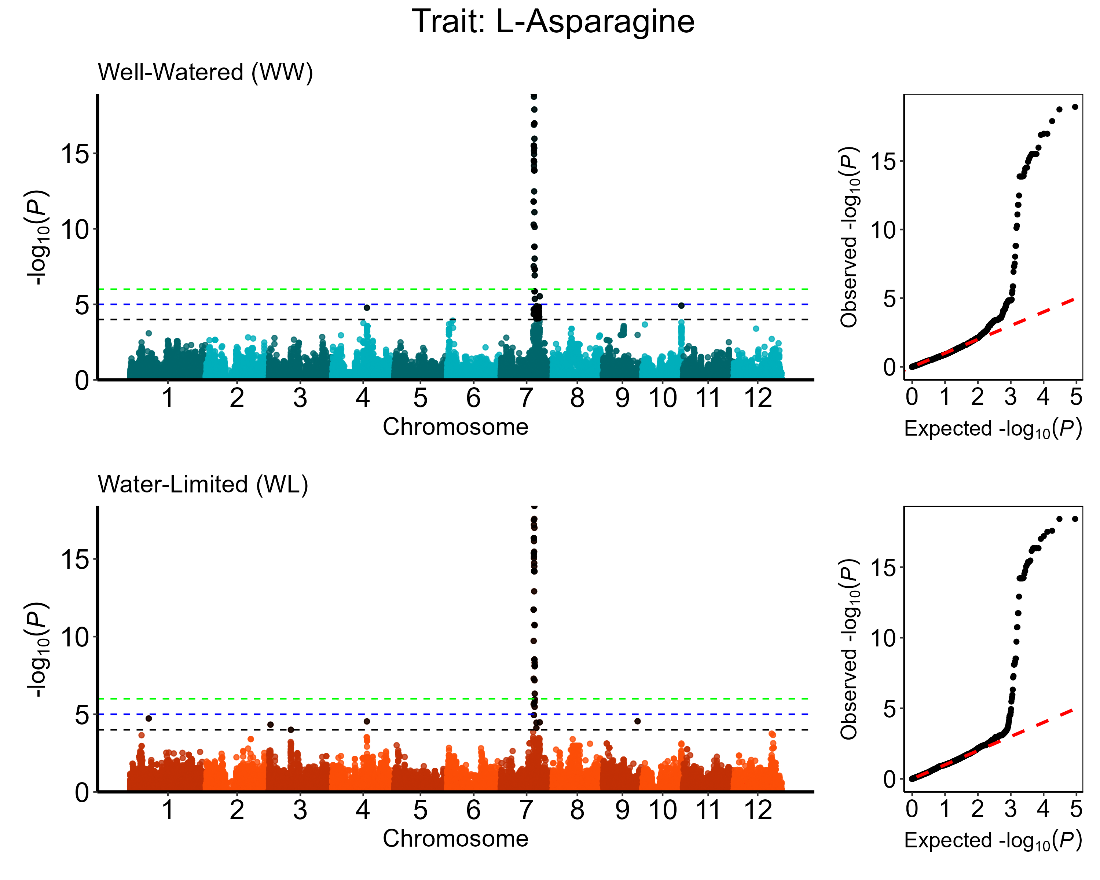

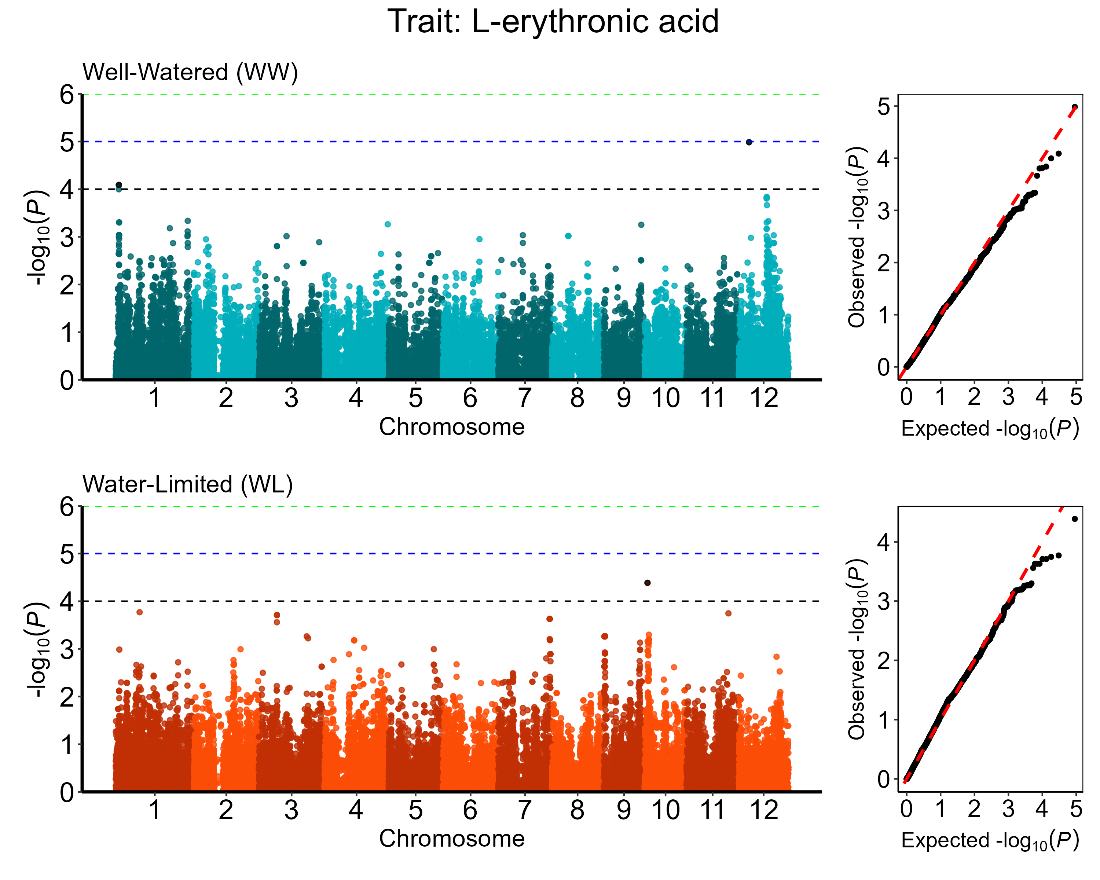

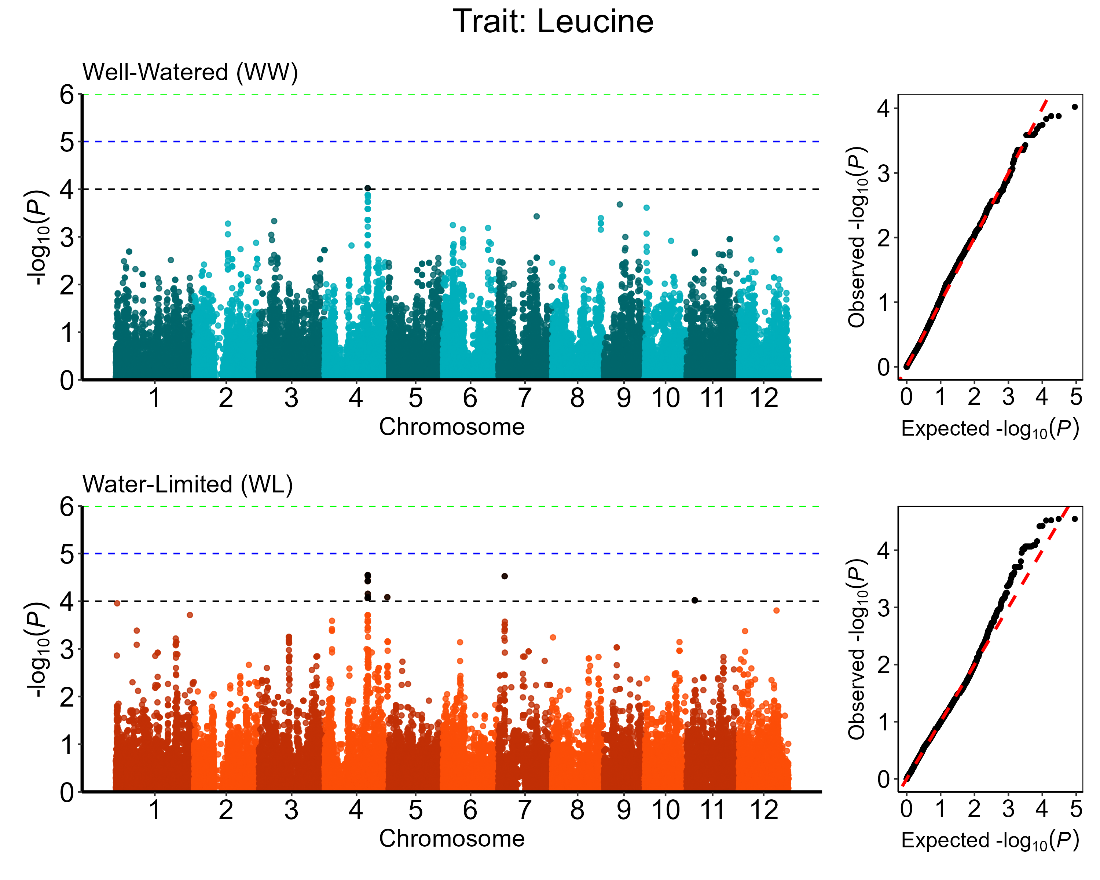

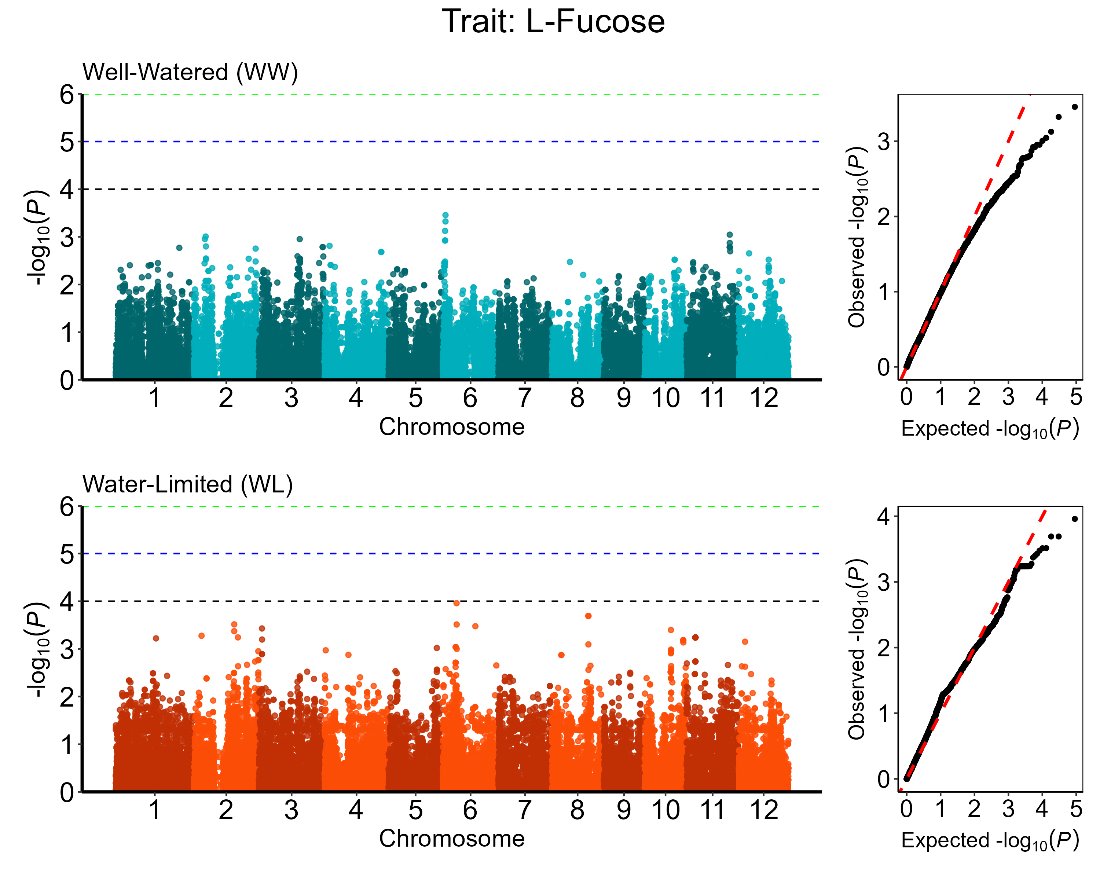

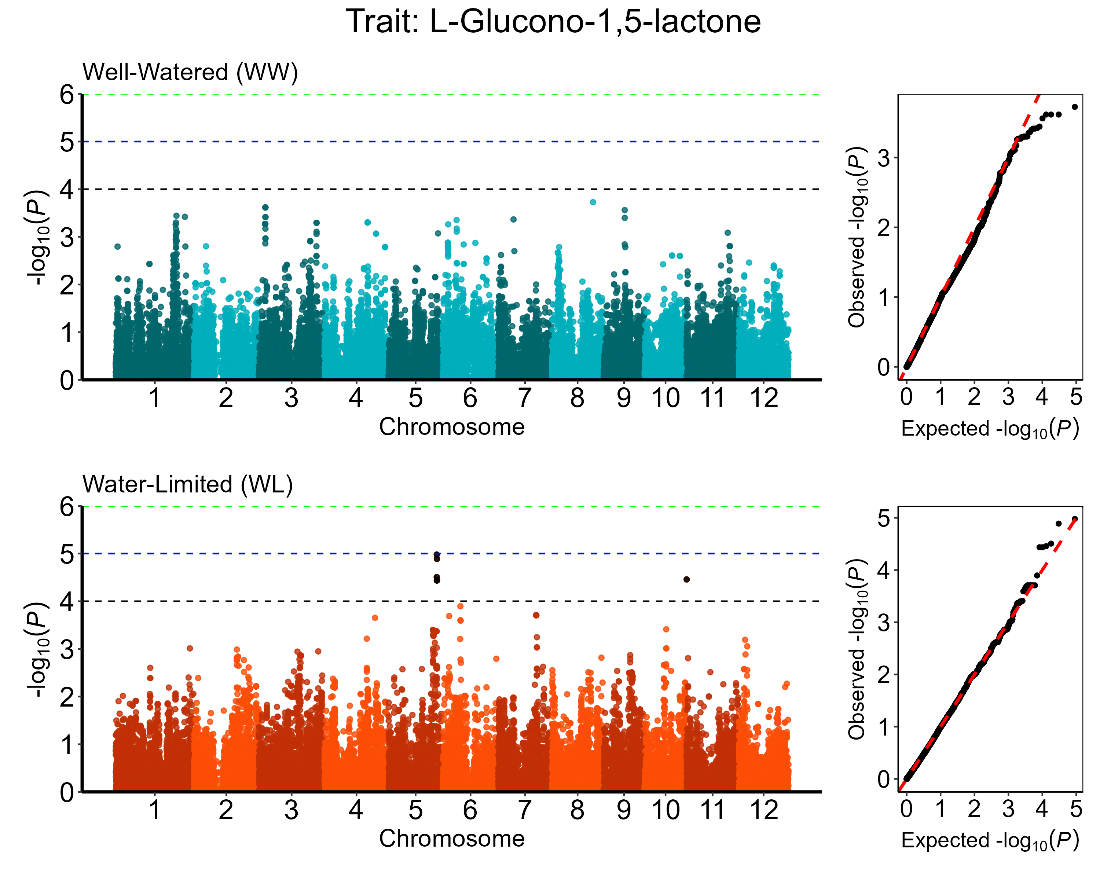

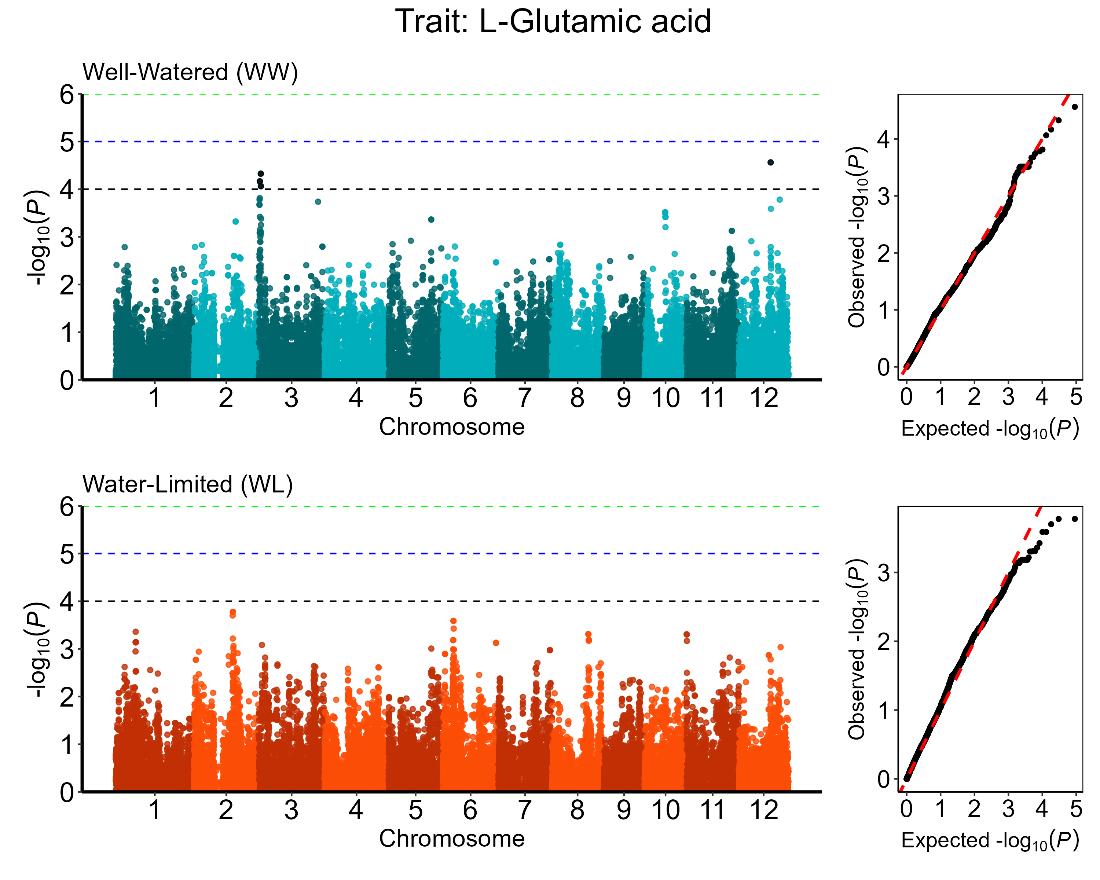

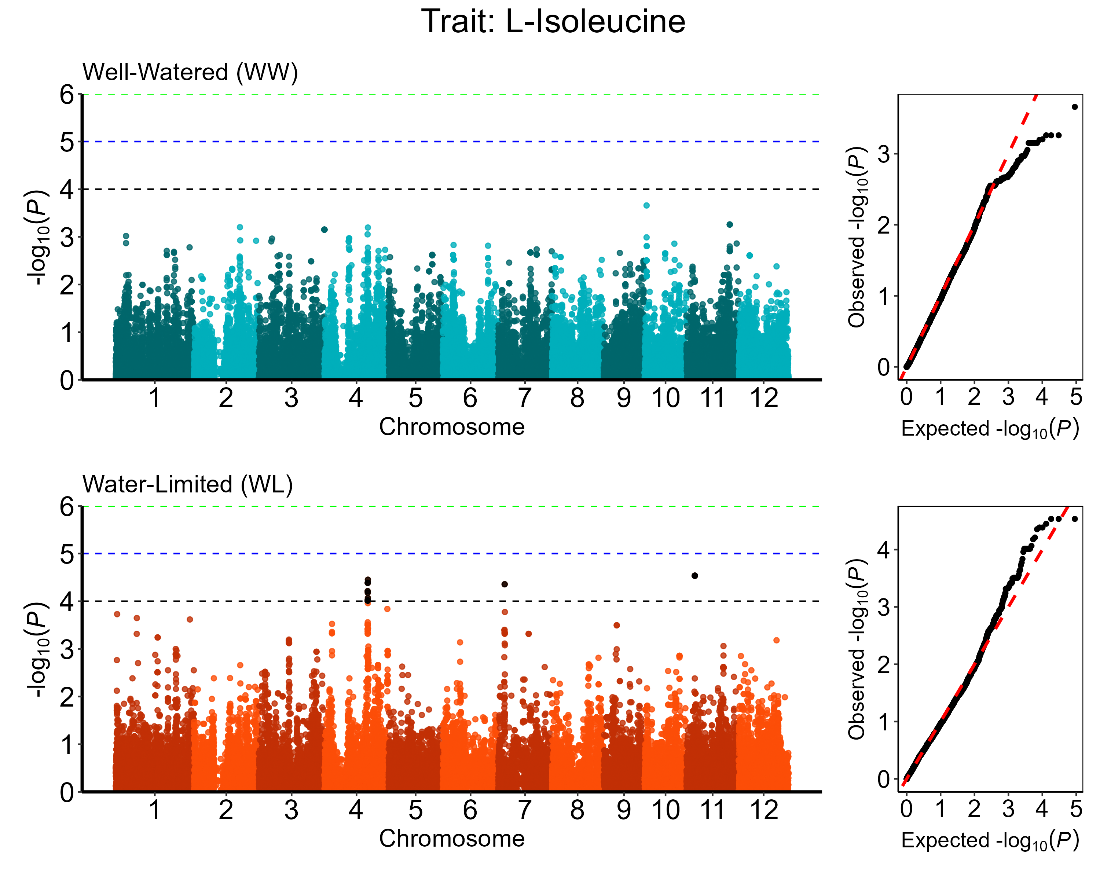

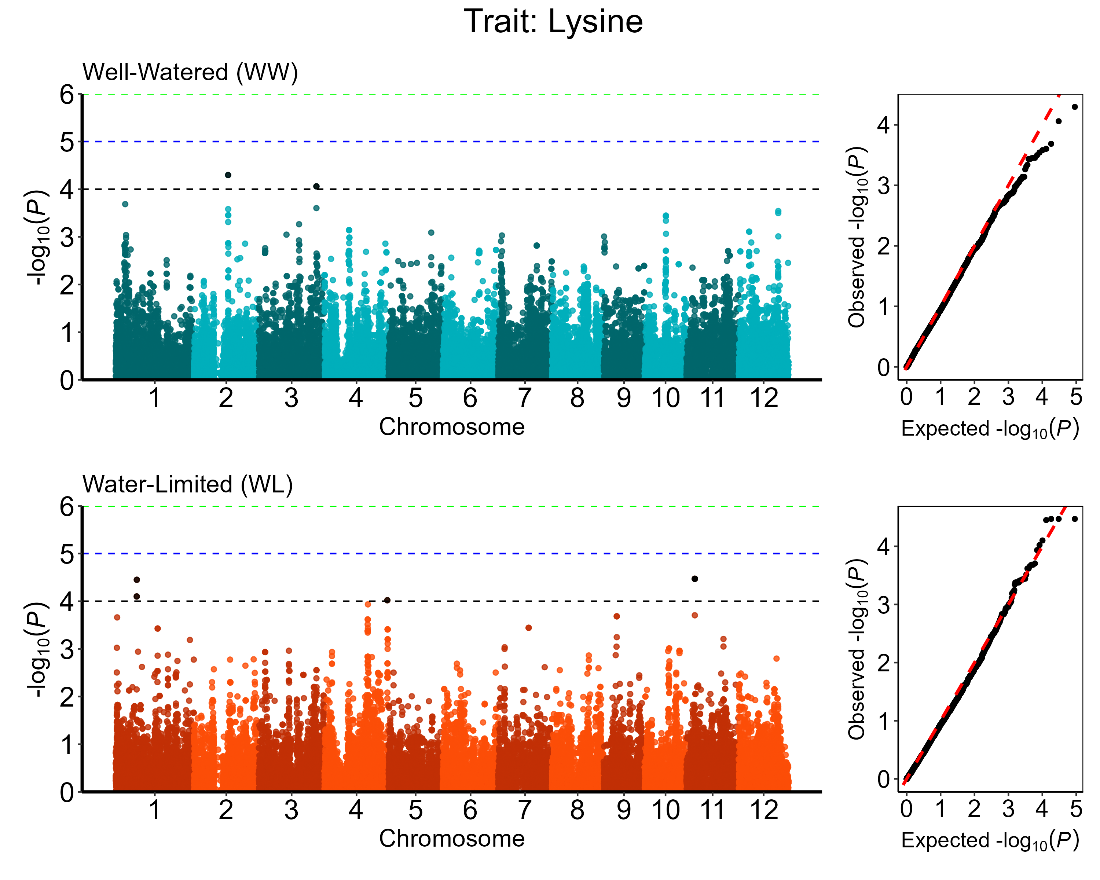

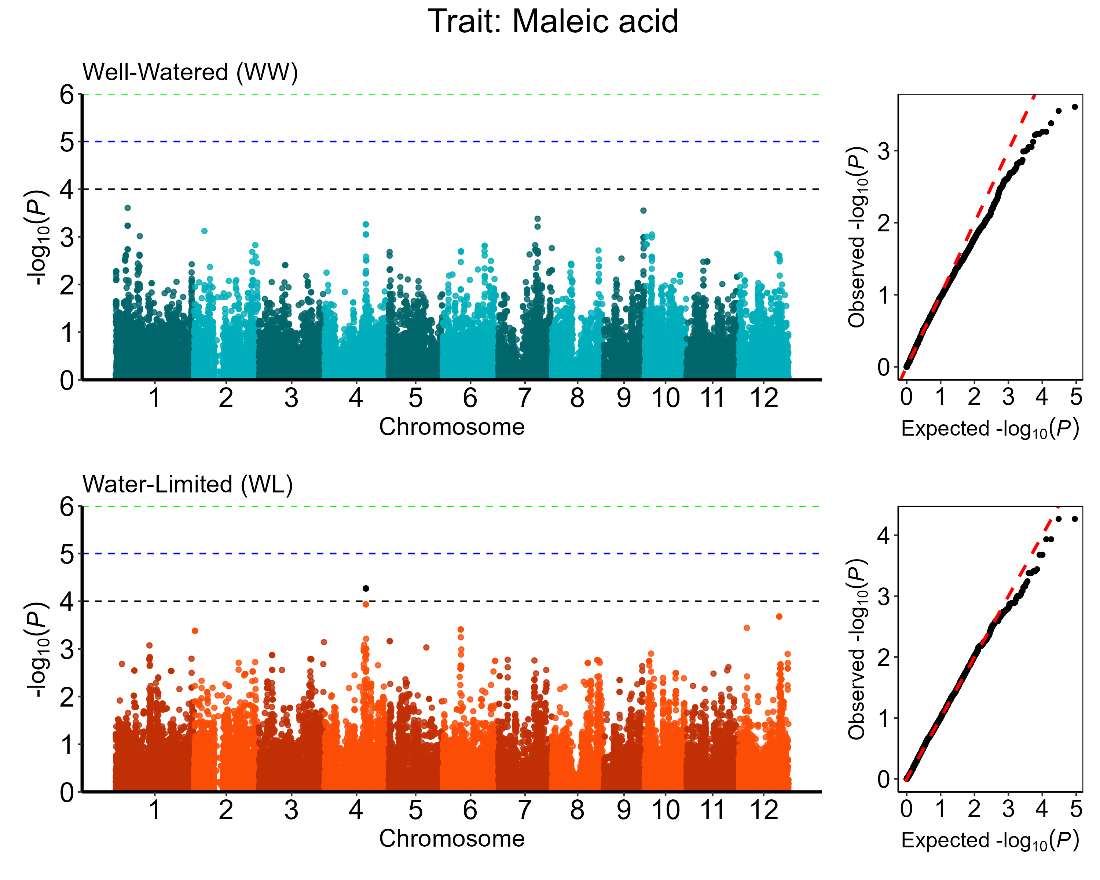

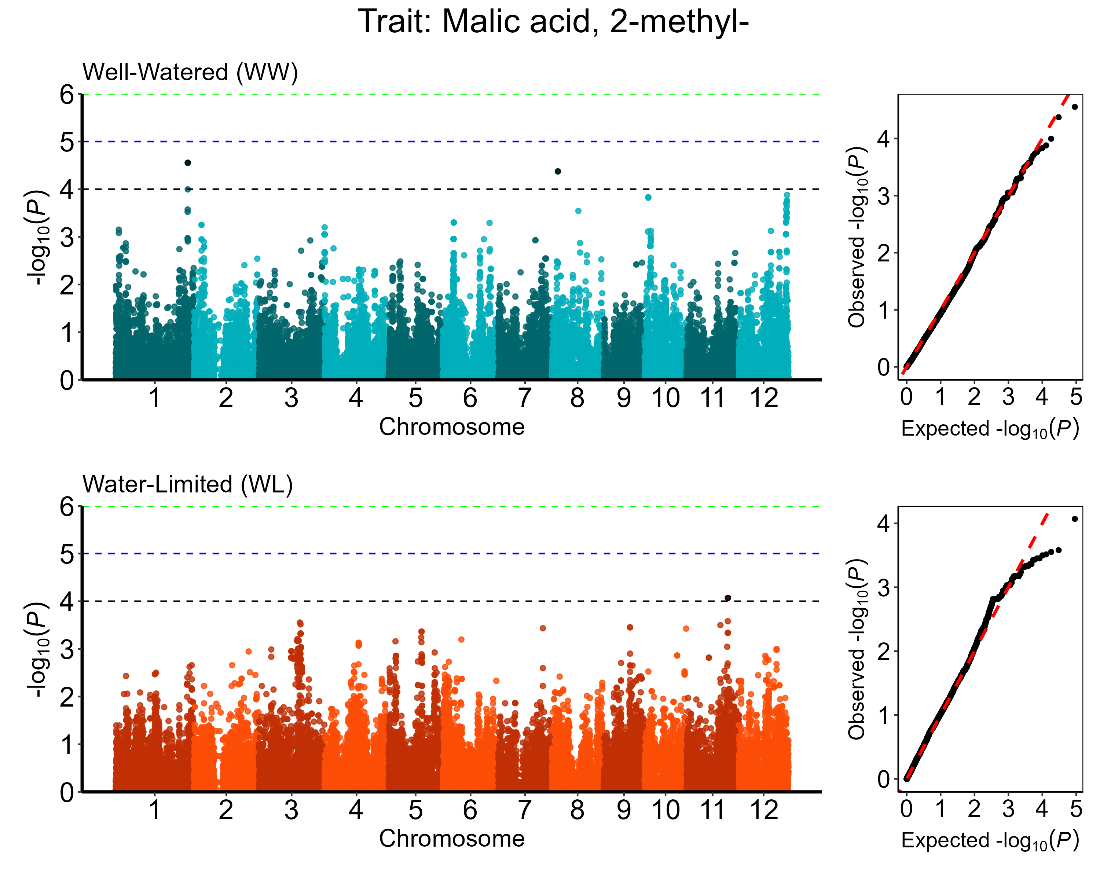

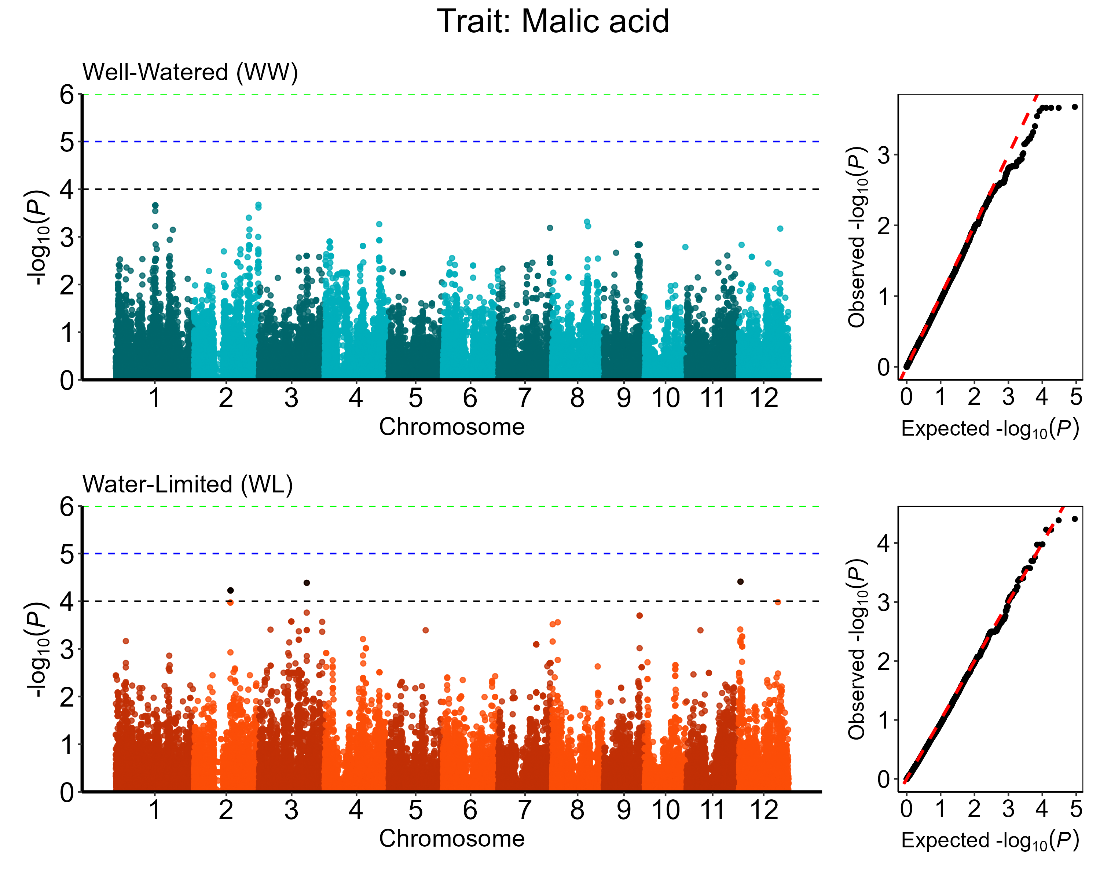

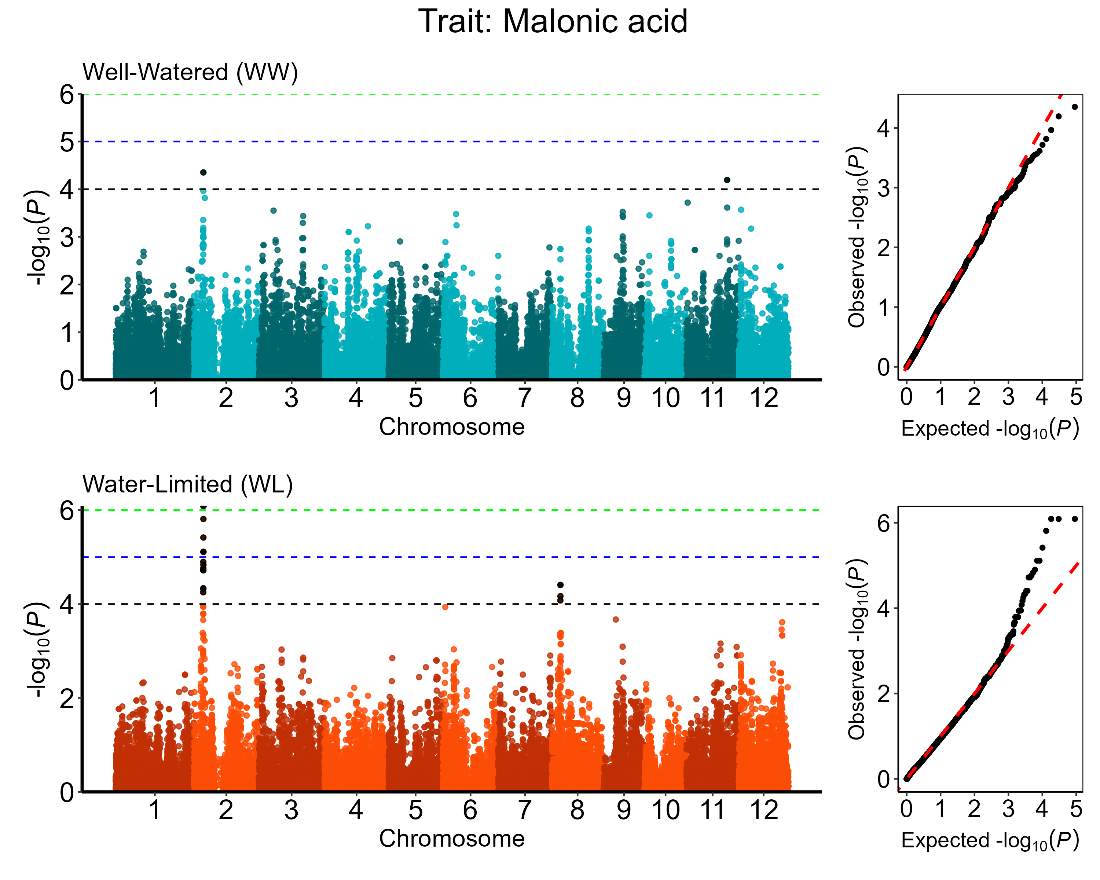

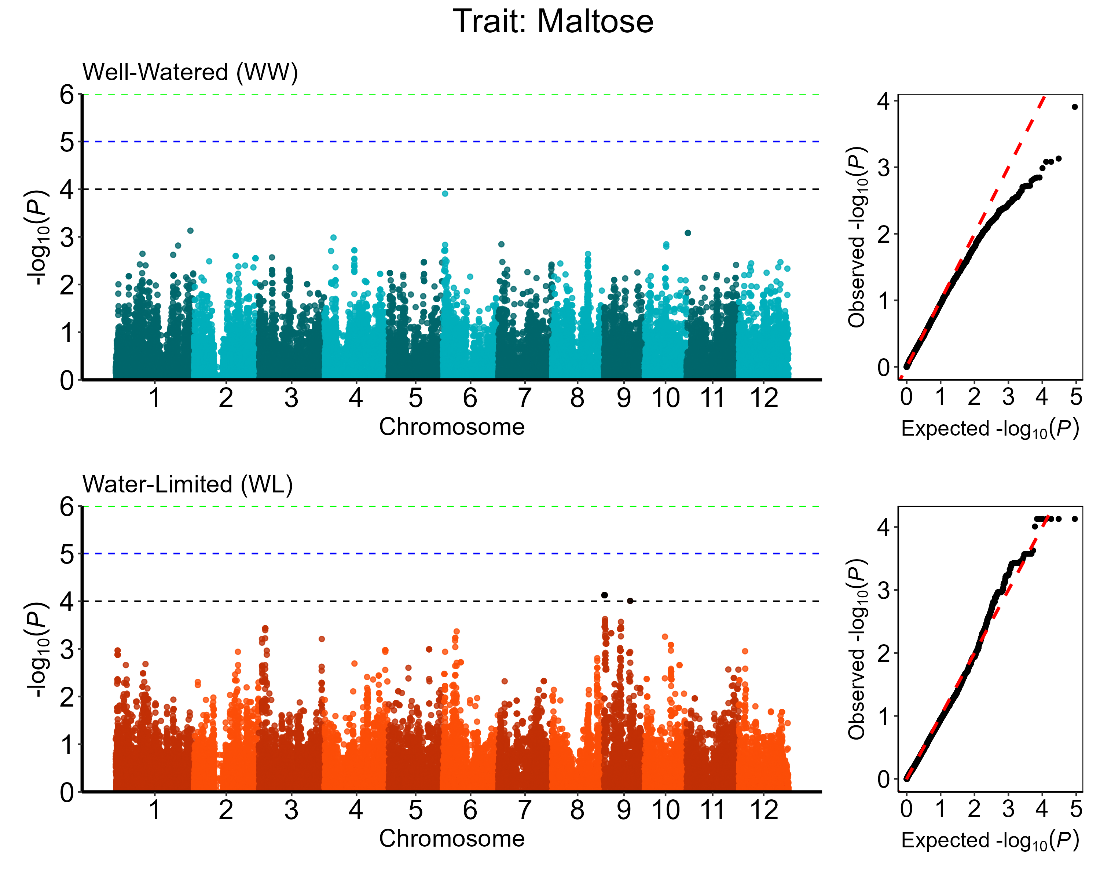

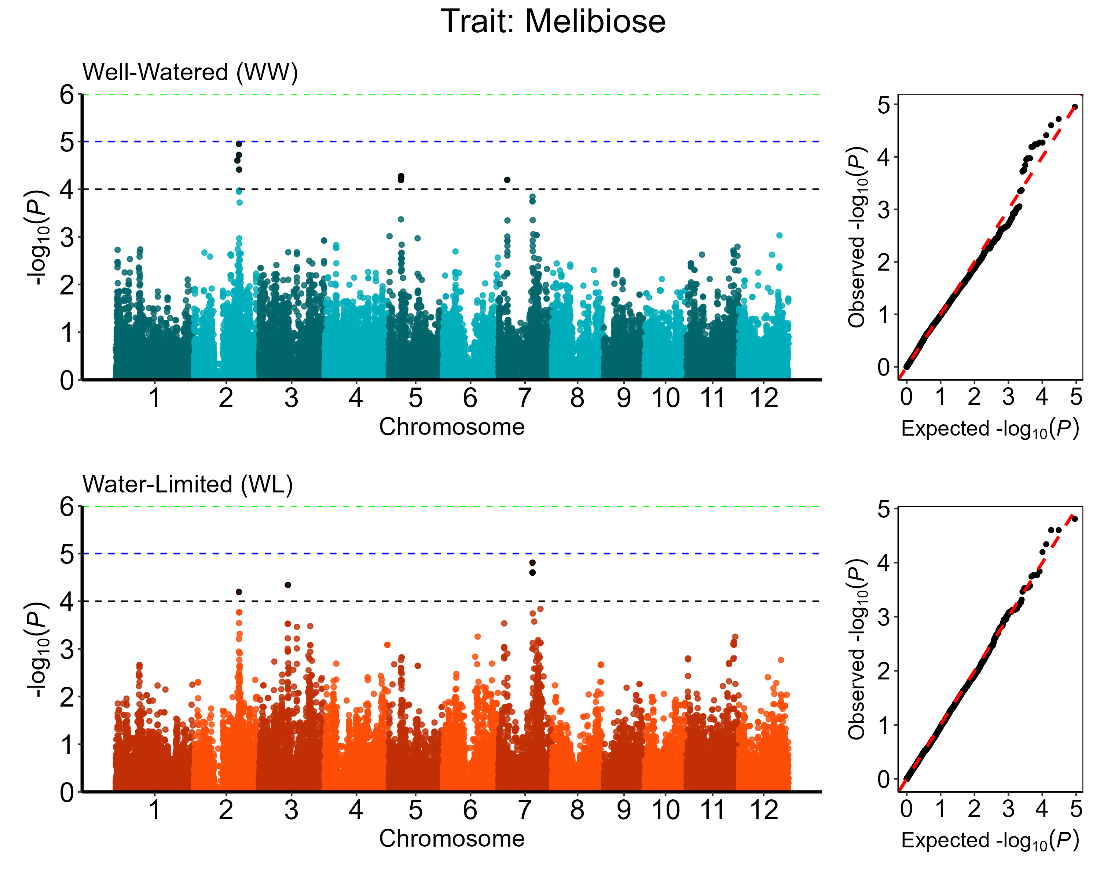

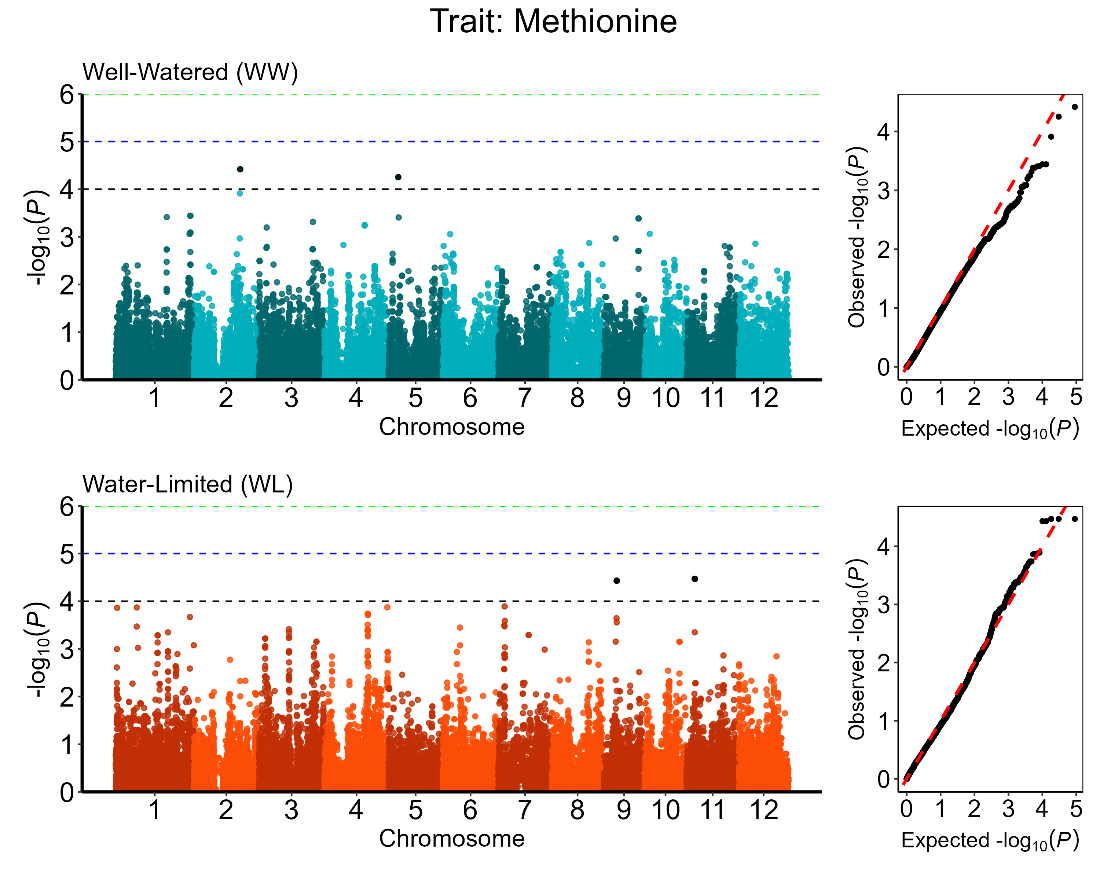

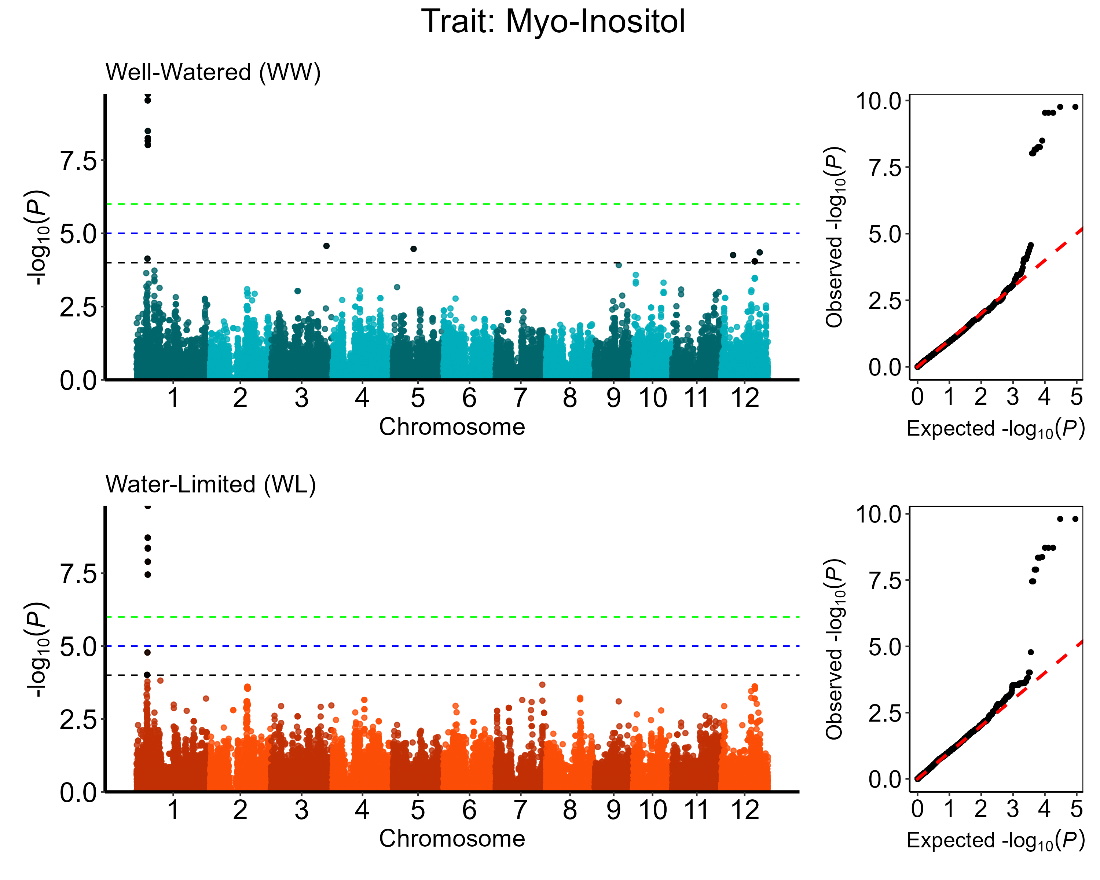

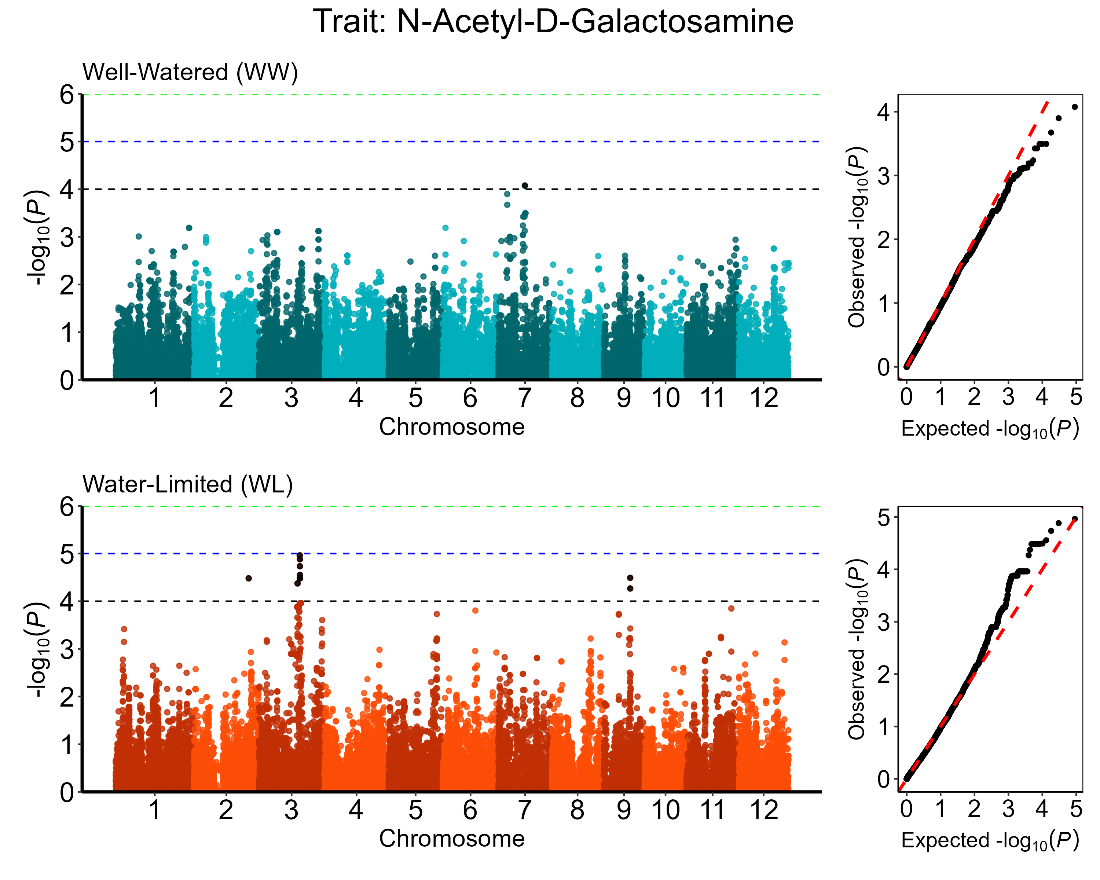

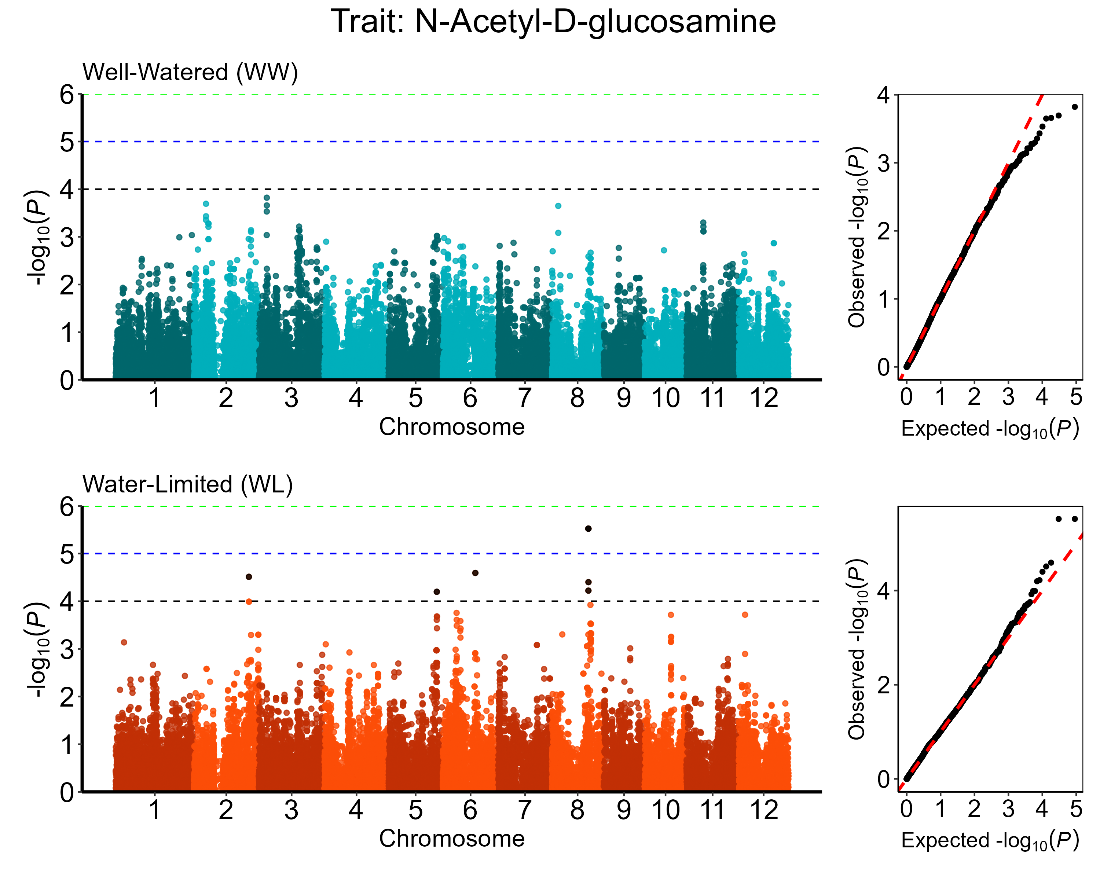

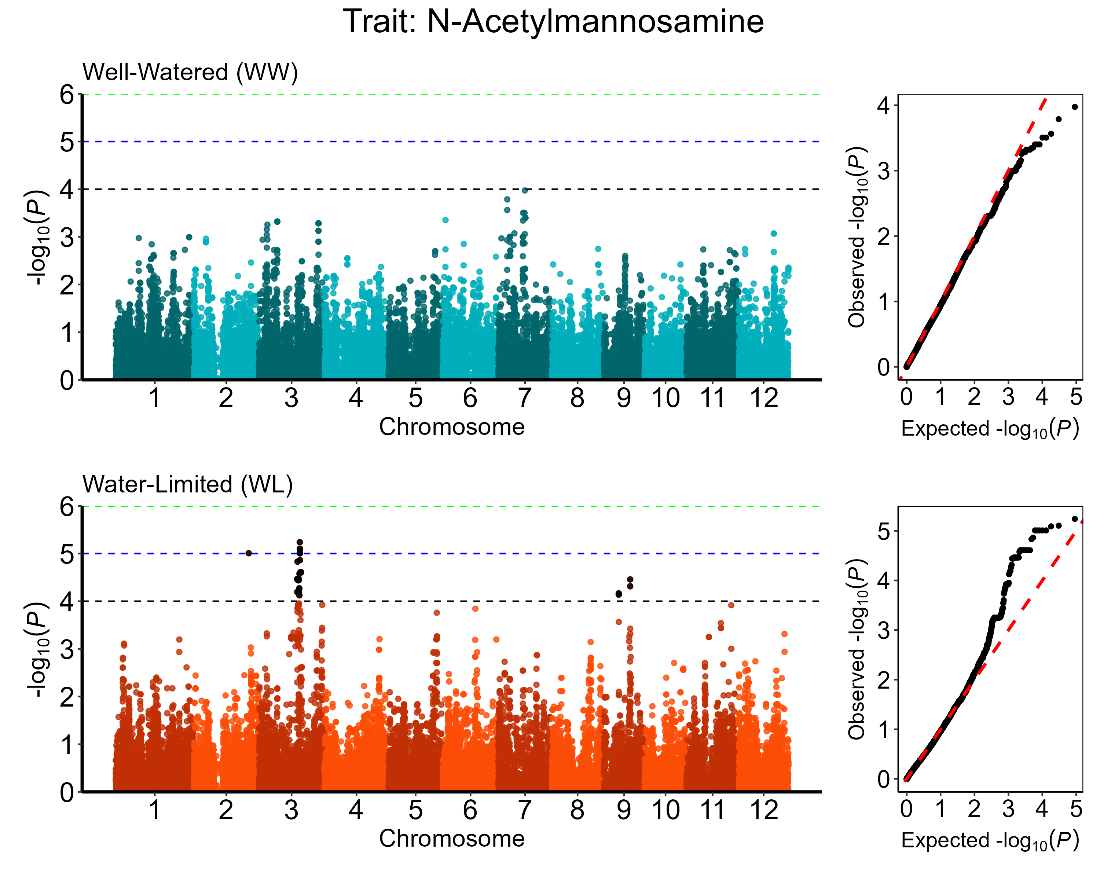

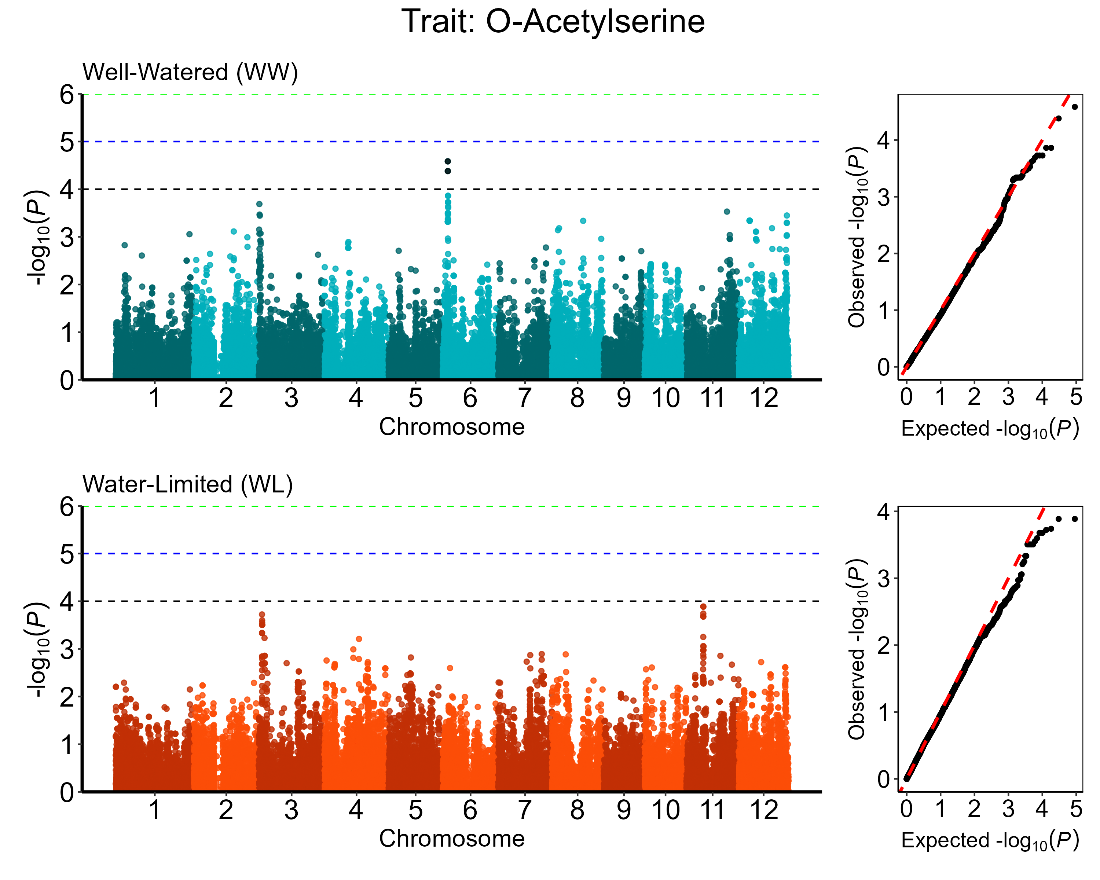

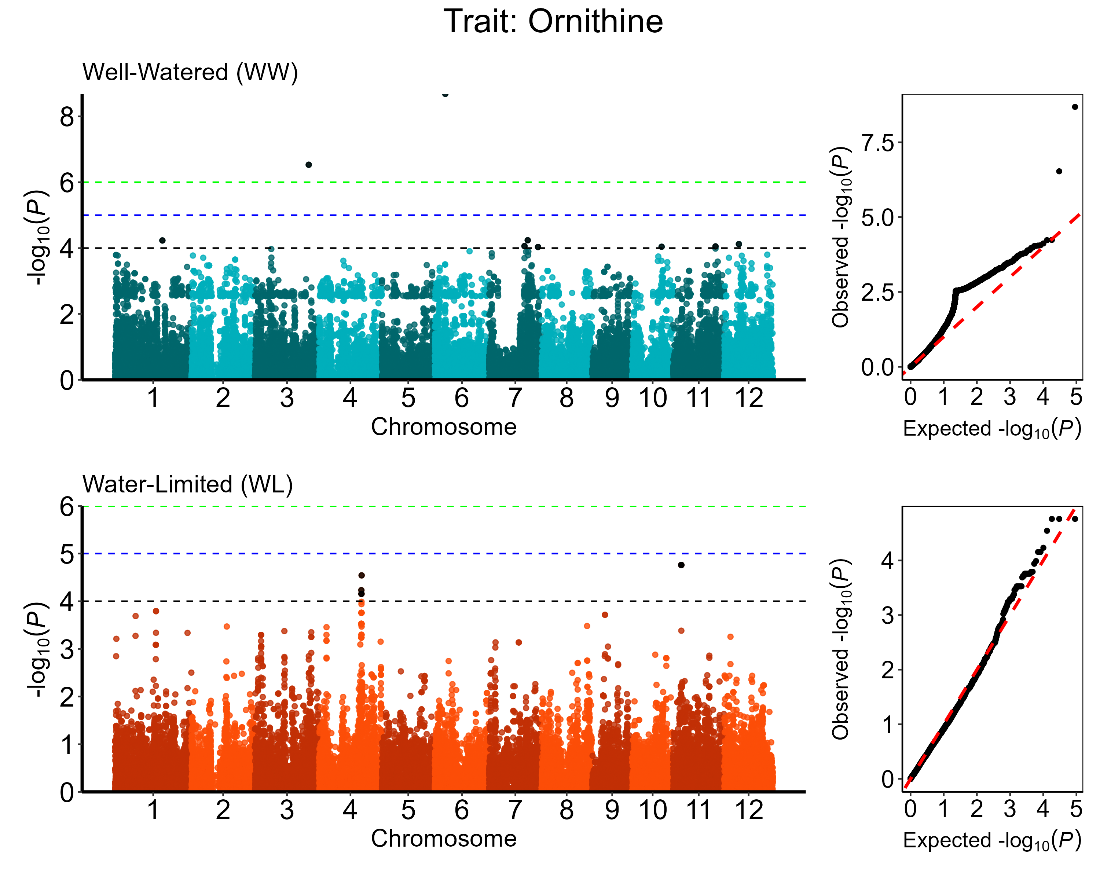

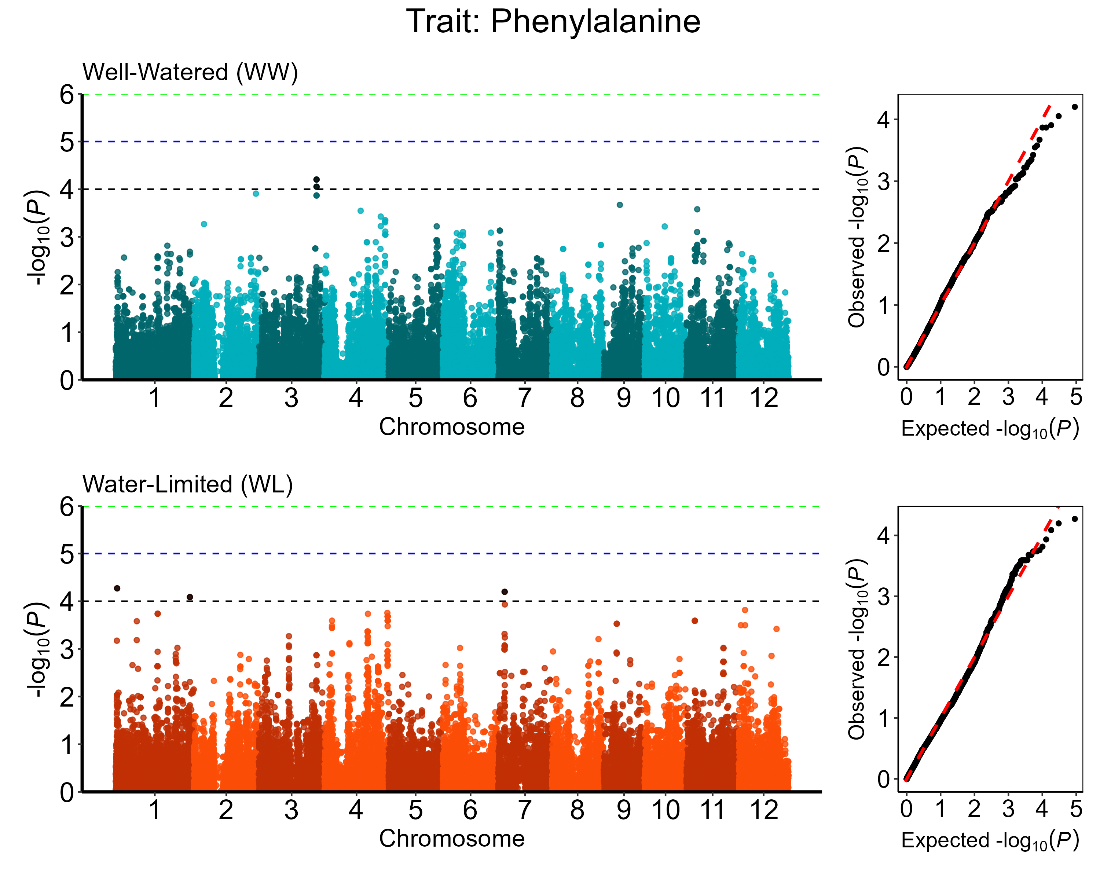

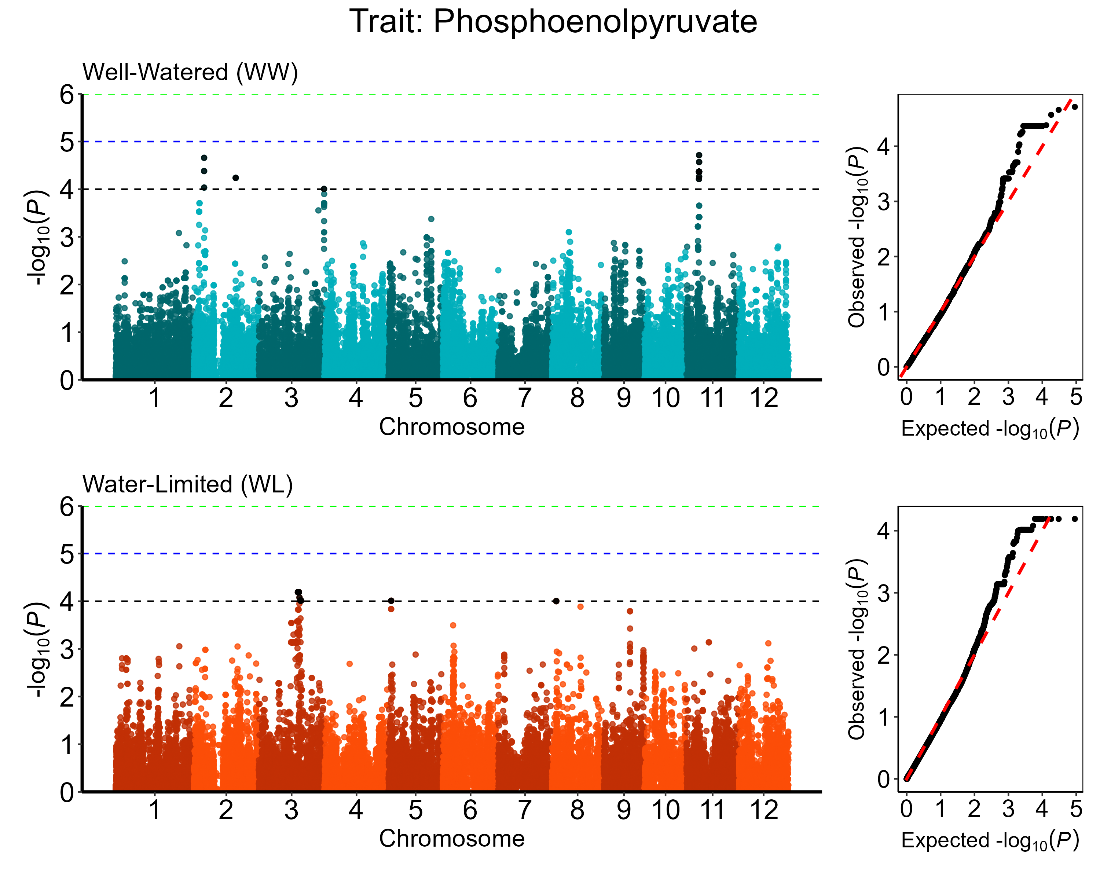

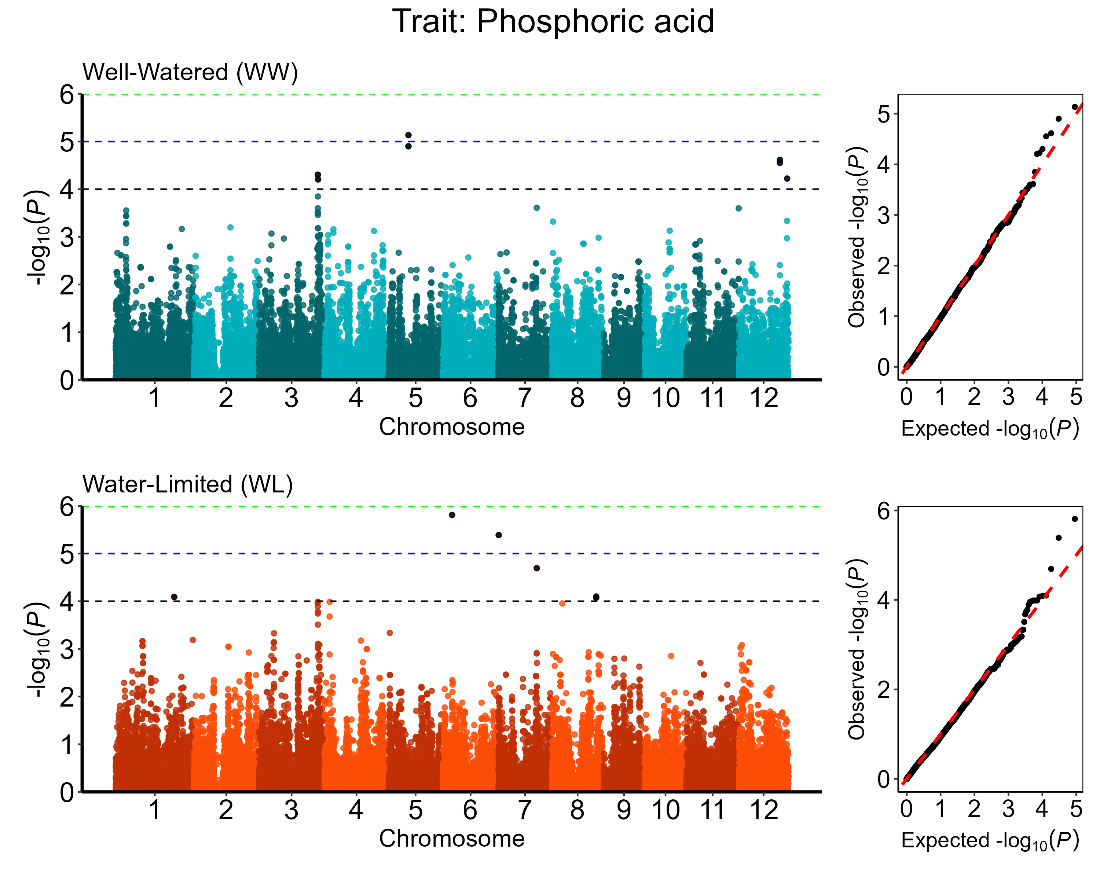

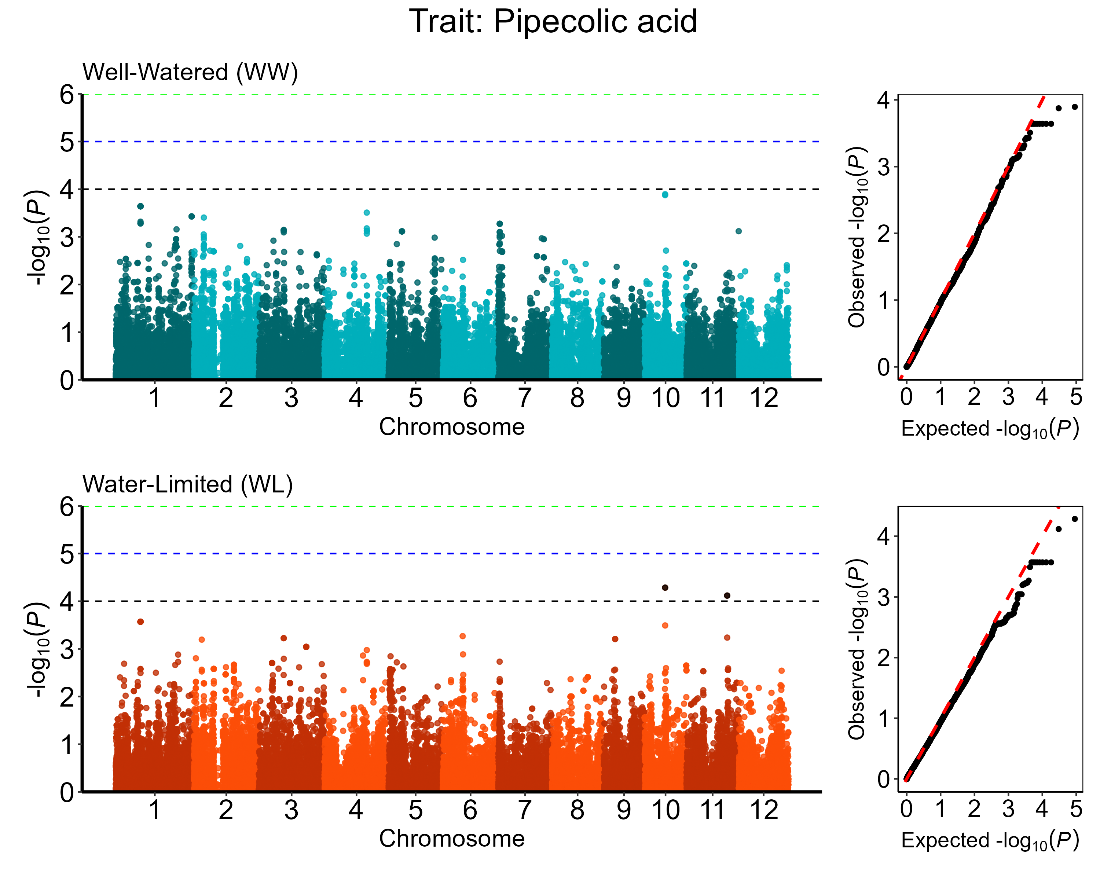

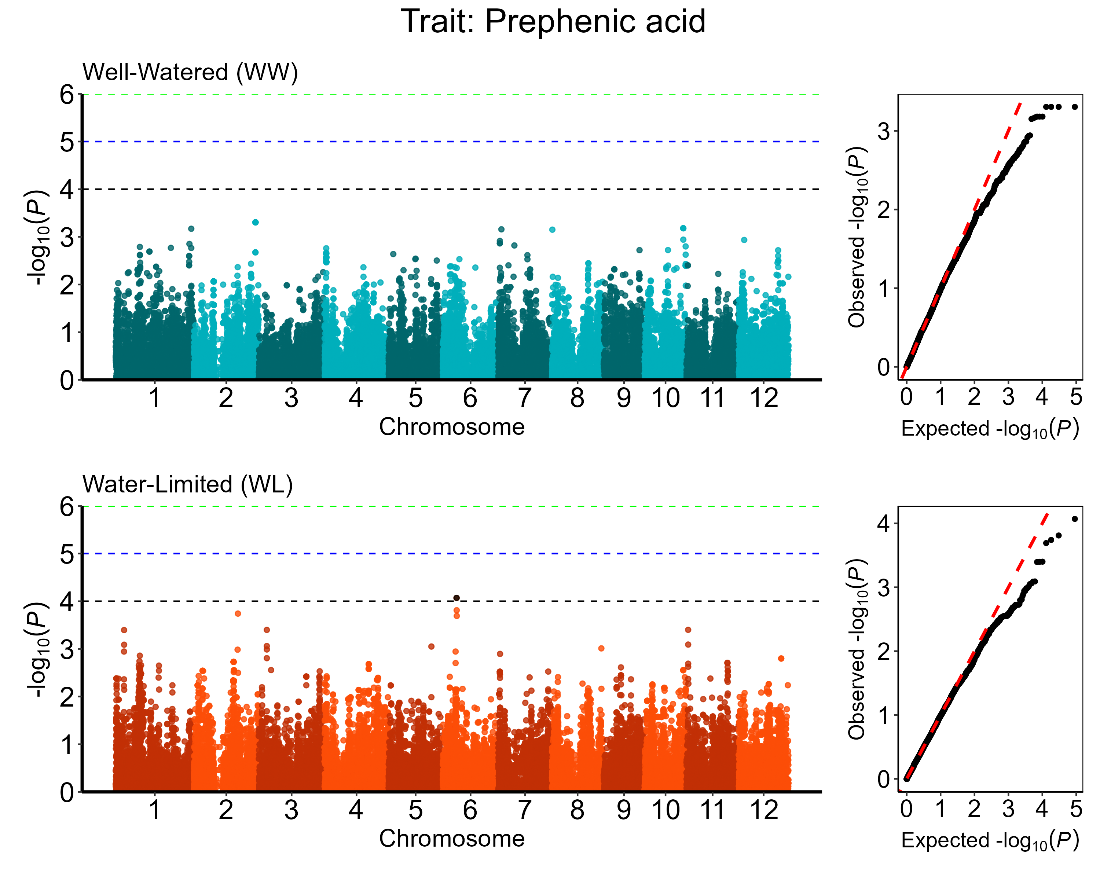

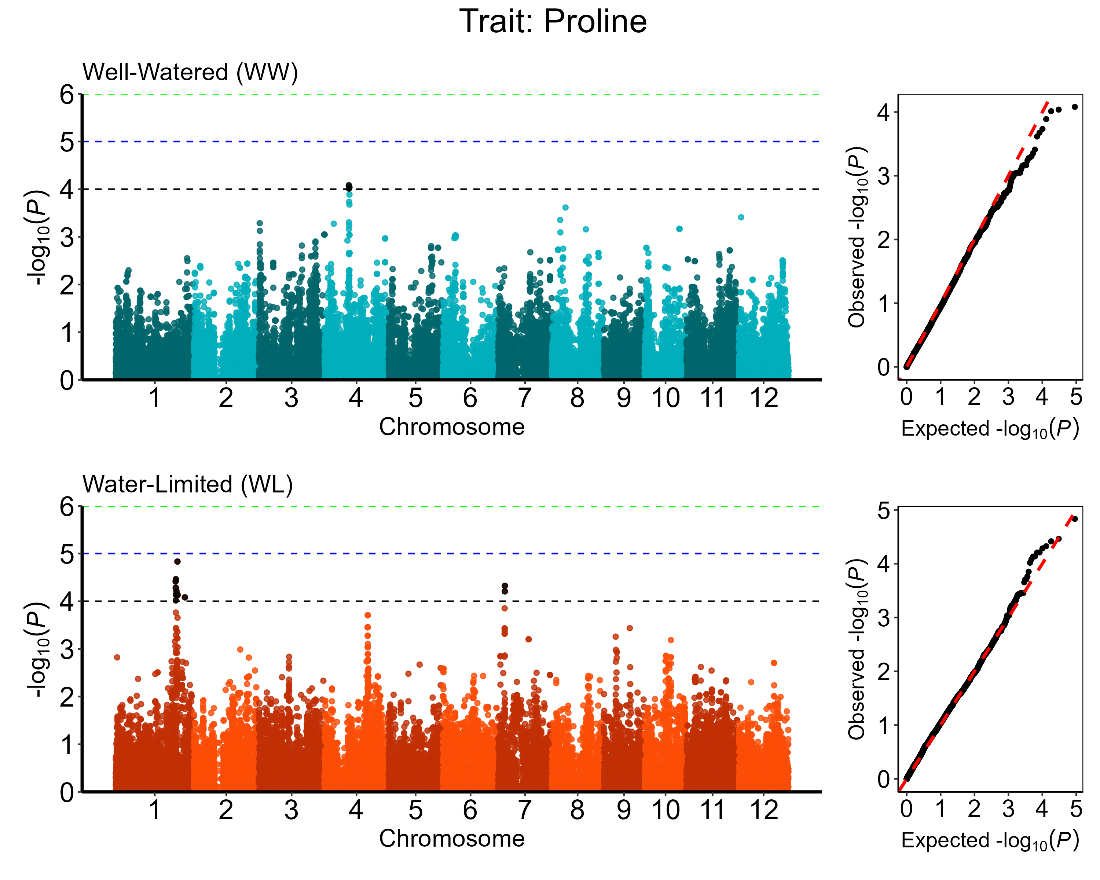

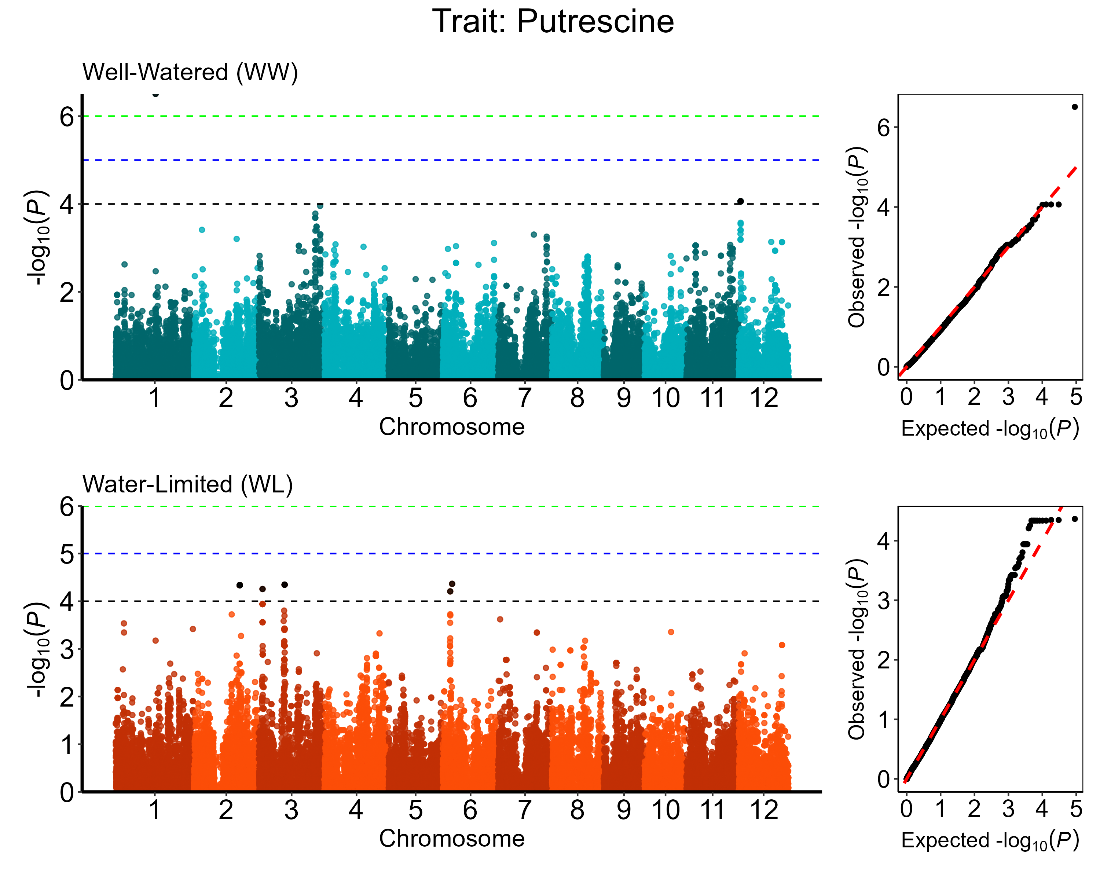

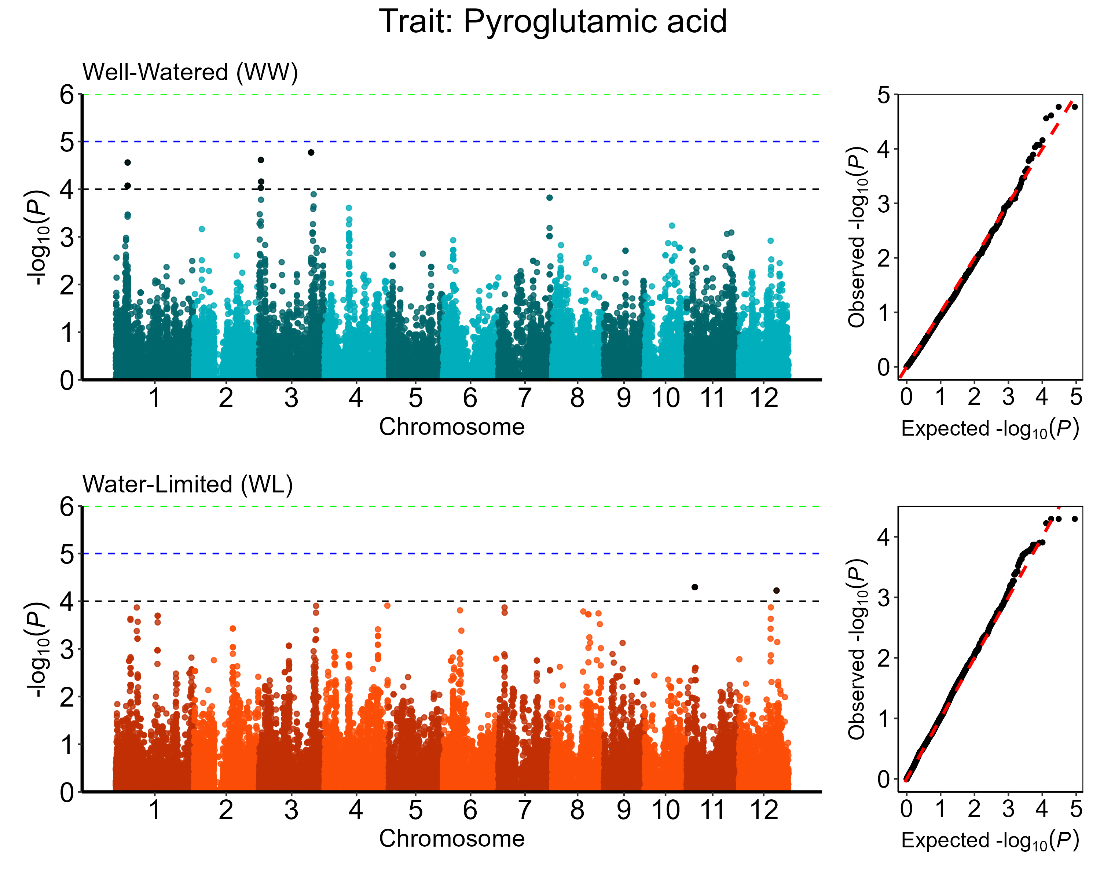

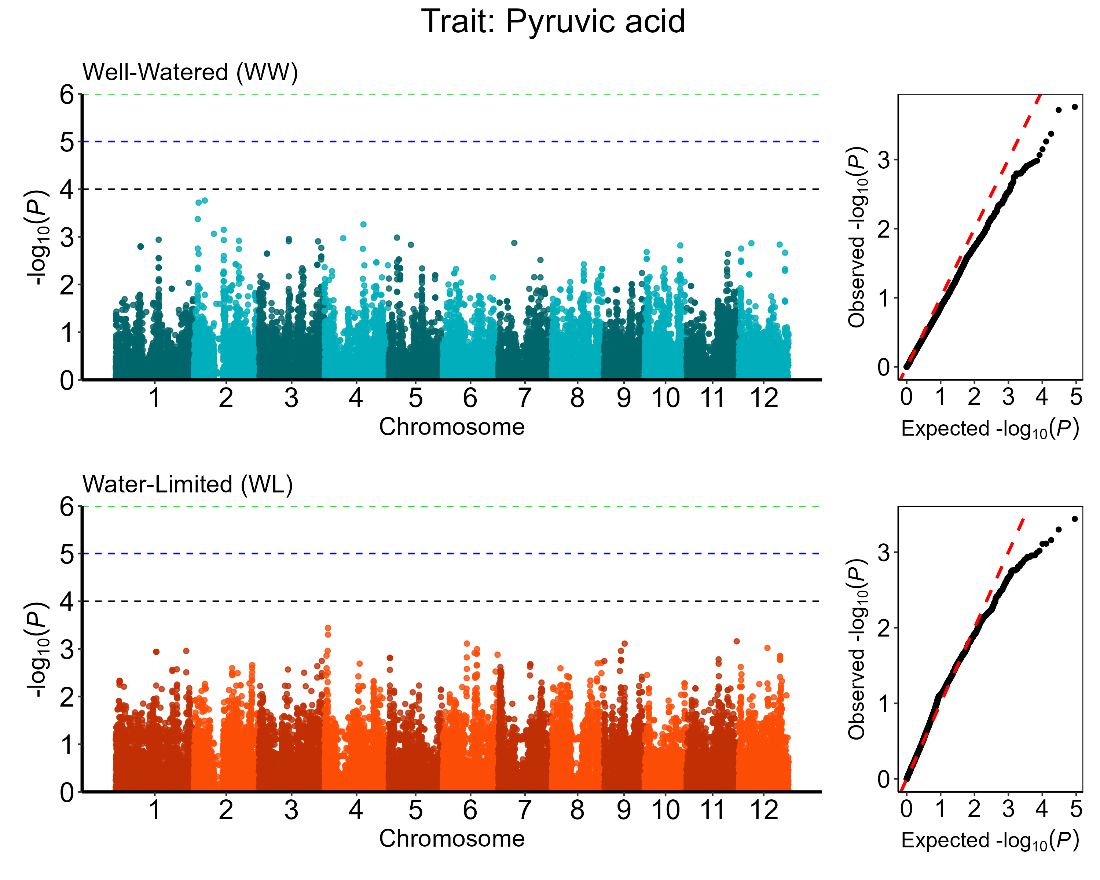

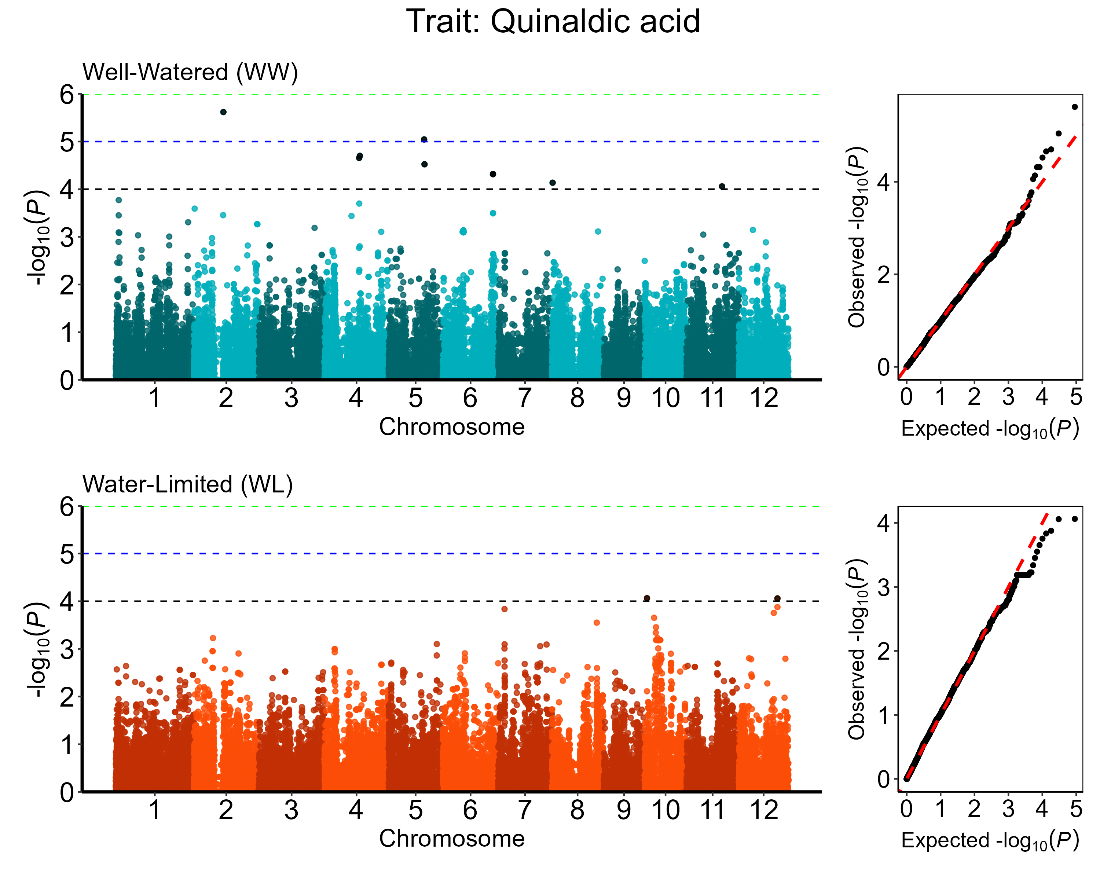

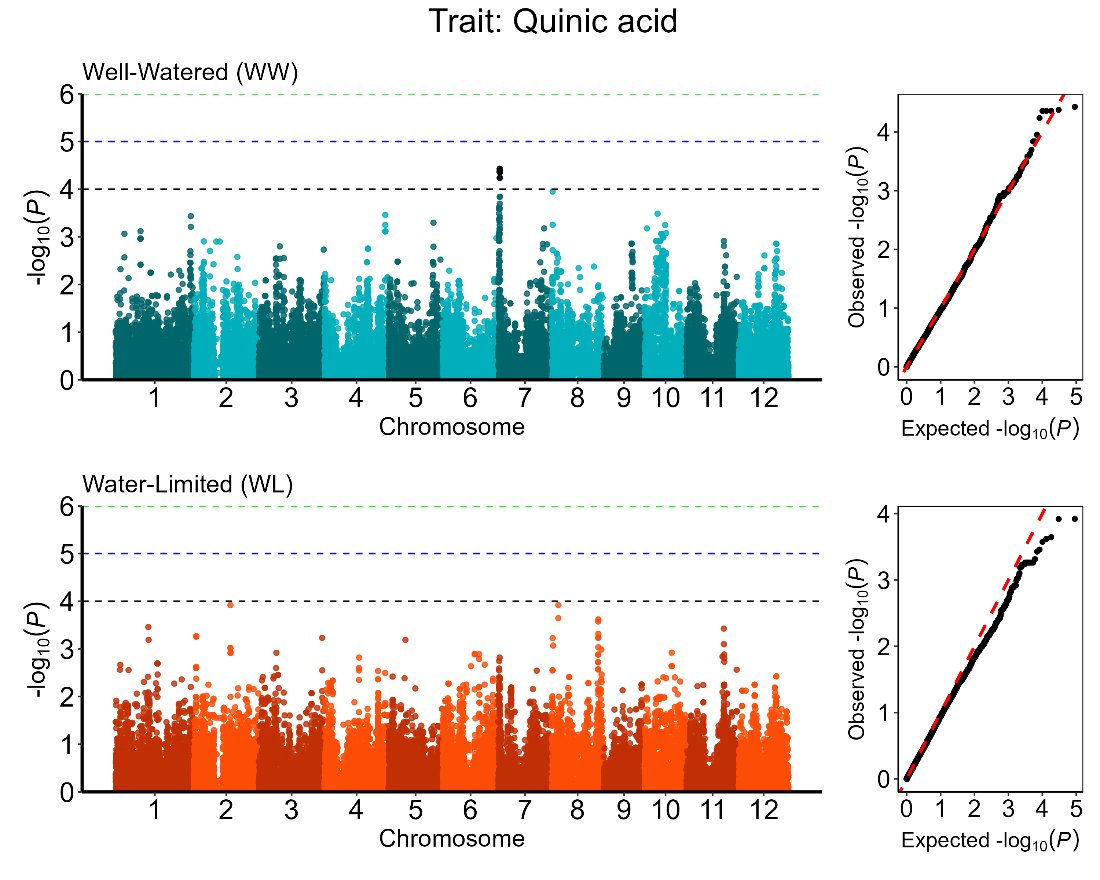

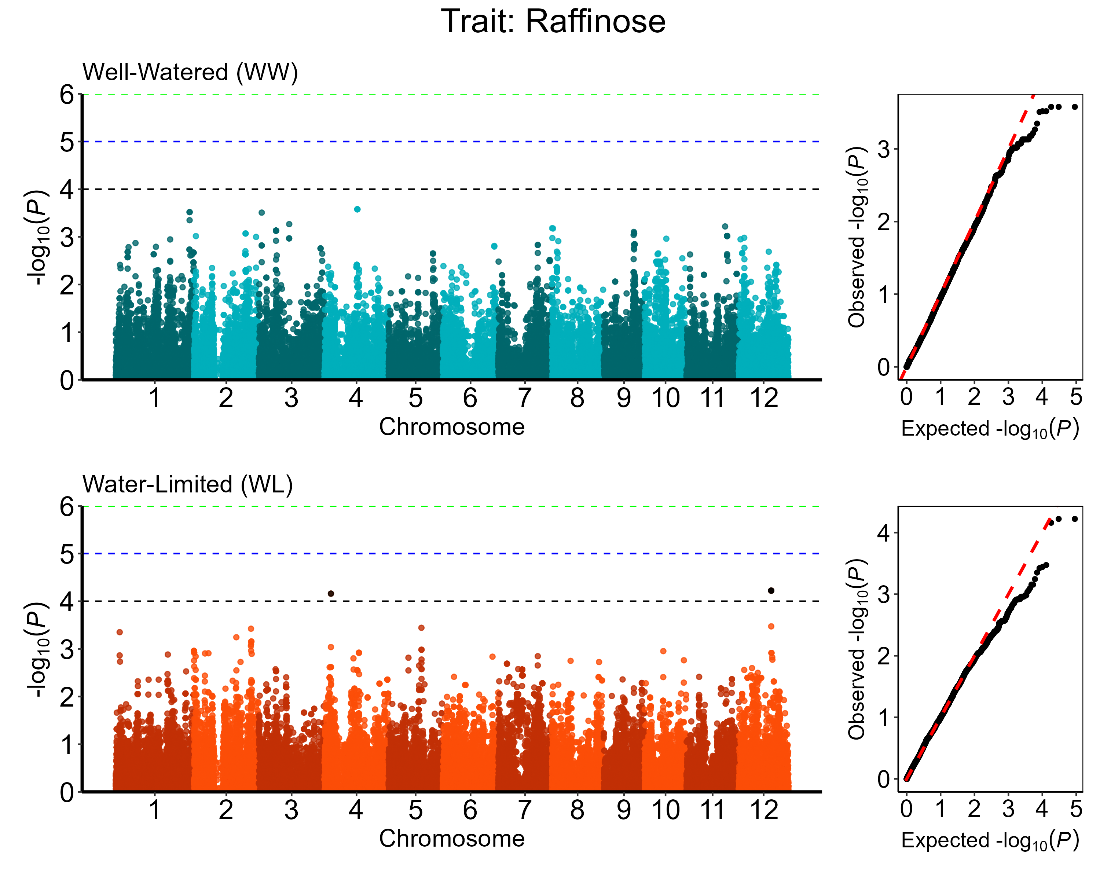

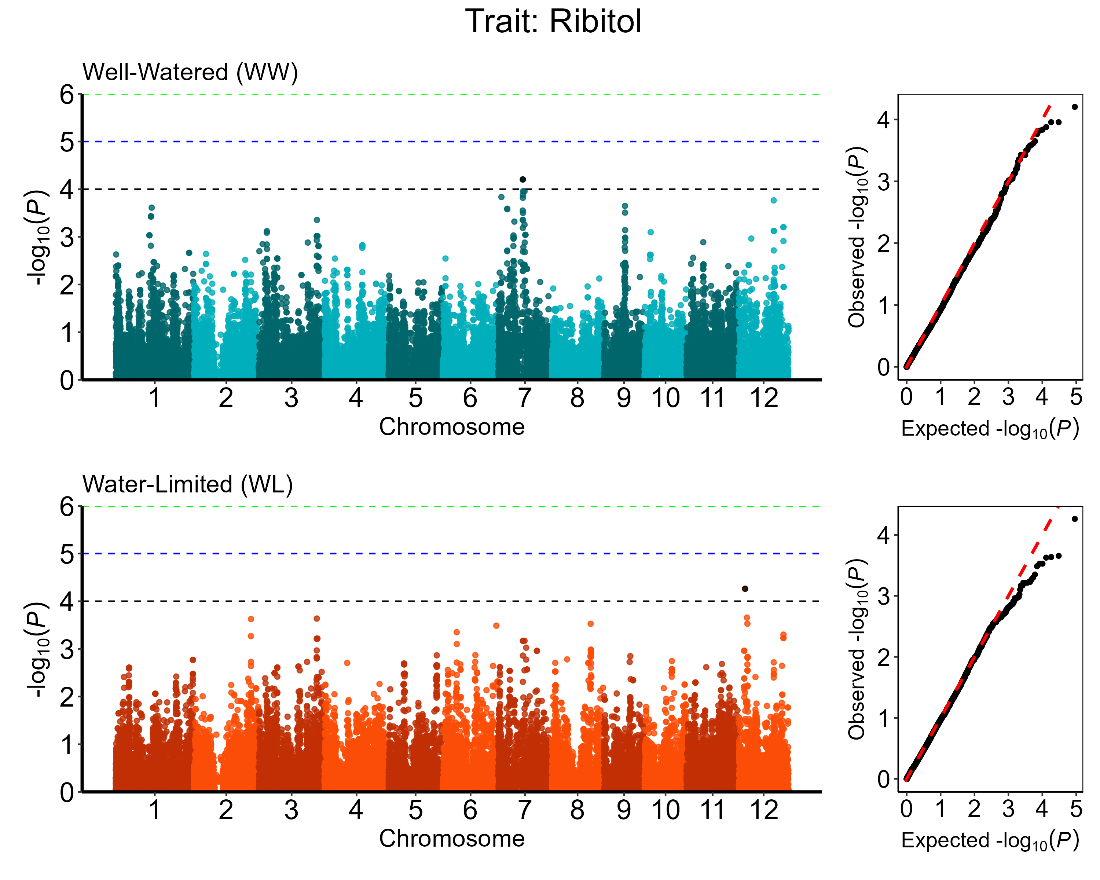

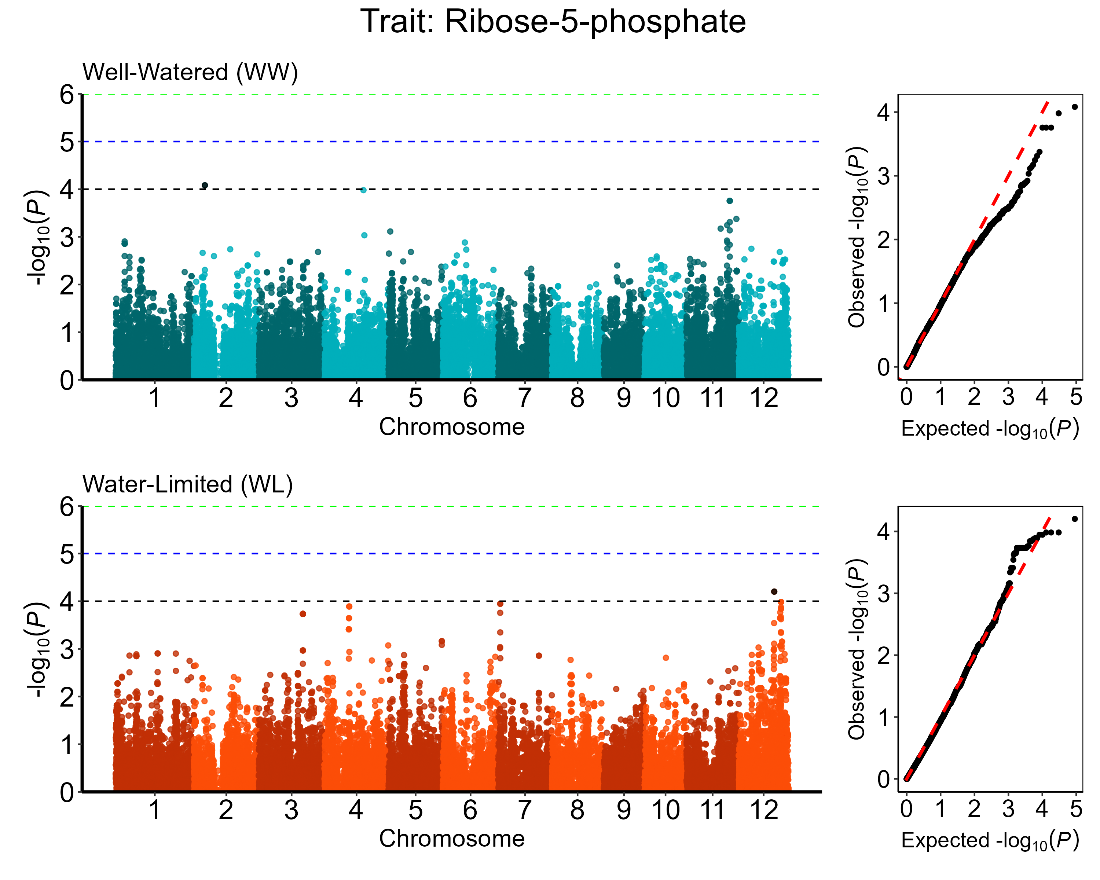

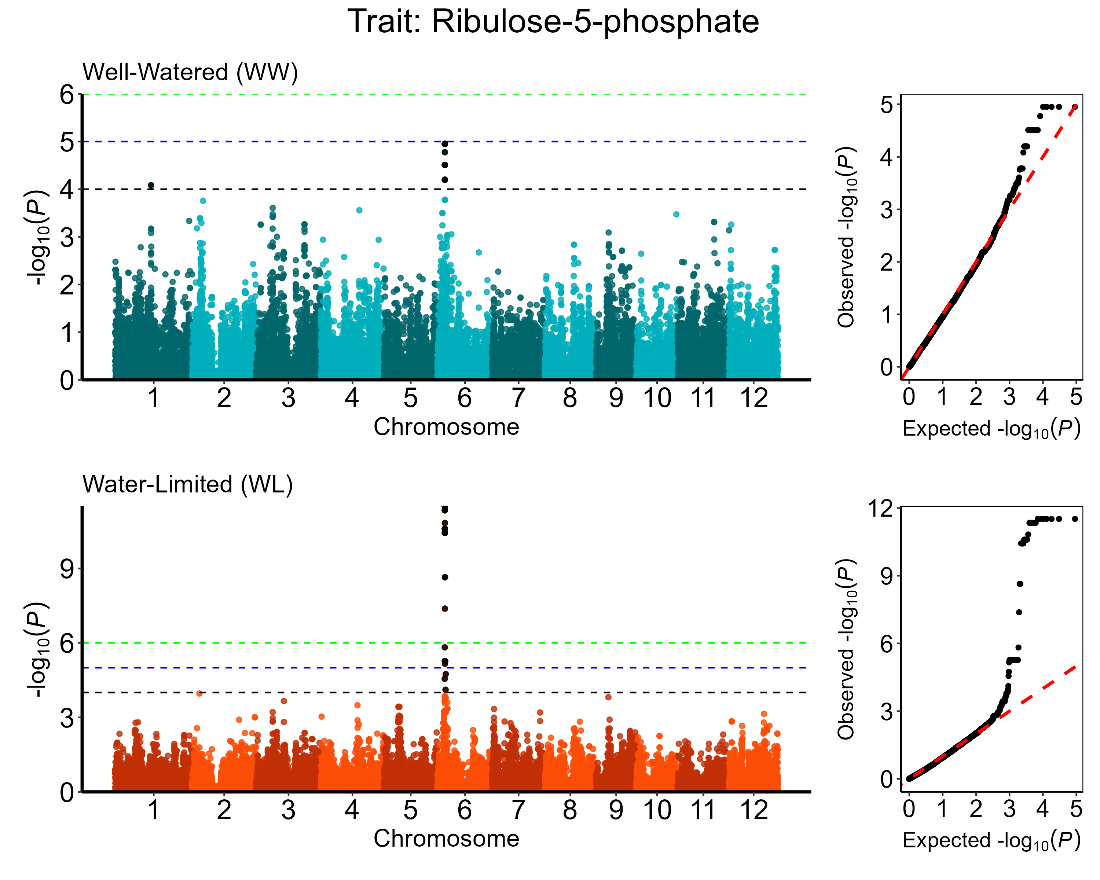

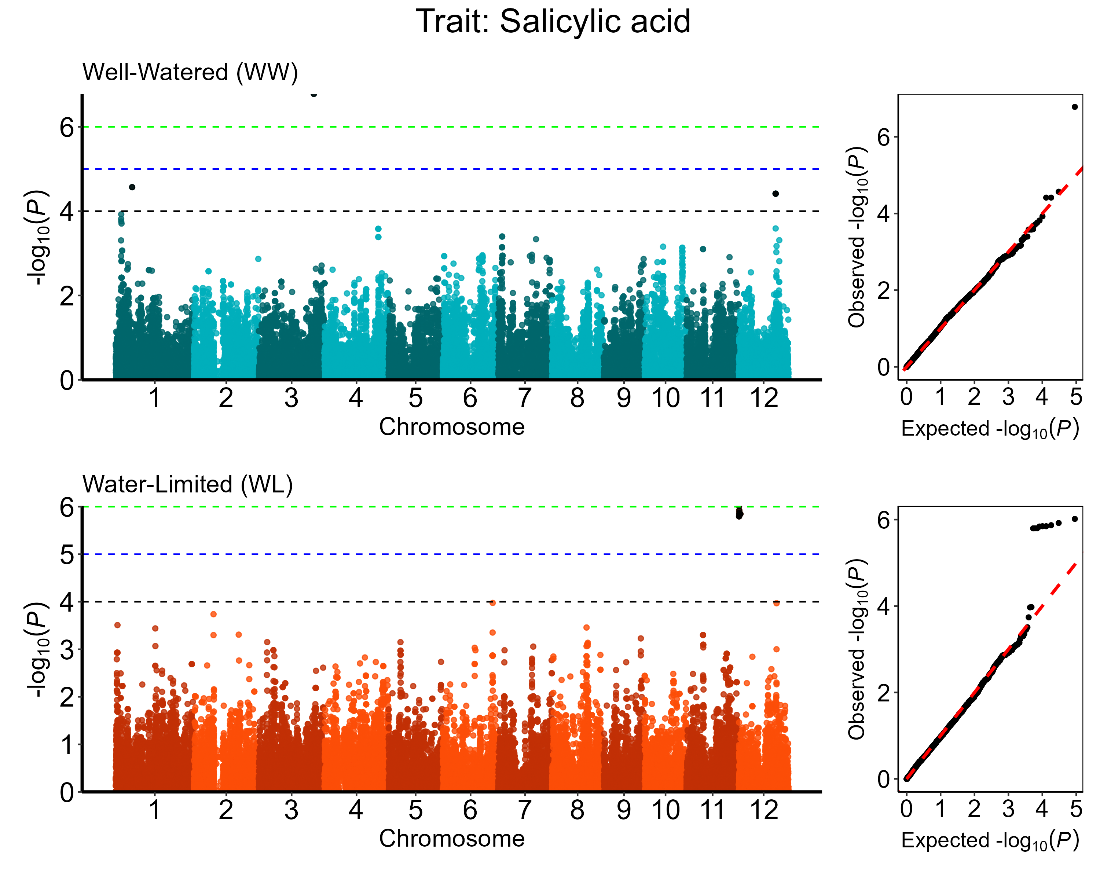

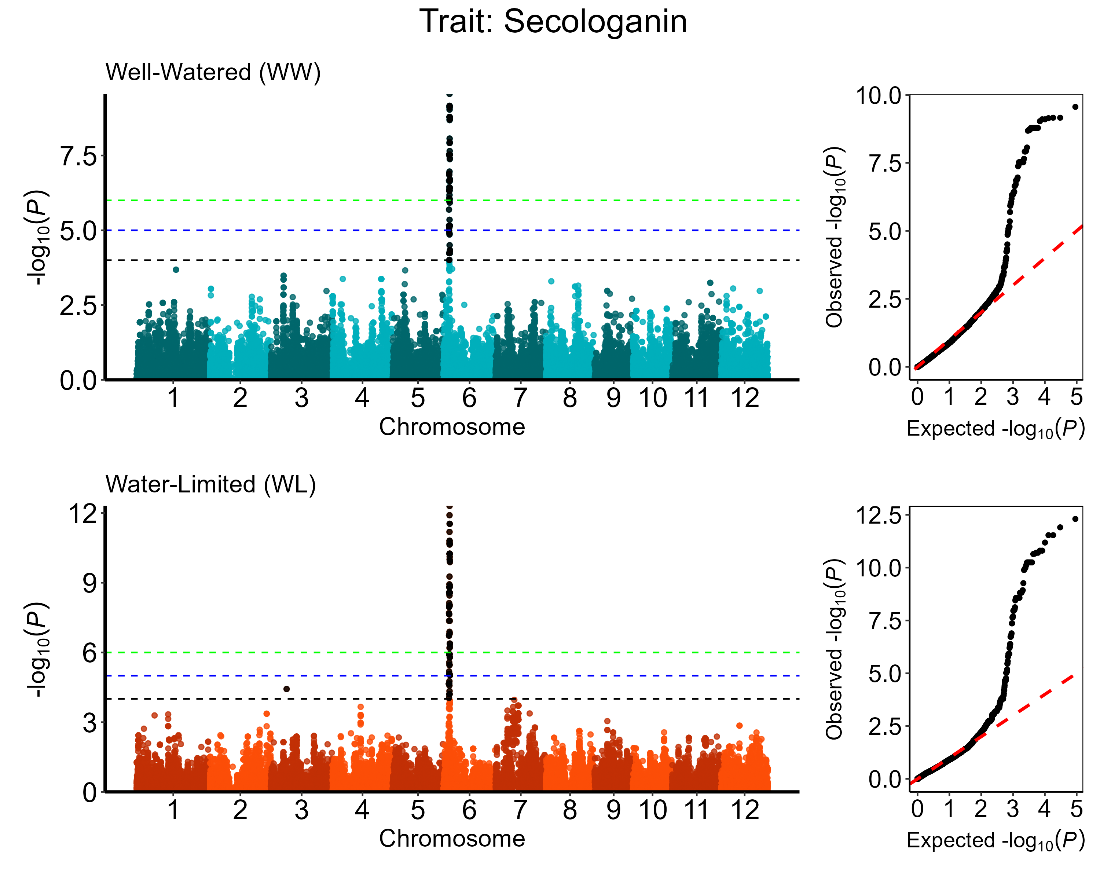

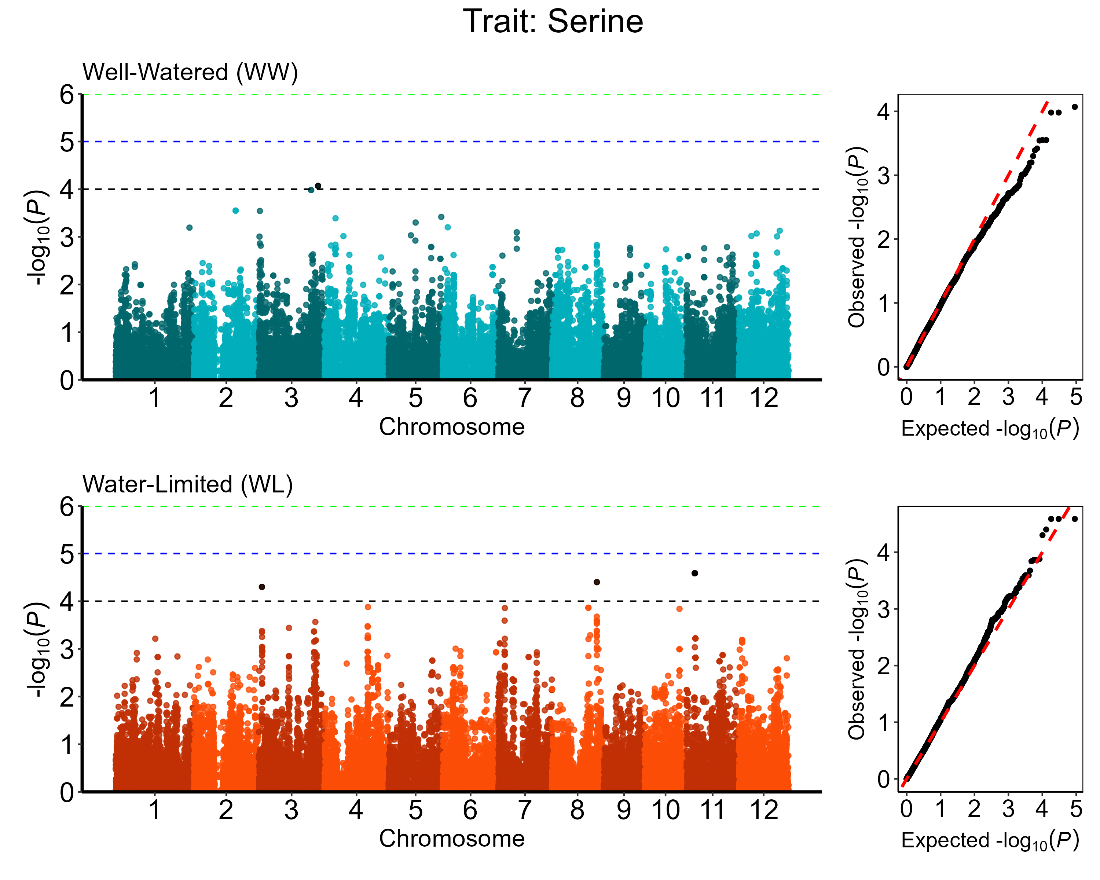

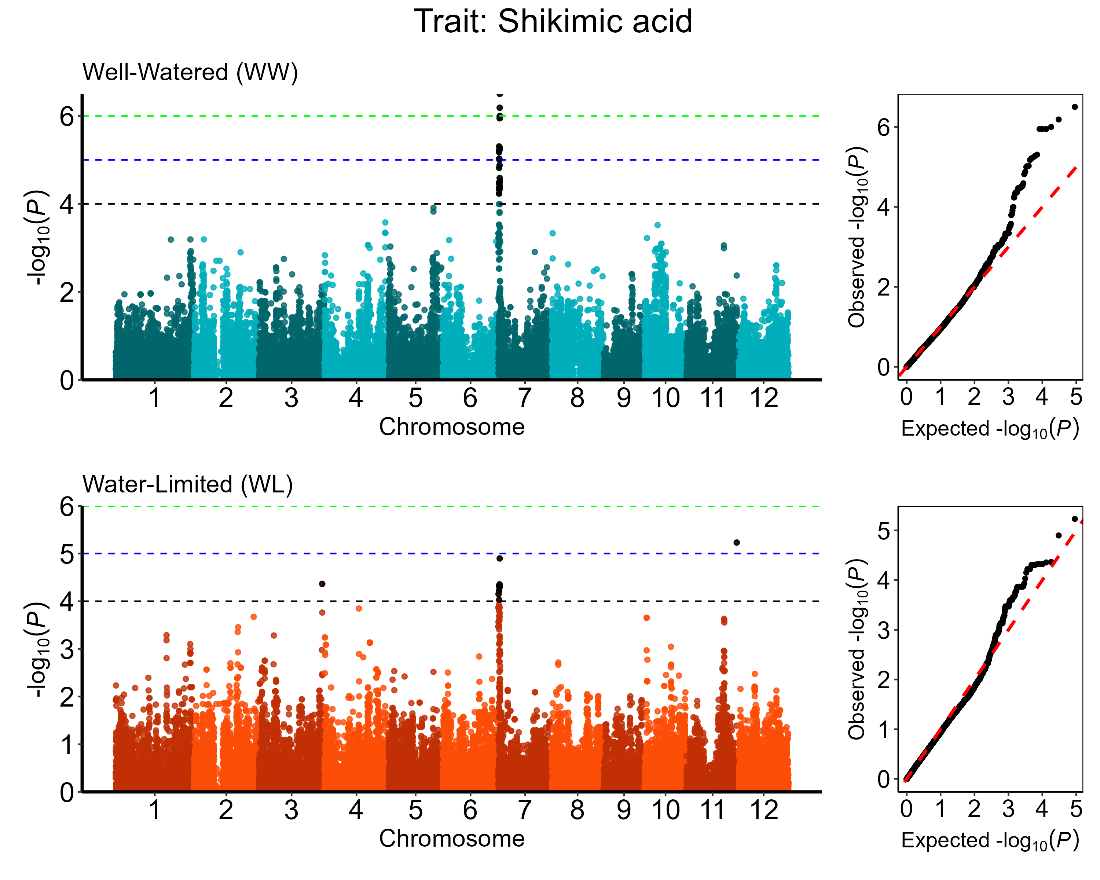

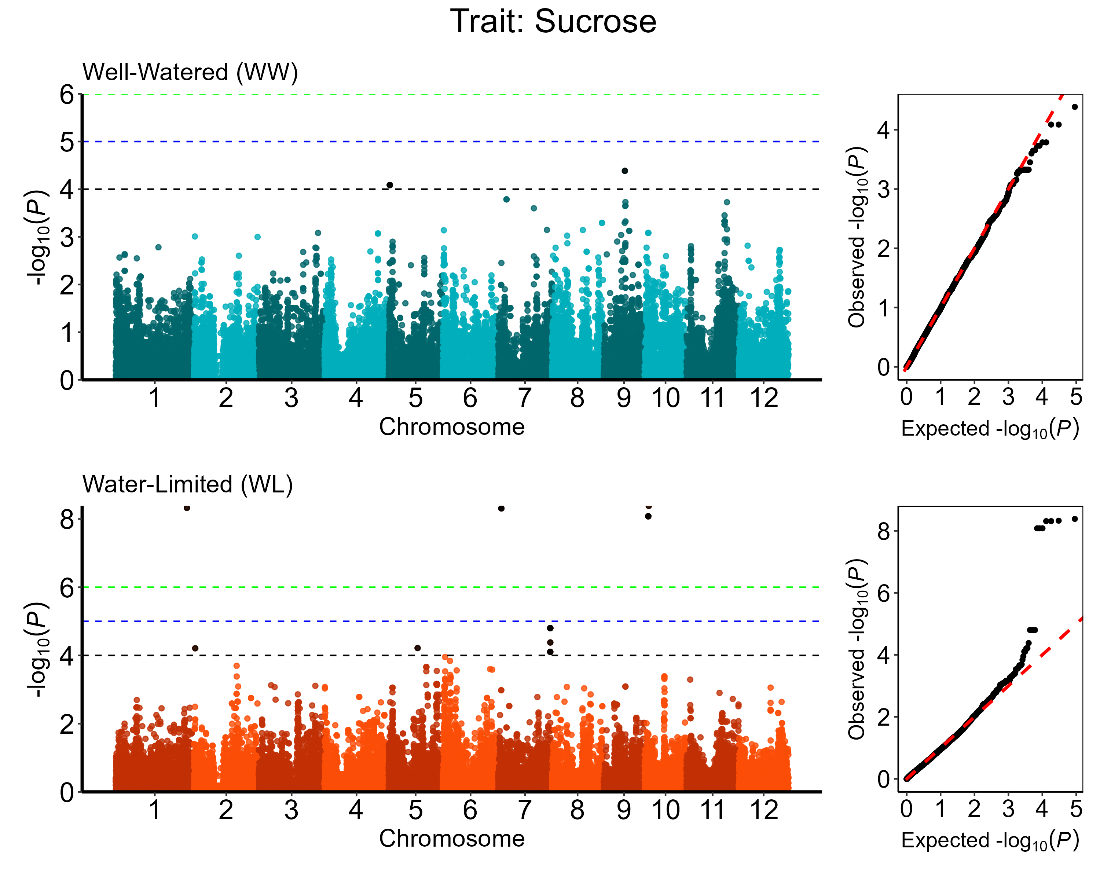

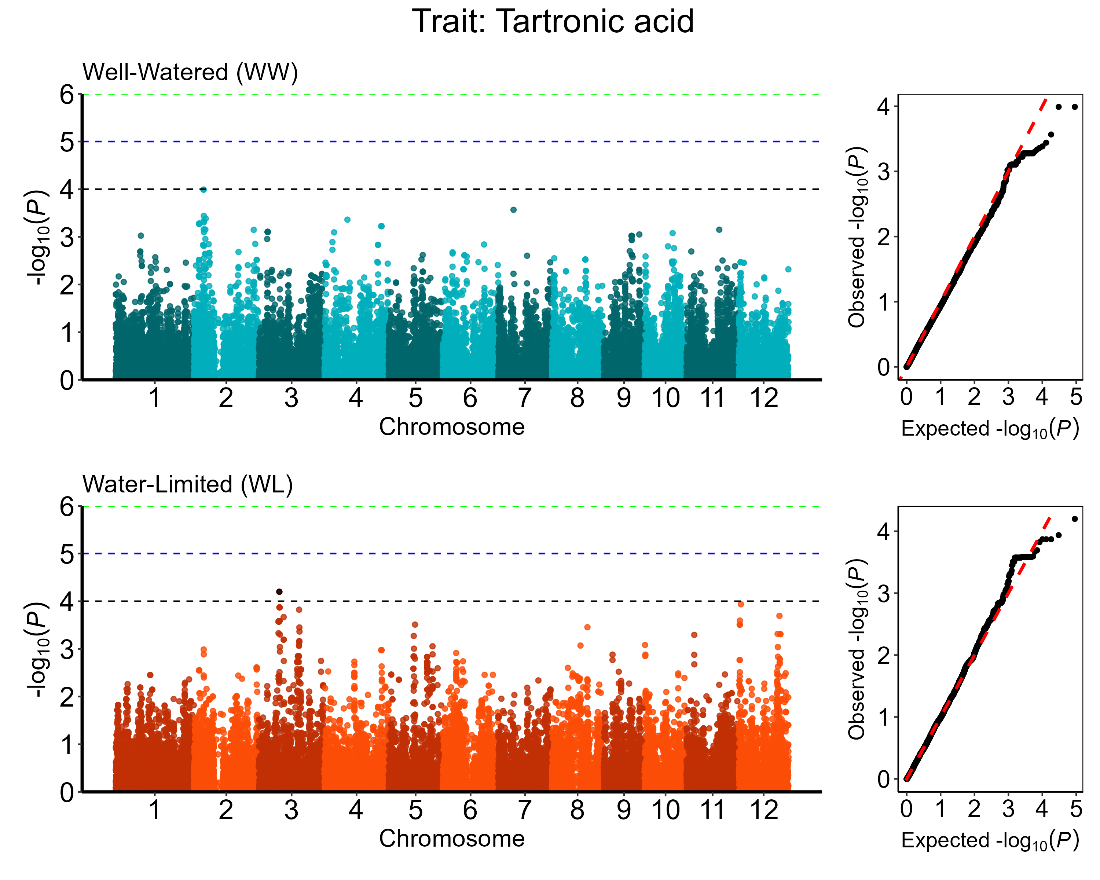

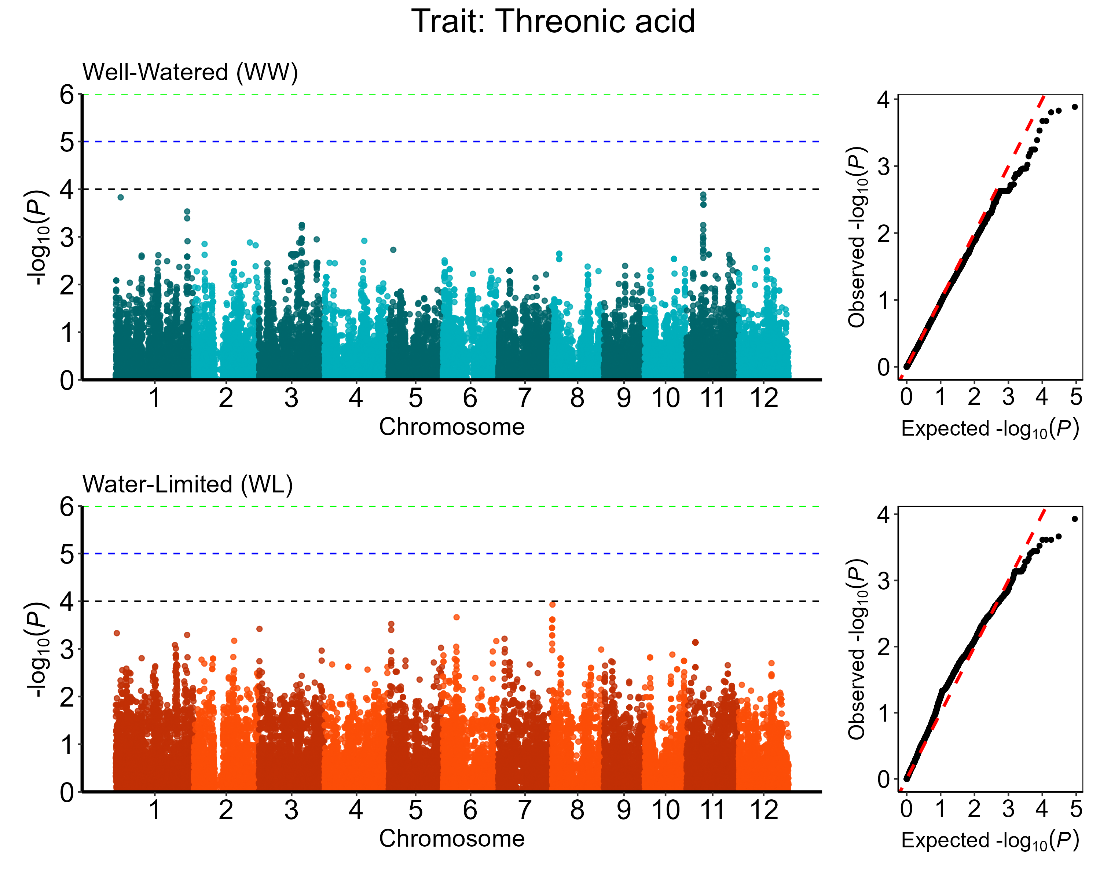

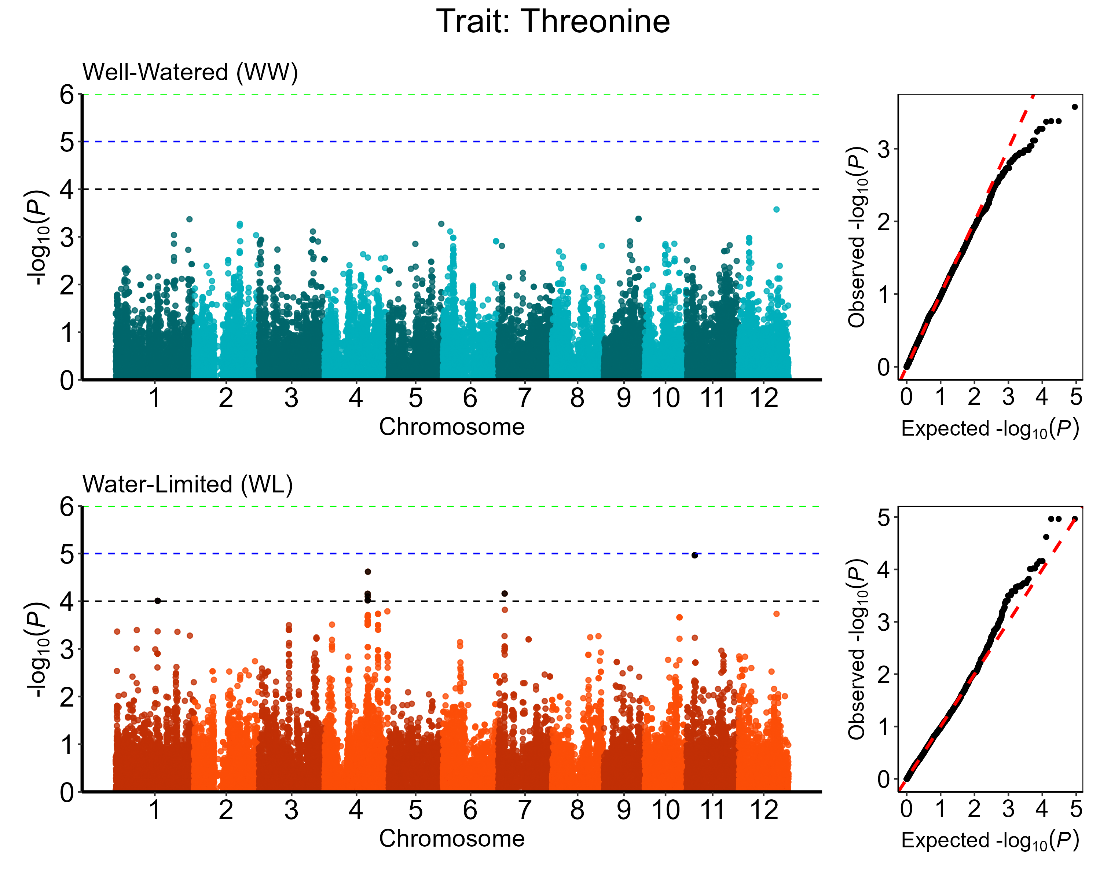

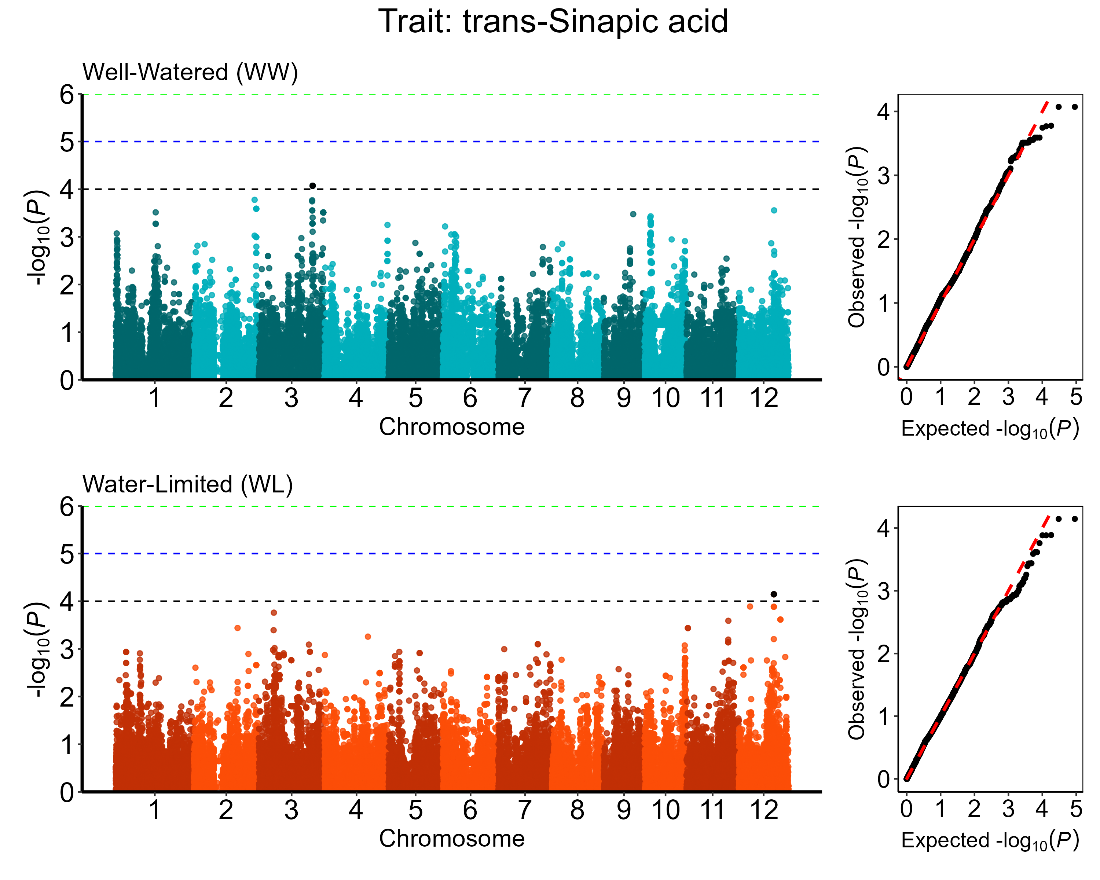

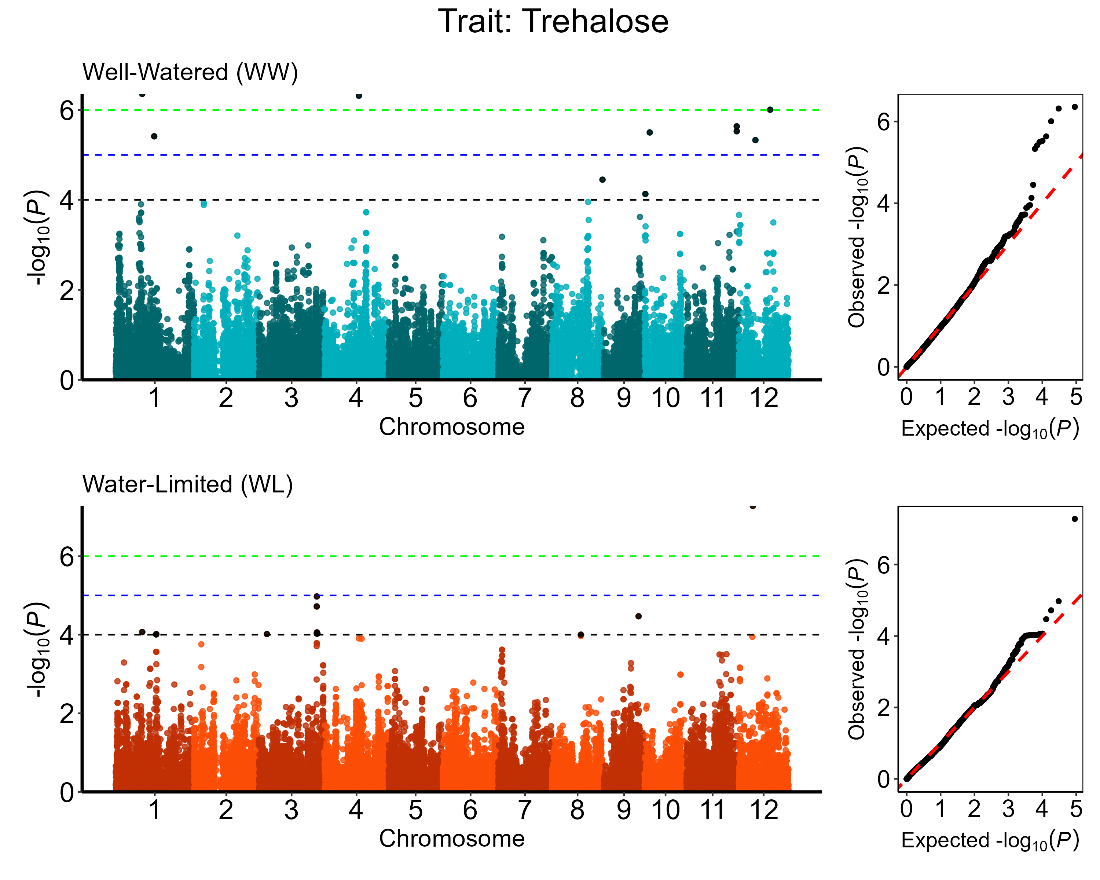

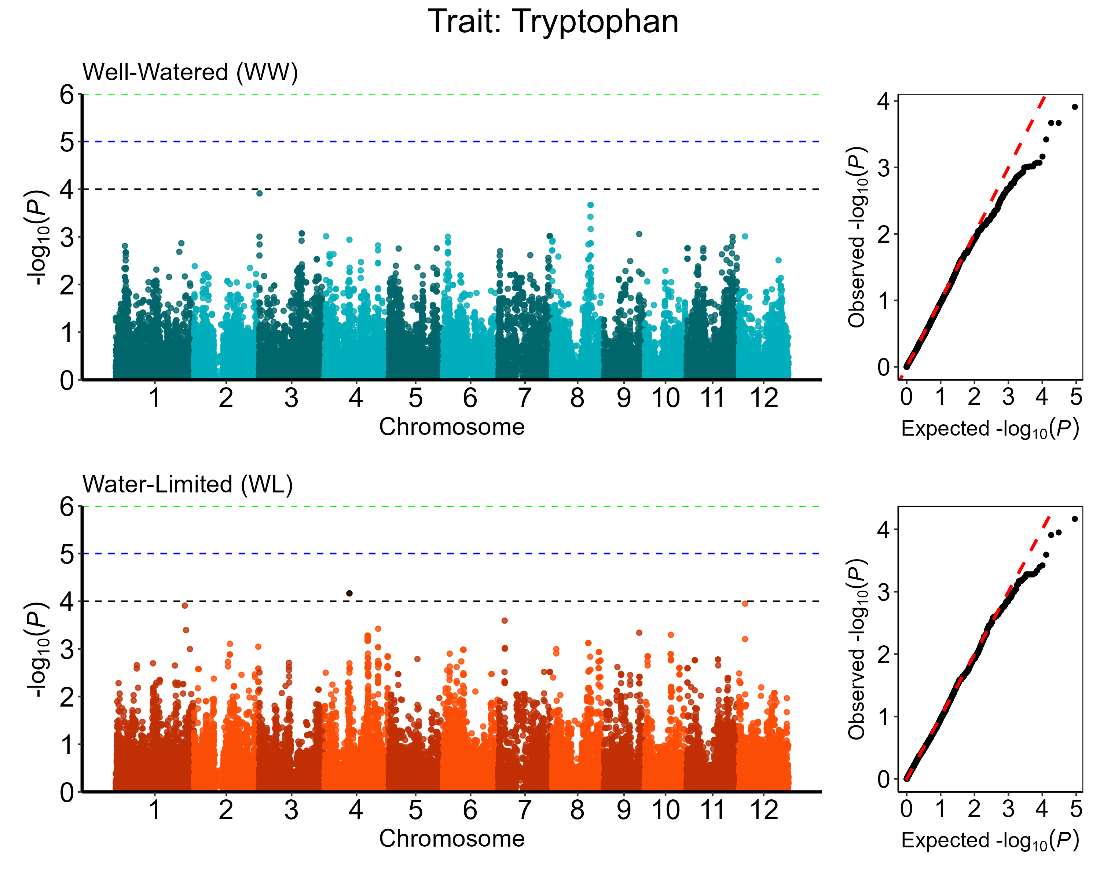

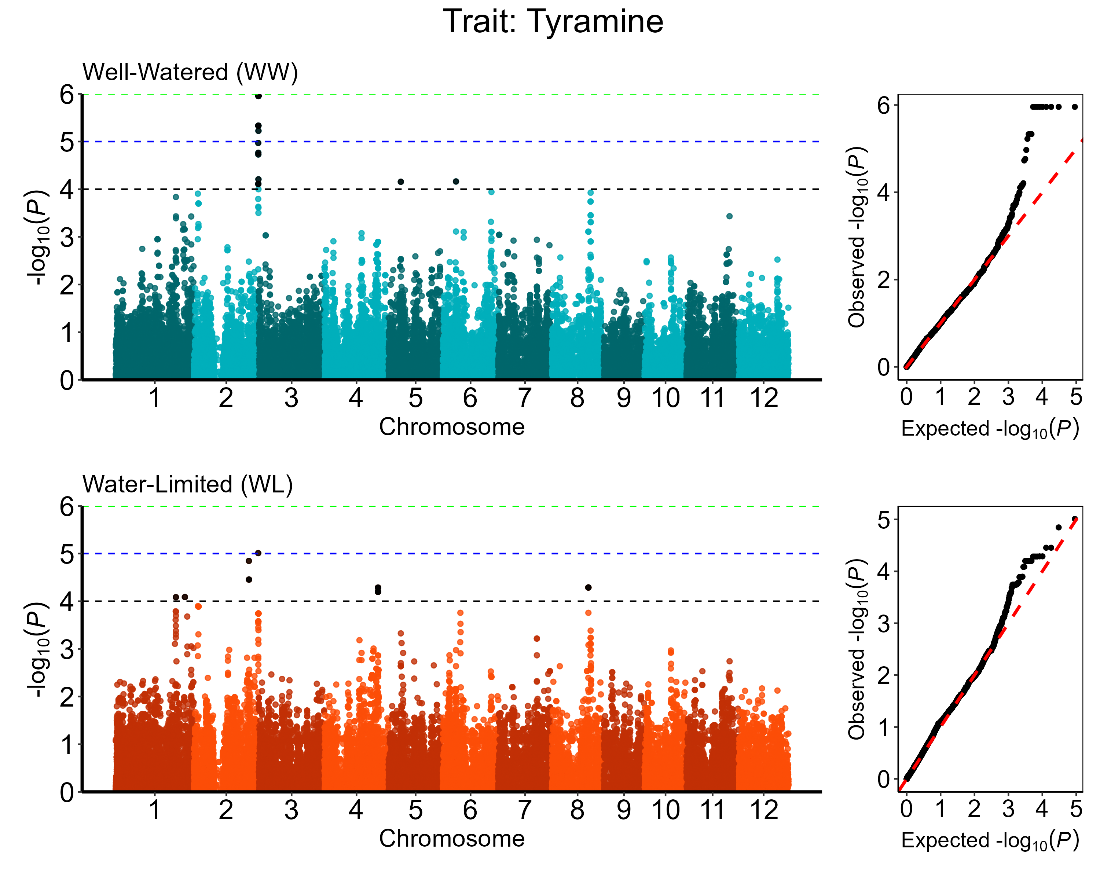

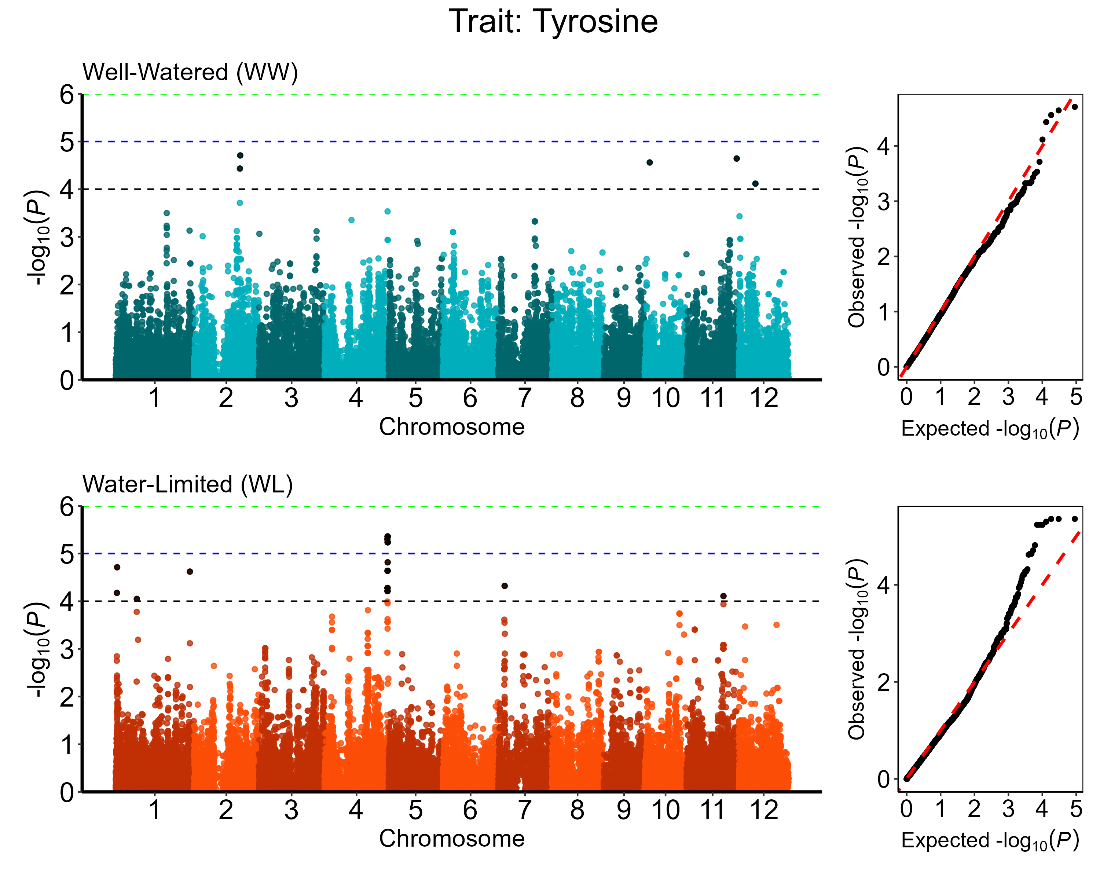

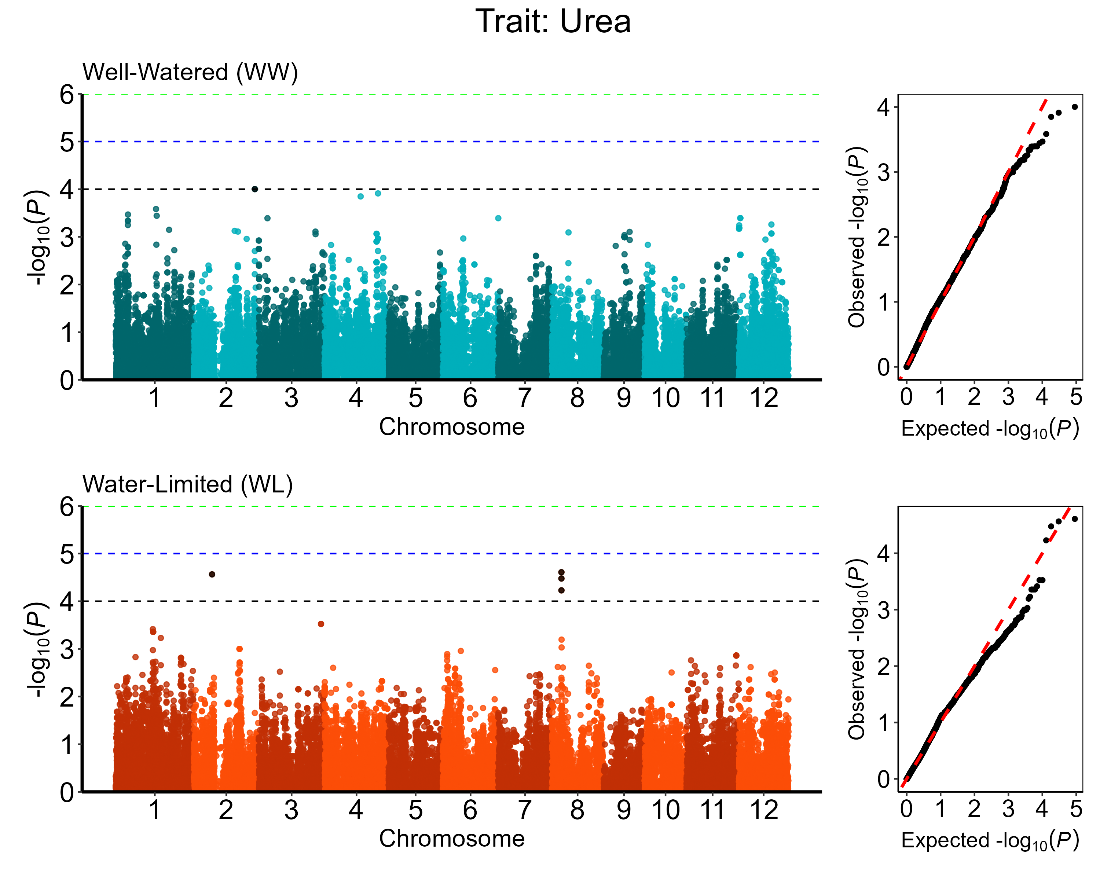

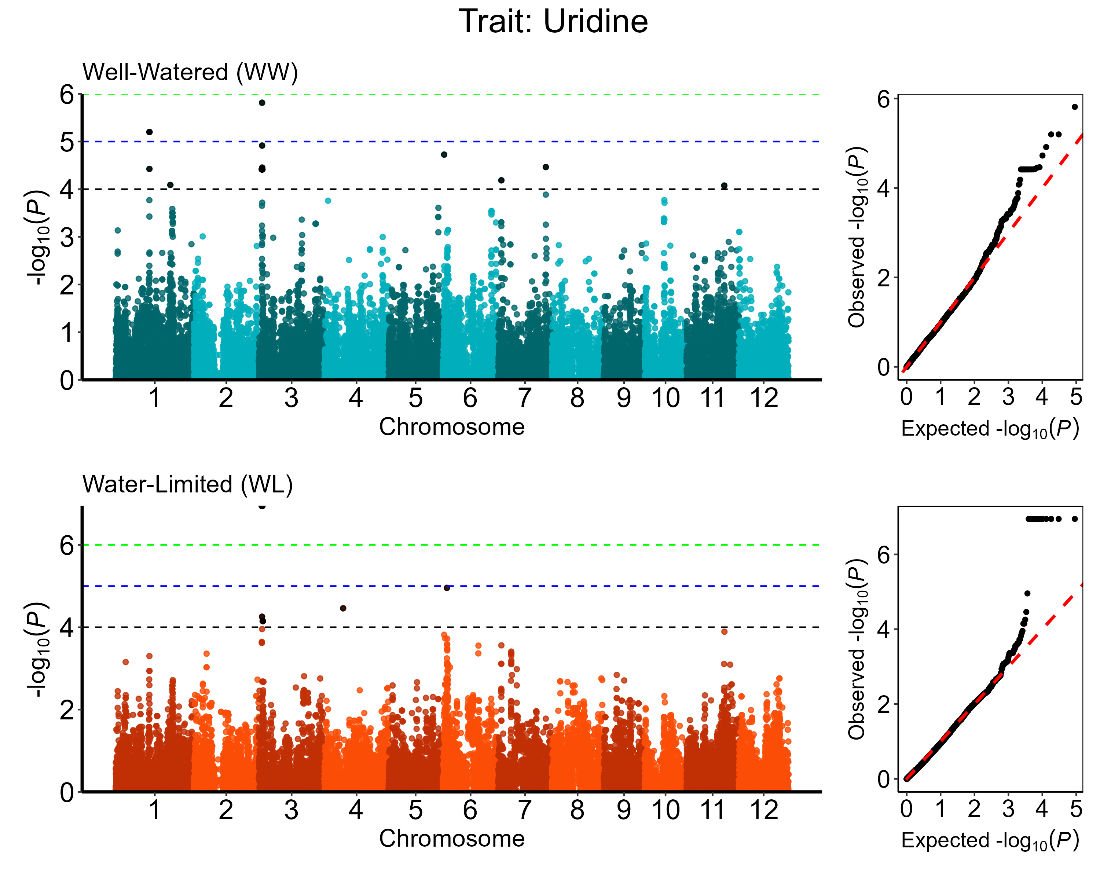

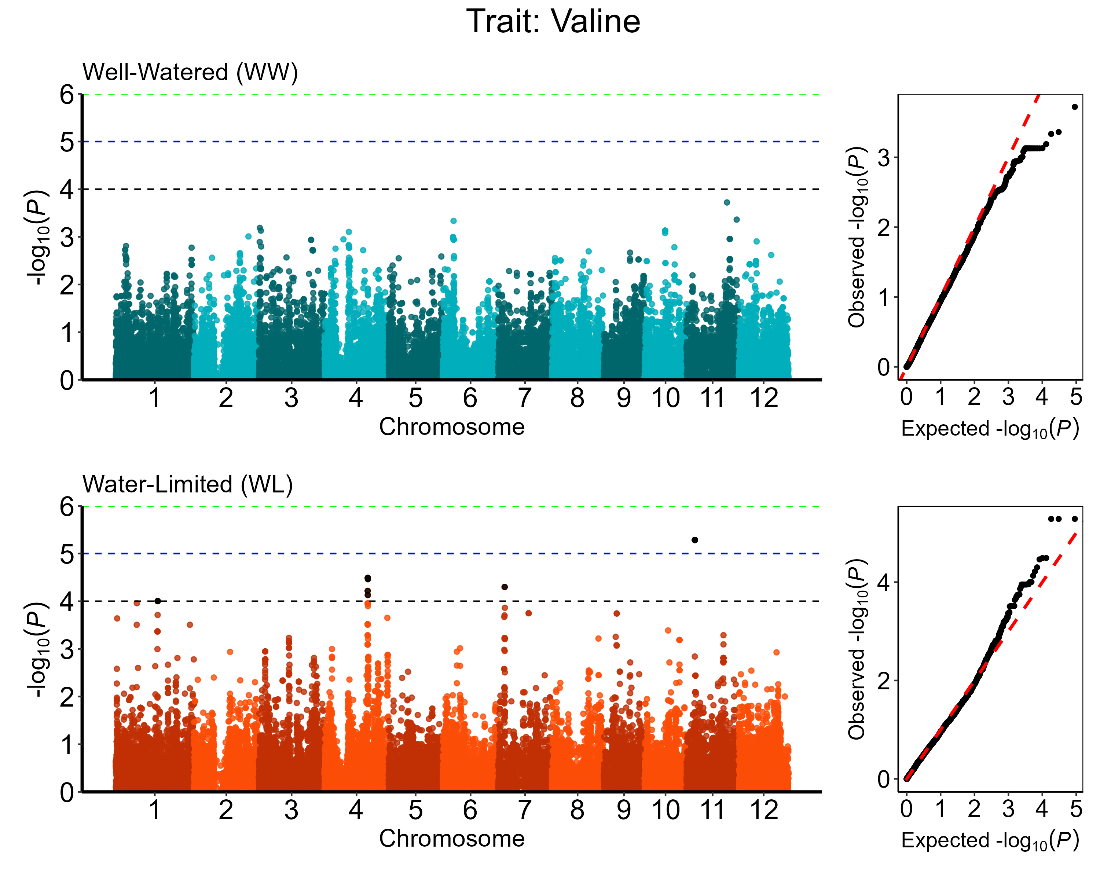

**Supplementary Fig. S11A – S99D. Manhattan and QQ plots of GWA mapping of the 89 polar primary metabolites traits under well-watered (WW) and water-limited (WL) conditions.**

Dashed horizontal lines indicate the significance thresholds used to classify association signals: low significance (-log_10_*(P)* ≥ 4; black), medium significance (-log_10_*(P)* ≥ 5; purple), and high significance (-log_10_*(P)* ≥ 5.96; green), where the high-significance threshold corresponds to the Bonferroni-corrected threshold (α = 0.05).

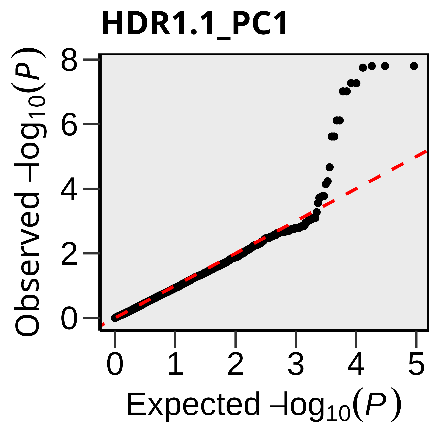

**HDR1-1_PC1**

**HDR1-1_PC1**

**HDR1-2_PC1**

**HDR1-2_PC1**

**Supplementary Fig. S100A – S104B. Manhattan and QQ plots of GWA mapping of the PC1 of all 4 HDRs (HDR1-1, HDR1-2, HDR4 and HDR11).**

Dashed horizontal line (-log_10_*(P)* ≥ 4; black) indicates the significance threshold used to classify association signals.
