## Supplementary file4 (Fig. S105 - S107) for "GWAS of flag leaf primary metabolites uncovers metabolic QTL hubs linked to grain yield stability under drought in field-grown *indica* rice"

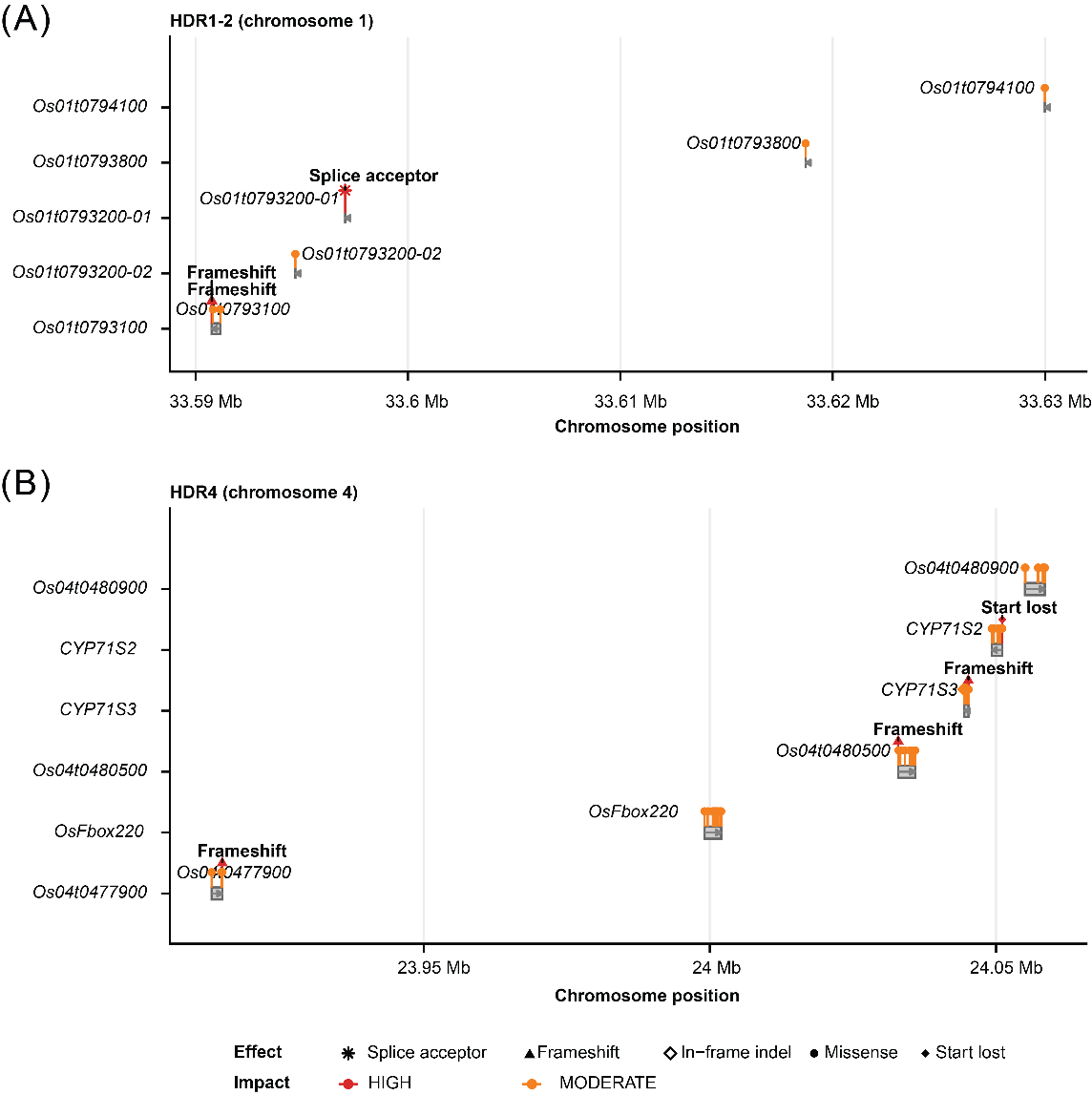

**Supplementary Fig. S105. Candidate genes and predicted functional variants within HDR1-2 and HDR4.** (A) Genomic distribution of predicted coding variants within the HDR1-2 interval on chromosome 1. Gene models are shown along the y-axis and positioned according to their physical coordinates on the x-axis. Symbols denote variant effects, including frameshift and splice acceptor mutations, while colors indicate predicted impact severity (HIGH or MODERATE). (B) Genomic distribution of predicted coding variants within the HDR4 interval on chromosome 4. Variant effects include frameshift and start-loss mutations, with colors representing predicted impact severity.

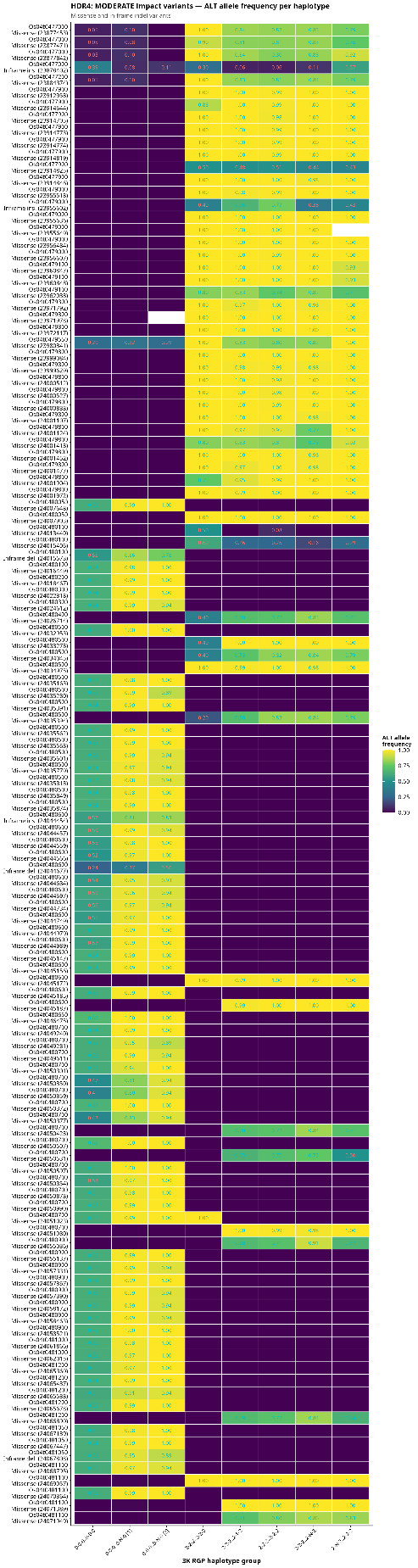
(A)

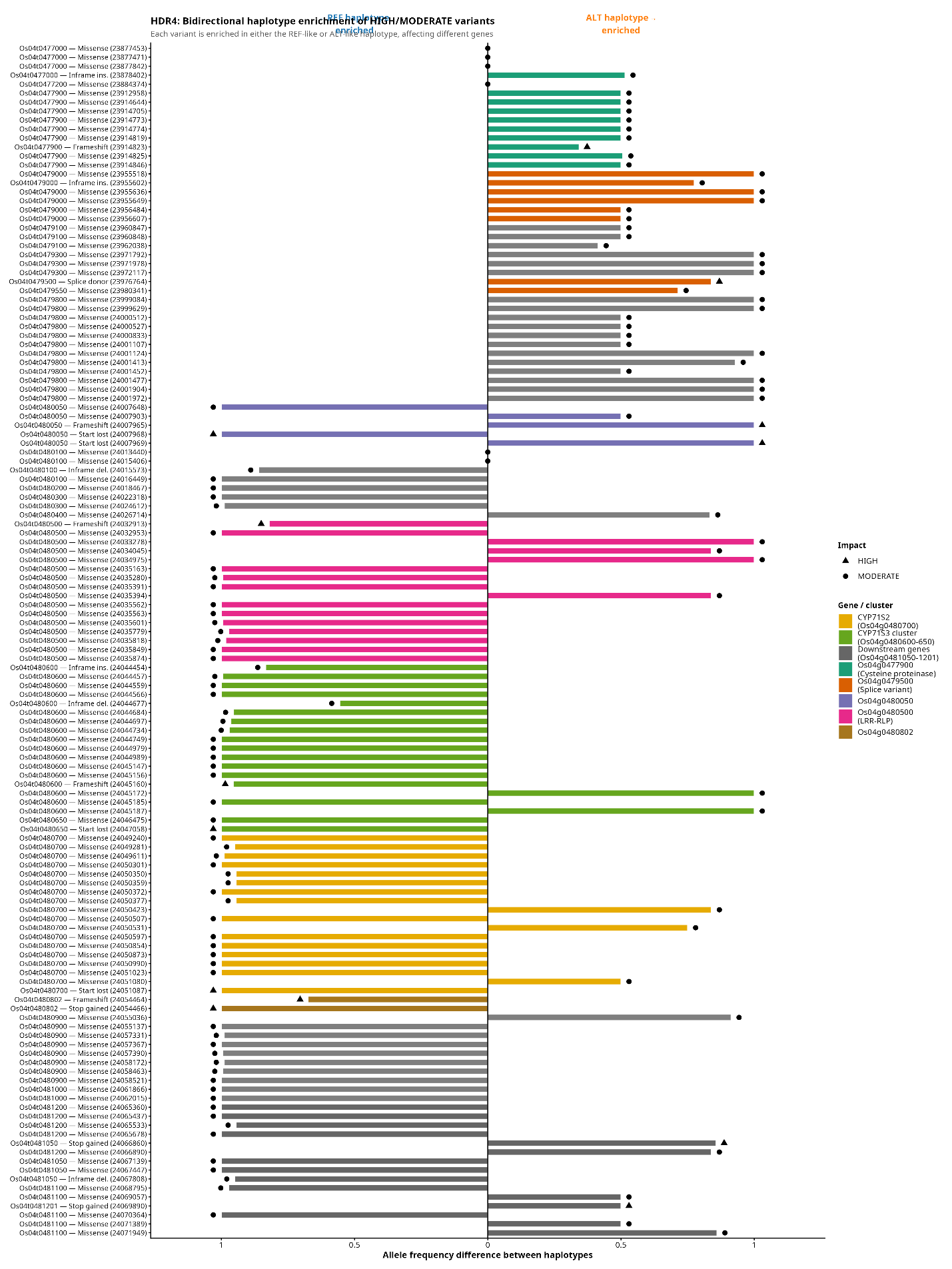
**Supplementary Fig. S106. Distribution of predicted variants on candidate genes within HDR4.** (A) Heatmap of predicted variants of MODERATE class within HDR4. Boxes are colored based on ALT allele frequency per haplotype. (B) Bidirectional haplotype enrichment of HIGH/MODERATE variants in HDR4. Lines are colored by gene/cluster and shapes (circle or triangle) at the end of the lines represent predicted impact class.

(B)

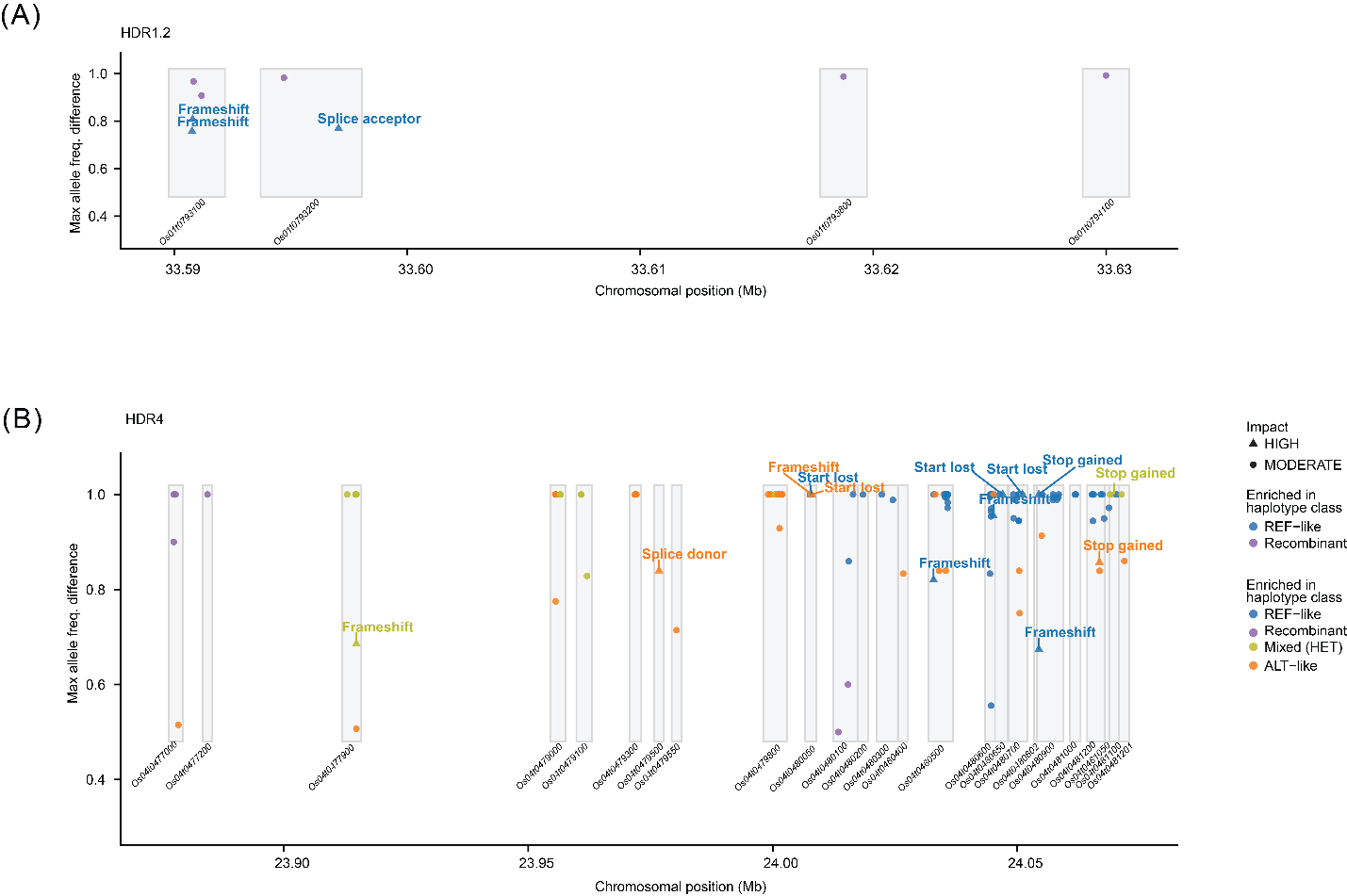

**Supplementary Fig. S107. Allele-frequency differentiation and functional variants within HDR1-2 and HDR4.** (A) Maximum allele-frequency differences between HDR1-2 haplotype groups for variants located within genes across the HDR1-2 interval on chromosome 1. Gray boxes denote gene boundaries, and points represent variants colored according to the haplotype class in which the alternative allele is enriched. Functional variants predicted to have high-impact effects, including frameshift and splice-acceptor mutations, are annotated. (B) Maximum allele-frequency differences between HDR4 haplotype groups for variants located within genes across the HDR4 interval on chromosome 4. Gray boxes indicate gene boundaries, while points represent individual variants colored by the haplotype class in which the alternative allele is enriched (REF-like, recombinant, mixed/heterozygous, or ALT-like). Symbols indicate predicted impact severity (HIGH or MODERATE), and selected high-impact variants, including frameshift, start-loss, stop-gained, and splice-donor mutations, are annotated.
